# Comprehensive characterization of N- and O- glycosylation of SARS-CoV-2 human receptor angiotensin converting enzyme 2

**DOI:** 10.1101/2020.05.01.071688

**Authors:** Asif Shajahan, Stephanie Archer-Hartmann, Nitin T. Supekar, Anne S. Gleinich, Christian Heiss, Parastoo Azadi

## Abstract

The emergence of the COVID-19 pandemic caused by SARS-CoV-2 has created the need for development of new therapeutic strategies. Understanding the mode of viral attachment, entry and replication has become a key aspect of such interventions. The coronavirus surface features a trimeric spike (S) protein that is essential for viral attachment, entry and membrane fusion. The S protein of SARS-CoV-2 binds to human angiotensin converting enzyme 2 (hACE2) for entry. Herein, we describe glycomic and glycoproteomic analysis of hACE2 expressed in HEK293 human cells. We observed high glycan occupancy (73.2 to 100%) at all seven possible N-glycosylation sites and surprisingly detected one novel O-glycosylation site. To deduce the detailed structure of glycan epitopes on hACE2 that may be involved in viral binding, we have characterized the terminal sialic acid linkages, the presence of bisecting GlcNAc, and the pattern of N-glycan fucosylation. We have conducted extensive manual interpretation of each glycopeptide and glycan spectrum, in addition to using bioinformatics tools to validate the hACE2 glycosylation. Our elucidation of the site-specific glycosylation and its terminal orientations on the hACE2 receptor, along with the modeling of hACE2 glycosylation sites can aid in understanding the intriguing virus-receptor interactions and assist in the development of novel therapeutics to prevent viral entry. The relevance of studying the role of ACE2 is further increased due to some recent reports about the varying ACE2 dependent complications with regard to age, sex, race, and pre-existing conditions of COVID-19 patients.

## Introduction

In late 2019, the emergence of the highly transmissible coronavirus disease (COVID-19) led to a global health crisis within weeks and was soon declared a pandemic. The new underlying pathogen belongs to the family of the *coronaviridae* and has been named severe acute respiratory syndrome coronavirus 2 (SARS-CoV-2), initially termed 2019 novel coronavirus (2019-nCoV) (Gorbalenya, A.E., Baker, S.C., et al. 2020). More than six months into the pandemic, no promising vaccine candidates or treatments are currently available for COVID-19 (Li, G.D. and De Clercq, E. 2020, World Health Organization (WHO) 2020). The elucidation of key structures involved in the transmission of SARS-CoV-2 will provide insights towards design and development of suitable vaccines and drugs against COVID-19 to curb the global public health crisis (Wang, N., Shang, J., et al. 2020).

Located on the viral surface is the spike (S) protein, which attaches the SARS-CoV-2 pathogen to target cells in the human body. The trimeric spike protein belongs to the class I fusion proteins. Its two subunits S1 and S2 orchestrate its entry into the cell: The S1 subunit facilitates the attachment of the virus via its receptor binding domain (RBD) to the host cell receptor, while the S2 subunit mediates the fusion of the viral and human cellular membranes (Hoffmann, M., Kleine-Weber, H., et al. 2020, Shang, J., Ye, G., et al. 2020, Walls, A.C., Park, Y.J., et al. 2020). The glycosylation pattern of the spike protein, which carries N-linked, as well as O-linked glycosylation, has been the subject of several recent scientific studies (Shajahan, A., Supekar, N.T., et al. 2020b, Watanabe, Y., Allen, J.D., et al. 2020, Zhang, Y., Zhao, W., et al. 2020). Insight into the glycosylation pattern expands the understanding of the viral binding to receptors, the fusion event, host cell entry, replication, as well as the design of suitable antigens for vaccine development. Our recent study on the site-specific quantitation of N-linked and O-linked glycosylation of the subunits S1 and S2 of the SARS-CoV-2 spike protein revealed the presence of an O-glycosylation site at the RBD of subunit S1 (Shajahan, A., Supekar, N.T., et al. 2020b), although Watanabe *et al*. (9) reported lower abundances of O-linked glycans on the full-length spike protein. A number of amino acids within the RBD have been shown to be crucial determinants for binding in general and the binding affinity in particular of the virus to the host cell receptors (Andersen, K.G., Rambaut, A., et al. 2020, Wan, Y., Shang, J., et al. 2020). Several studies have identified the human angiotensin-converting enzyme 2 (hACE2, ACEH) as the key virus receptor of SARS-CoV-1 in the early 2000s (Kuba, K., Imai, Y., et al. 2005, Li, W., Moore, M.J., et al. 2003). Moreover, hACE2 was identified recently as the receptor for cell entry of SARS-CoV-2 (Hoffmann, M., Kleine-Weber, H., et al. 2020, Zhou, P., Yang, X.L., et al. 2020). However, the SARS-CoV-1 and SARS-CoV-2 pathogens exploit the hACE2 receptor for cell entry only, which is unrelated to its physiological function (Li, F. 2013).

hACE2 is a type-I transmembrane protein and includes an extracellular, a transmembrane, and a cytosolic domain within a total of 805 amino acids (Jiang, F., Yang, J., et al. 2014). Interestingly, it has been found that the transmembrane form can be cleaved to a soluble form of hACE2 (sACE2) that lacks the transmembrane and cytosolic domains but is enzymatically active, since the catalytic site, as well as the zinc-binding motif lie within the extracellular region (Tipnis, S.R., Hooper, N.M., et al. 2000). hACE2 is secreted by endothelial cells and is involved in the renin-angiotensin system (RAS) (Donoghue, M., Hsieh, F., et al. 2000).

The presence of N-glycosylation was experimentally verified by treatment with the endoglycosidase PNGase F and monitoring the shift in the gel migration of hACE2 to the predicted molecular mass of ~85 kDa. (Li, W., Moore, M.J., et al. 2003, Tipnis, S.R., Hooper, N.M., et al. 2000). An earlier analysis on hACE2 N-linked glycans via sequential exoglycosidase digestion and HPLC after 2-aminobenzamide labeling identified mainly biantennary N-linked glycans with sialylation and core fucosylation, along with sialylated tri- and tetra-antennary N-glycans. A preprint and a patent have described glycoproteomics of hACE2 on non-human cells. However, glycosylation on such systems is not expected to reflect what is found in human ACE2 (Schuster, M., Loibner, H., et al. 2013, Sun, Z., Ren, K., et al. 2020). Nevertheless, we could not find any reports with detailed structure elucidation of hACE2 glycans, even on multiplatform glycosylation resource glygen.org (Chen, R., Jiang, X., et al. 2009, Kristiansen, T.Z., Bunkenborg, J., et al. 2004, Towler, P., Staker, B., et al. 2004, York, W.S., Mazumder, R., et al. 2020). The study of glycosylation can help in understanding the key roles glycans play during the physiological function of hACE2 and more importantly, its involvement in facilitating viral binding.

Within this study, we report the quantitative site-specific N-linked and O-linked glycan characterization of hACE2 by a glycomic and glycoproteomic approach. We identified glycosylation at all seven potential N-glycosylation sites on hACE2 and also report one O-glycosylation site. The N- and O-glycans were released from hACE2 and structurally characterized by MALDI-MS and ESI-MS^n^. Particular focus was placed upon the location of fucosylation, the presence of bisecting GlcNAc residues, and the linkages of sialic acids.

## Results

### Implication of glycosylation on hACE2 binding with SARS-CoV-1 and SARS-CoV-2 S protein

We examined the co-crystallization structures of SARS-CoV-1 S protein trimer with hACE2 (PDB ID - 6ACG) and SARS-CoV-2 RBD with hACE2 dimer (PDB ID - 6M17) to understand the binding location of S protein with hACE2 and the proximity of glycosylation sites (Song, W., Gui, M., et al. 2018, Yan, R., Zhang, Y., et al. 2020). Recently, it was reported that a dimeric ACE2 can accommodate two S protein trimers, each through a monomer of ACE2 (Yan, R., Zhang, Y., et al. 2020). The binding location of individual monomers of the S protein with each monomer of the hACE2 dimer is shown in Figure 1. The image indicates that the N-glycosylation and its epitopes, particularly at sites N90 and N322, could play a critical role in the binding of hACE2 to the RBD of S protein. Our recent study on the glycosylation of individual S protein subunits (Shajahan, A., Supekar, N.T., et al. 2020b) showed the prevalence of high mannose type glycans at the RBD of SARS-CoV-2 spike protein (N331 and N343), while the results of Watanabe *et al*. on trimeric S protein (Watanabe, Y., Allen, J.D., et al. 2020) indicated a predominance of complex glycans at these sites. The relevance of the different types of glycans and their implication on the receptor binding needs further investigation.

**Figure 1:**
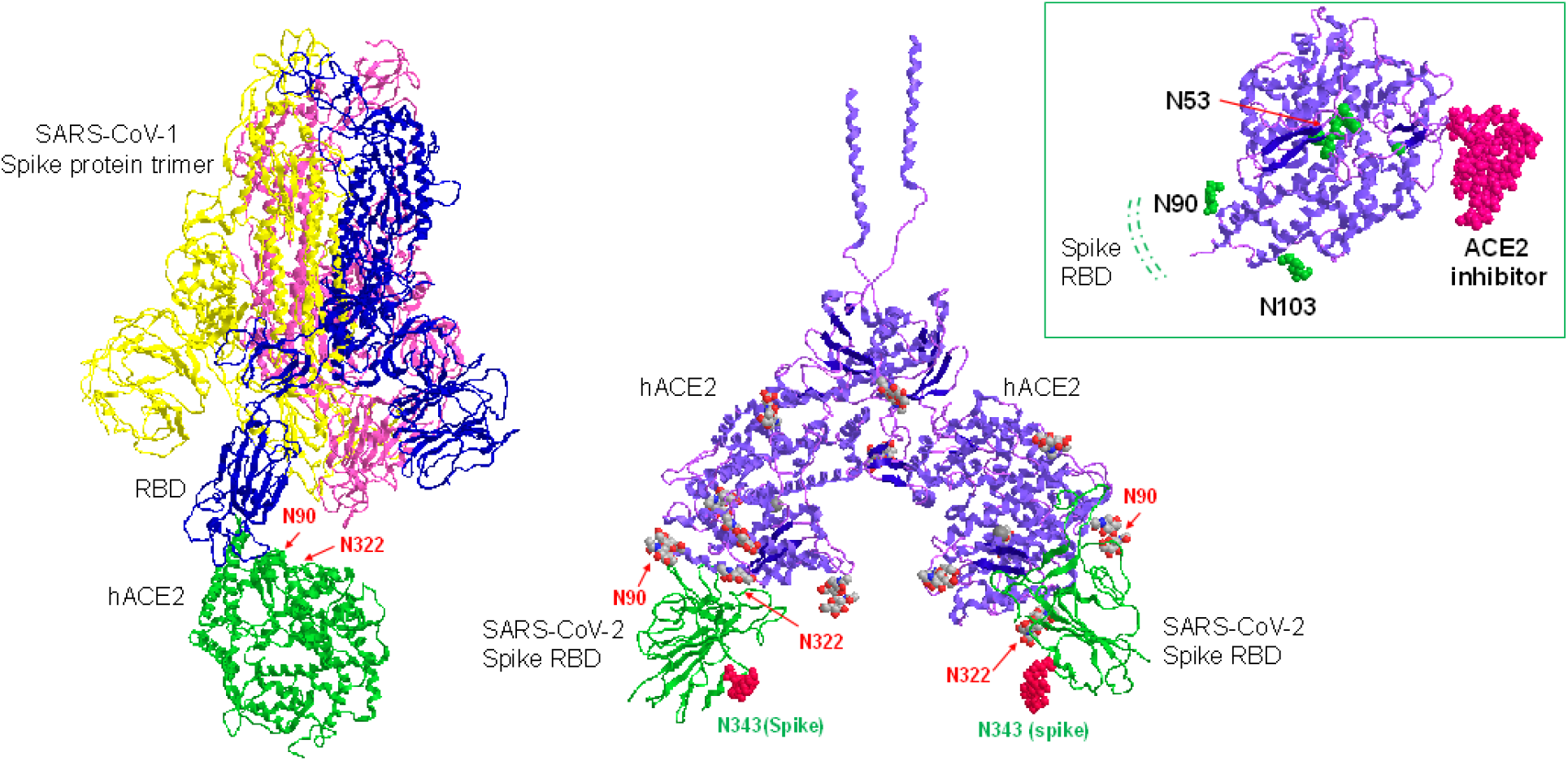
Co-crystallization structure of SARS-CoV-1 trimer with hACE2 (PDB ID – 6ACG) and SARS-CoV-2 RBD with hACE2 (PDB ID - 6M17) shows the binding of individual monomer of spike (S) protein with peptidase domain (PD) of each monomer of hACE2 dimer. Inset – binding of hACE2 with inhibitor (PDB ID – 1R4L). N-glycosylation and its sialylation at sites N90 and N322 could a play critical role in the binding of hACE2 with RBD of S protein but the glycosylation may not have direct influence on inhibitor binding or hACE2 activity

We evaluated the proximity of the N-glycans to the peptidase activity and inhibitor binding sites of hACE2 (Figure 1). Inspection of the co-crystallization structure of hACE2 (PDB ID – 1R4L) with its inhibitor shows that the inhibitor is bound inside the cleft formed by the two monomers of ACE2, which is also the site of peptidase enzymatic activity (Towler, P., Staker, B., et al. 2004). The RBD-binding region with its N-glycans, on the other hand, is located outside the cleft, nearly on the opposite face of each ACE2 monomer, as shown in Figure 1.

### Mapping N-glycosylation on hACE2

For the amino acid numbering we have followed the numbering used in UniProt [Q9BYF1] and the literature, although the numbering differs from that of the recombinant hACE2 we used for the study, which is missing the 1-17 signal peptide. Initially, we conducted a combination of trypsin and chymotrypsin digests to map the N-glycosylation on hACE2. Even though all the sites were covered by this digestion, the glycoform assignment at site N53 was ambiguous, as the peptide + HexNAc fragment (Y1 ion) was missing from the spectra. Thus, we conducted a combination of GluC and chymotrypsin digestion to characterize the N-glycosylation at site N53. To ensure whether the detected glycans were indeed derived from ACE2, we searched the LC-MS/MS data of the ACE2 digest against the complete human UniProt database (downloaded on Aug.2018) and validated that hACE2 was the most abundant protein in the digest with a log probability value of 511.82, while the second abundant protein had a log probability value of 83.49 (Table S2). Beside ACE2, no other detected proteins showed presence of glycopeptide spectra with acceptable fragment ions. We observed substantial (73.2 to 100 % total site occupancy rate) glycan occupancy at all seven predicted N-glycosylation sites of hACE2 (Figures 2, 3, and S1-S7). While sites N53, N90, N103, N322, N546 showed no unglycosylated peptides, sites N432 and N690 were 26.8 % and 1.1 % unoccupied, respectively. Complex-type, biantennary glycans were much more abundant than high-mannose and hybrid type glycans across all N-glycosylation sites. We discovered highly processed sialylated, complex-type bi-antennary, tri-antennary and tetra-antennary glycans on all sites (Figures 2 and 3). Contrasting sialylation and glycan extension patterns were observed among the N-glycosylation sites. Site N53 featured predominantly sialylated and extended complex type glycans, whereas sites N90, N103, N322, and N546 carried abundant non-sialylated, bianntennary structures. Sites N432 and N690 were occupied with sialylated and non-sialylated N-glycans of almost equal abundance. All sites showed high levels of mono-fucosylated glycans.

**Figure 2:**
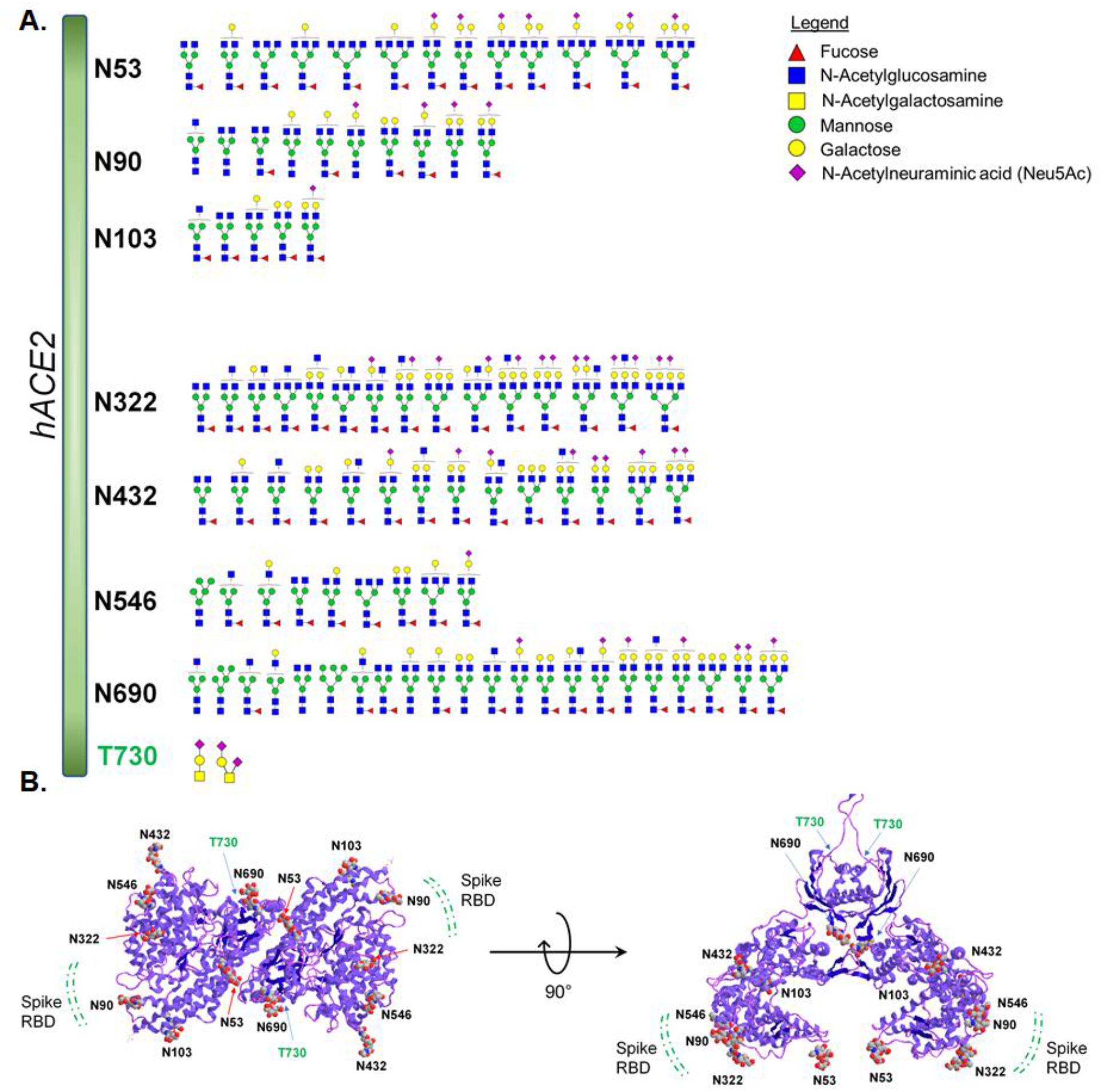
Glycosylation profile on hACE2 characterized by high-resolution LC-MS/MS. **A**. All the seven potential N-glycosylation sites were found occupied along with one O-glycosylation site bearing core-1 type O-glycans. Mostly complex type glycans were observed in all N-glycosylation sites. Some N-glycosylation sites were partially glycosylated. Monosaccharide symbols follow the SNFG (Symbol Nomenclature for Glycans) system (Varki, A., Cummings, R.D., et al. 2015). **B**. 3D model showing the location of N- and O-glycans on hACE2 dimer (PDB ID – 6M18). The potential binding location of SARS-CoV-2 RBD with hACE2 receptor is predicted based on co-crystallization data available on SARS-CoV-1 with hACE2 and the N-glycans at N90 and N322 could influence the receptor binding.

**Figure 3:**
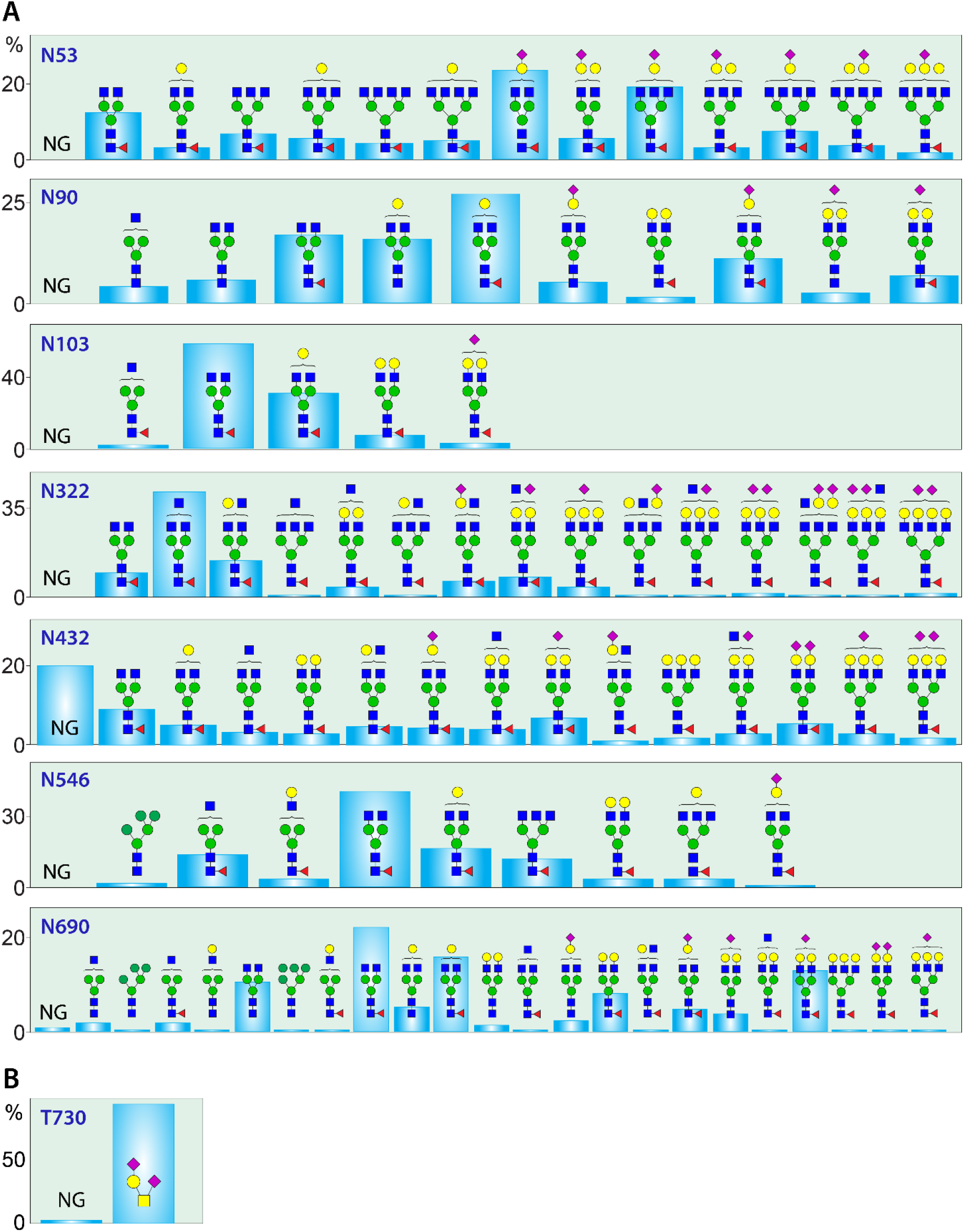
Quantitative glycosylation profile of glycans on hACE2 characterized by high-resolution LC-MS/MS. **A**. relative ratio of non-glycosylated peptide and glycoforms detected on seven N-glycosylation sites; **B**. Relative ratio of non-glycosylated peptide and glycoforms detected on O-glycosylated site Thr730 (T730). RA – Relative abundances. NG-non-glycosylated peptide. Monosaccharide symbols follow the SNFG system (Varki, A., Cummings, R.D., et al. 2015).

The N-glycan structures at each site were confirmed by the presence of characteristic peptide fragments, glycan oxonium ions and neutral losses, in addition to high resolution precursor mass determination (Figures 4A, S1-S7, Table S2).

**Figure 4:**
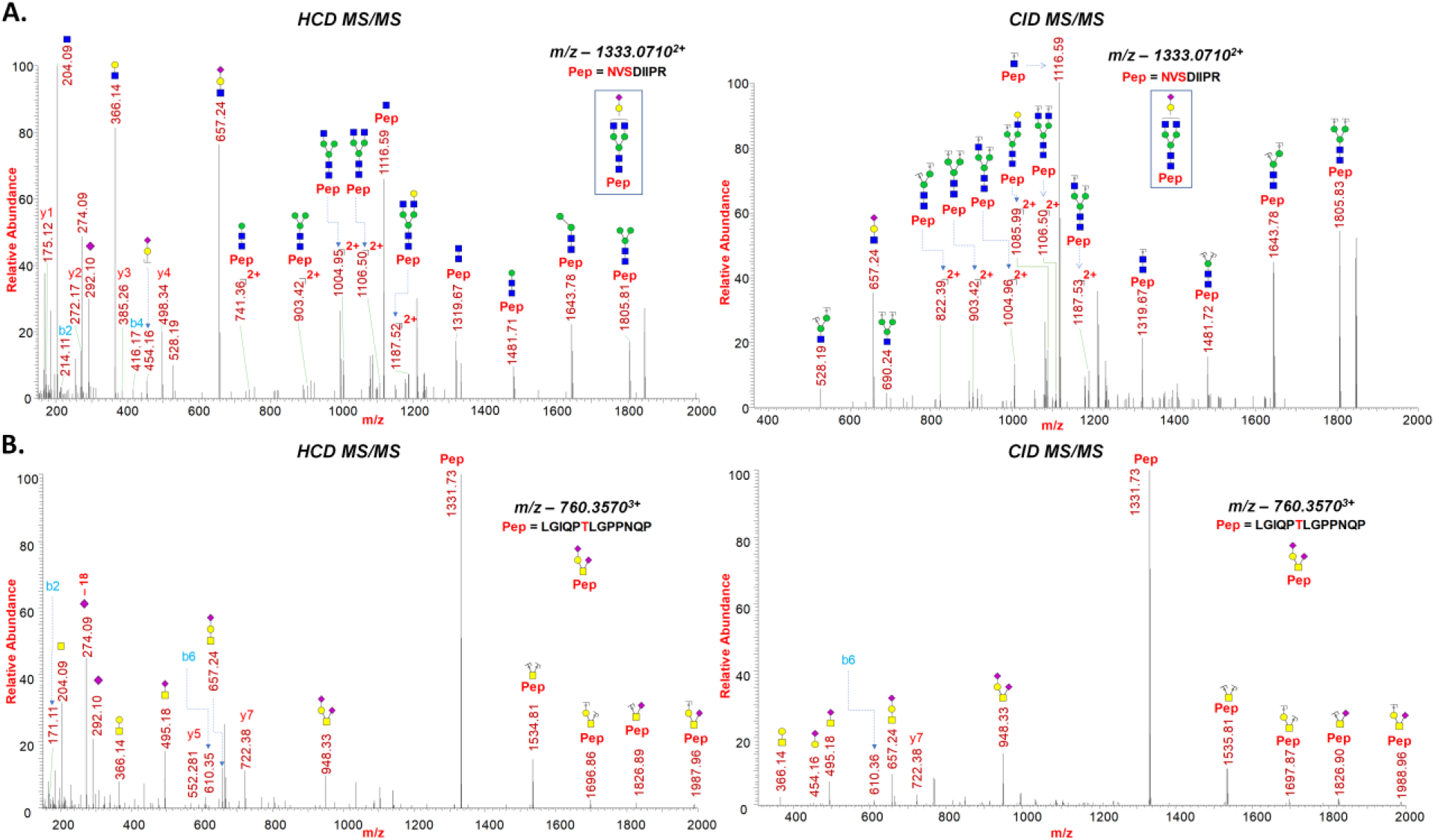
**A**. Representative HCD and CID MS/MS spectra of intact N-glycopeptide NVSDIIPR with assigned N-glycan (GlcNAc_2_Man_3_GlcNAc_2_Gal_1_NeuAc_1_) at N690, **B**. Representative HCD and CID MS/MS spectra of intact O-glycopeptide with assigned O-glycan (GalNAc_1_Gal_1_NeuAc_2_) at Thr730.

MS fragmentation of glycopeptides by collision-induced dissociation (CID) preferentially dissociates glycans while generating few peptide backbone fragments, which complicates the identification of peptide sequences. Moreover, low m/z signature ions of glycopeptides, such as glycan oxonium ions may not be detected in the CID MS/MS spectra. Conversely, HCD enables detection of small diagnostic oxonium ions of monosaccharides and the peptide backbone while preserving labile modifications at the glycan cores (Cao, L., Qu, Y., et al. 2016). Thus, by employing tandem HCD and CID fragmentation we obtained a wide spectrum of characteristic fragments of glycopeptides such as oxonium ions, peptide fragments, core glycan-peptide fragments (Y1, Y2 ions) and glycan neutral loss fragments (Figure 4A).

### Identification of O-glycosylation on hACE2

Through a glycoproteomics approach, we have identified, one O-glycosylation site on hACE2 by searching the LC-MS/MS data for common O-glycosylation modifications. We have observed very strong evidence for the presence of O-glycosylation at site Thr730 on peptide LGIQPTLGPPNQP, as the precursor masses, oxonium ions, neutral losses, and peptides fragments (b and y ions) were detected with high mass accuracy (Figures 2, 3, 4B, and S8, Table S2).

The Core-1 mucin type O-glycan GalNAcGalNeuAc_2_ was observed as the predominant glycan on site Thr730 (Figures 2, 3, and 4B). Almost 97 % of the peptide with Thr730 was occupied by O-glycan GalNAcGalNeuAc_2_ (Figures 3, 4B, and S8). Even though we observed minor peak of O-glycan corresponding to GalNAcGalNeuAc in the chromatogram, we did not consider that for the calculation since such structure can be generated by in-source fragmentation.

### Comparison of glycosylation sites of human ACE2 with other related species

We aligned the amino acid sequence of human, bat, pig, cat, mouse, rabbit, Malayan pangolin, and chicken ACE2 and compared the N-glycosylation sites among these species. This is shown schematically in Figure 5. Whilst displaying overall sequence similarities of about 53 percent, human ACE2 possesses 7, bat (Chinese rufous horseshoe bat) ACE2 also 7, cat ACE2 a total of 9, porcine ACE2 8, murine ACE2 only 6, rabbit ACE2 8, pangolin ACE2 7, and chicken ACE2 10 potential N-glycosylation sites. Our sequence alignment studies showed that in human ACE2, five N-glycosylation sites share similarities with bat ACE2, but only three N-glycosylation sites share similarities with mouse ACE2, which is not susceptible to SARS-CoV-1 binding. Pig ACE2 showed four similar sites with human, whereas cat ACE2 showed five sites that are similar (Figure 5). Based on recent reports, human, bat, cat, rabbit, and pangolin ACE2 exhibit binding affinity to SARS-CoV-2 viral RBD, while pig, mouse, and chicken ACE2 do not bind (Shi, J., Wen, Z., et al. 2020, Zhao, X., Chen, D., et al. 2020). Overall alignment of N-glycosylation sites among all these 8 species indicate that site N90 is the site that stands out among the susceptible and non-susceptible species. The susceptible species have an N90 glycosylation site, whereas the non-susceptible species do not. Interestingly, site N90 is closer to the SARS-CoV-2 RBD binding domain of ACE2 with respect to other sites, as there are reports indicating that abrogation of N90 glycosylation enhances the RBD binding with ACE2 (Procko, E. 2020, Stawiski, E.W., Diwanji, D., et al. 2020). Site N322 which is proximal to the SARS-CoV-2 RBD binding domain is aligned in human, bat, cat, pig and chicken but pig and chicken among them are non-susceptible.

**Figure 5:**
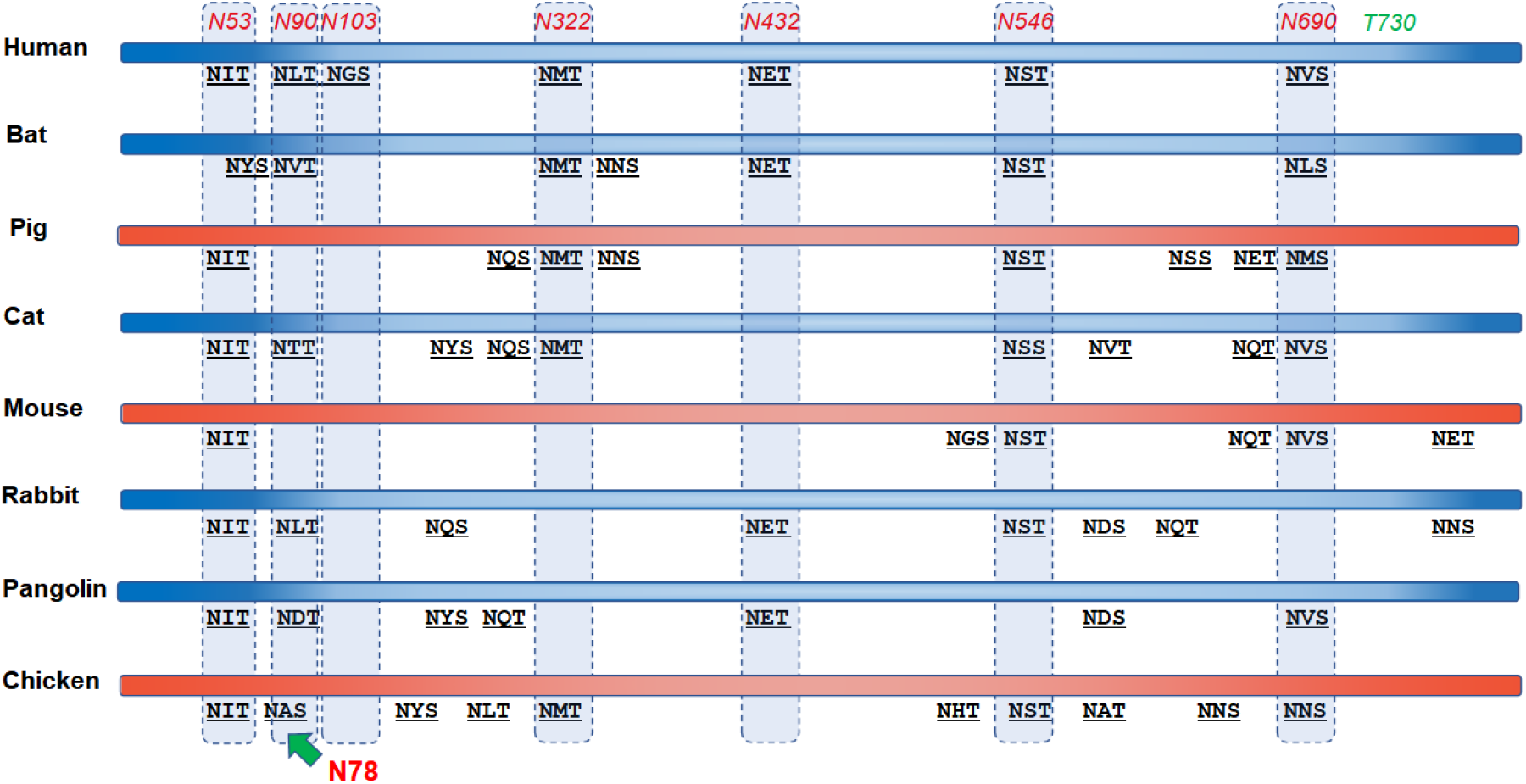
Schematic amino acid sequence alignment of glycosylation sites of ACE2 receptor from human, bat, pig, cat, mouse, rabbit, Malayan pangolin and chicken (UniProt identifiers human [Q9BYF1], bat [U5WHY8], pig [K7GLM4], cat [Q56H28], mouse [Q8R0I0], rabbit [G1TEF4], pangolin [XP_017505746.1], chicken [F1NHR4]). The number of N-linked glycosylation motifs, -NXS/T- (X≠P), accounts for 7 potential sites in the human ACE2, 7 potential N-linked glycosylation sites in the bat ACE2, 8 potential N-glycan sites in the porcine ACE2, 9 potential sites in the cat ACE2, 6 potential N-linked glycosylation sites in the murine ACE2, 8 sites in rabbit ACE2, 7 sites in pangolin ACE2, and 10 sites in chicken ACE2. Blue colored are susceptible species and orange are non-susceptible species.

### N- and O-glycomics analysis on hACE2

N- and O-glycomic studies of the hACE2 receptor were performed by methods described previously (Shajahan, A., Heiss, C., et al. 2017). Briefly, N-glycans were released by treating the reduced and alkylated hACE2 protein with the endoglycosidase PNGase F. The released N-glycans were separated by passage through a C18 solid phase extraction (SPE) cartridge, and de-N-glycosylated proteins on the cartridges were eluted with 80% acetonitrile with 0.1% formic acid. The O-linked glycans were then released from hACE2 by reductive *β*-elimination. The released N- and O- glycan fractions were then permethylated, a derivatization method that, in addition to increasing sensitivity, allows for further structural and linkage characterization *via* MS^n^ fragmentation. Moreover, permethylation allows unambiguous assignment of monosaccharide topology such as core and antennae fucosylation as it prevents monosaccharide migration or rearrangement at gaseous phase during collisional dissociation (Wuhrer, M., Deelder, A.M., et al. 2011). The permethylated N- and O- glycans were evaluated by MALDI-MS and the structures were assigned by ESI-MS^n^ performed through direct infusion (Figures 6, 7, S9-S11).

**Figure 6:**
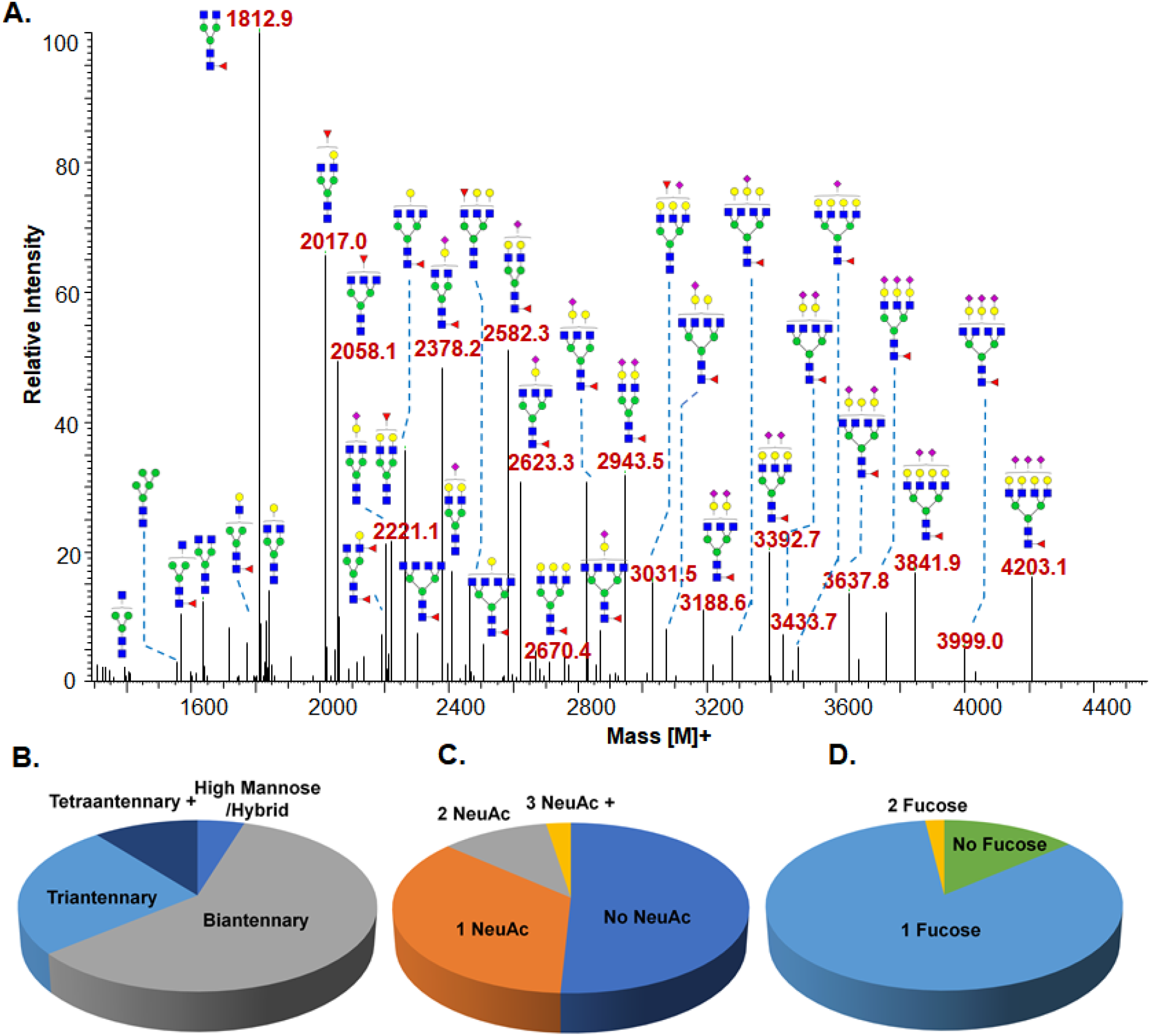
N-glycomics profiling of hACE2: N-glycans were released, purified, and permethylated prior to analysis with ESI-MS^n^. **A**. Deconvoluted profile of permethylated N-glycans from hACE*2* (Only major glycoforms are shown). Masses are shown as deconvoluted, molecular masses. The N-glycan structure were confirmed by ESI-MS^n^; **B**. Percentage breakdowns of N-glycan class: High mannose/Hybrid 5%; Biantennary 59%; Triantennary 25%; Tetraantennary 11%. **C**. Percentage breakdowns of Sialyation: Nonsialylated 51%; 1 NeuAc 36%; 2 NeuAc 11%; 3< NeuAc 3%. **D**. Percentage breakdown of Fucosylation: No Fucose 14%; 1 Fucose 84%; 2 Fucose 2%.

**Figure 7:**
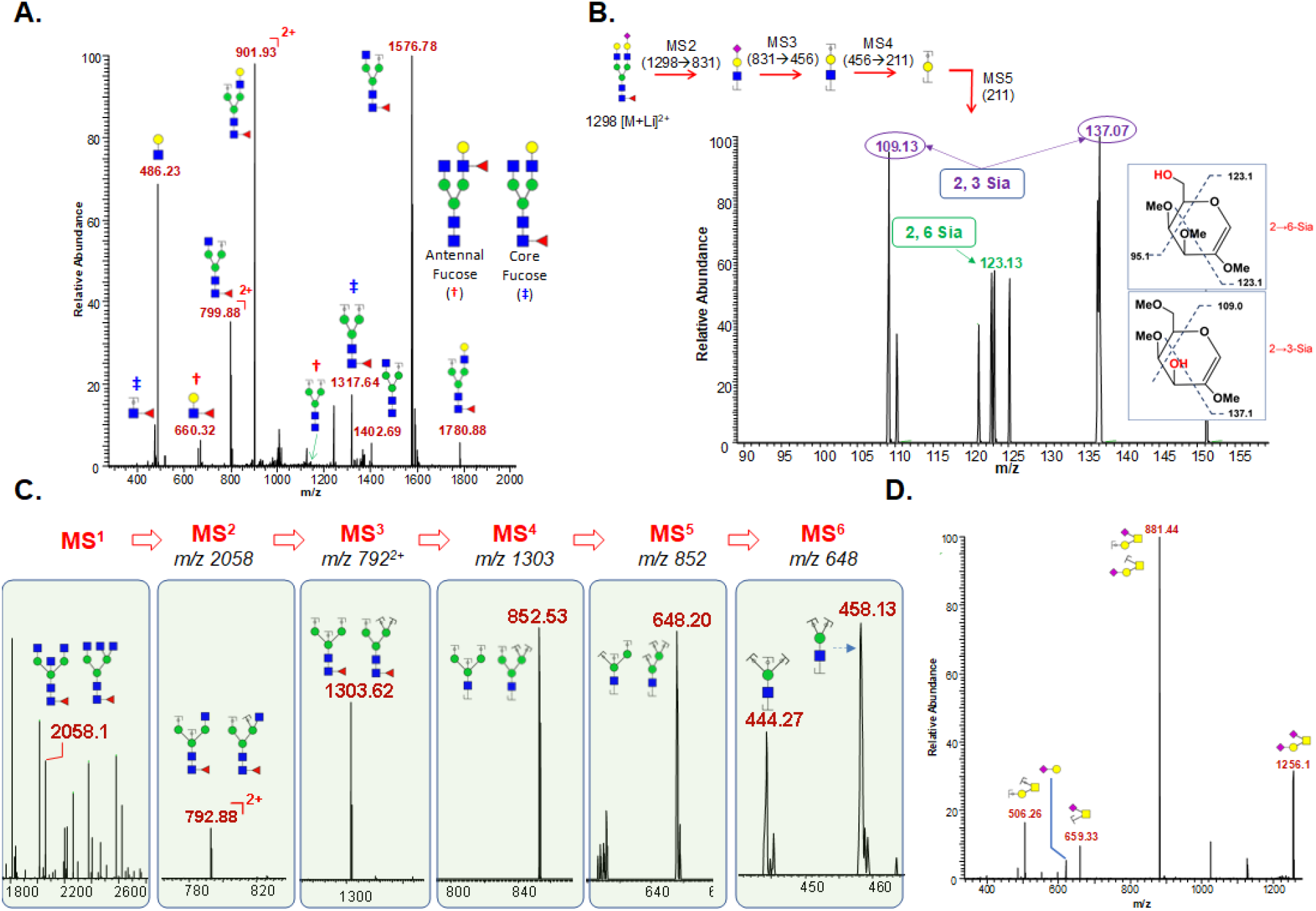
**A**. MS^2^ Fragmentation of an N-glycan observed at m/z 1133 (sodiated; z=2). MS^2^ Fragmentation reveals that there is a mixture of glycoforms that contain either core-linked or antennally-linked fucose; **B**. MS^5^ Fragmentation of a sialylated N-glycan observed at m/z 1298 (lithium adduct; z=2). MS^5^ Fragmentation breaks down the antennal arm to the sialylated galactose. Cross-ring fragmentation determines whether the sialyation was 2,3 or 2,6 linked; **C**: MS^6^ Fragmentation of a complex-type N-glycan observed at m/z 1052 (sodium adduct; z=2). MS^6^ Fragmentation breaks down the structure to the trimannose core. Fragmentation reveals the number of substituents, consistent with bisecting GlcNAc. **D**: MS^2^ Fragmentation of an O-glycan observed at m/z 1256 (sodiated; z=1).

Sequencing of the N-glycans obtained from hACE2 by MALDI-MS and ESI-MS^n^ indicated a highly diverse pool of glycoforms comprised of high mannose, hybrid, and complex-type structures (Figures 6 and S9, Table 1). The glycans were predominantly of the complex type and were primarily (~60%) biantennary, with triantennary and quaternary structures also present in significant amounts. More than 85% of the structures were fucosylated, and roughly half of the structures were sialylated. The glycomics results corroborate with the glycoproteomics results except that few minor glycoforms present on the glycomics data were not annotated to the N-glycosylation sites of ACE2 because of the lack of their MS^2^ scans on the LC-MS/MS data. The glycan structures were annotated based on high resolution precursor mass, ESI-MS^n^ fragmentation and the common biosynthetic pathways of mammalian N-glycans (Figure S11).

**Table I.**
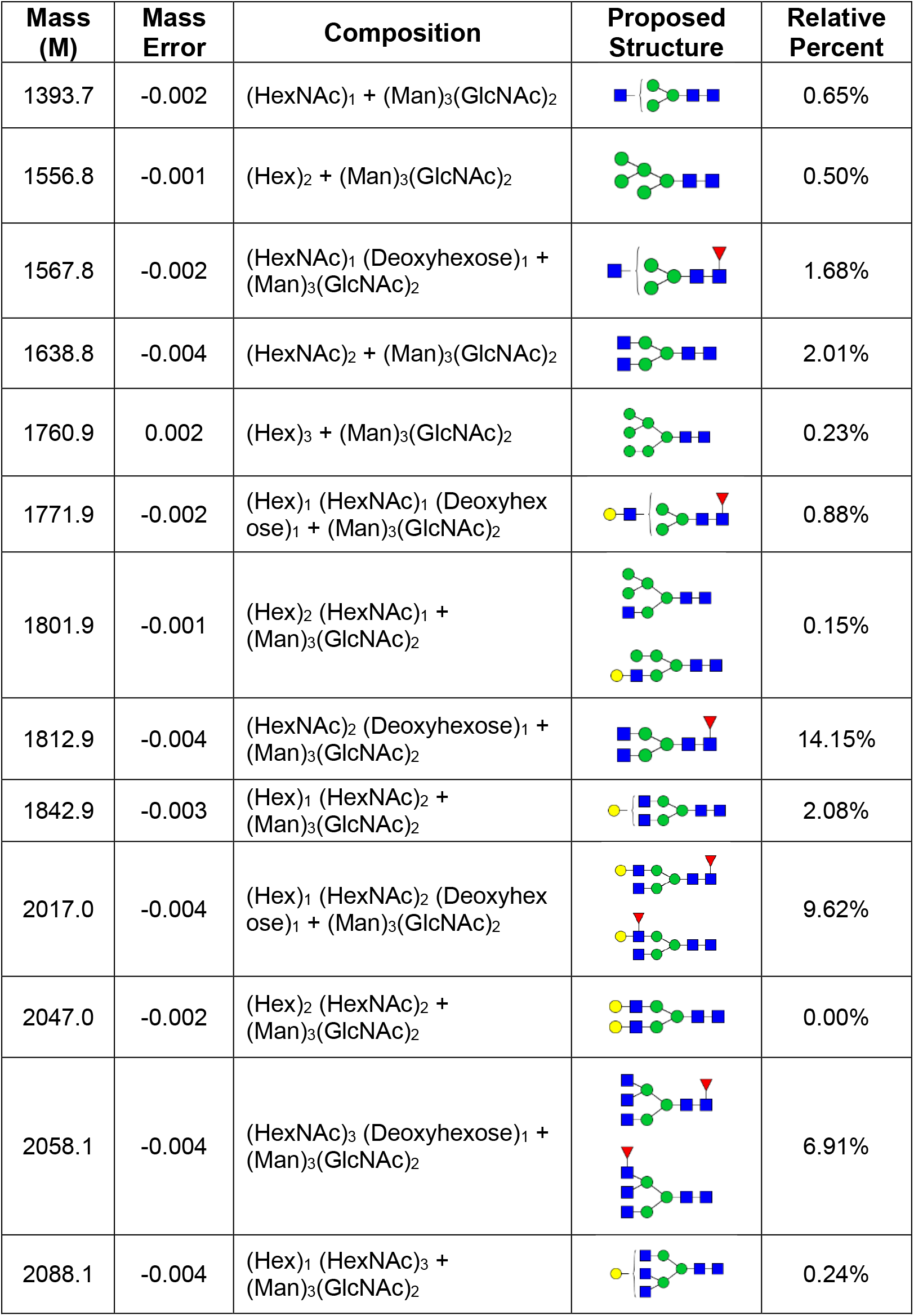
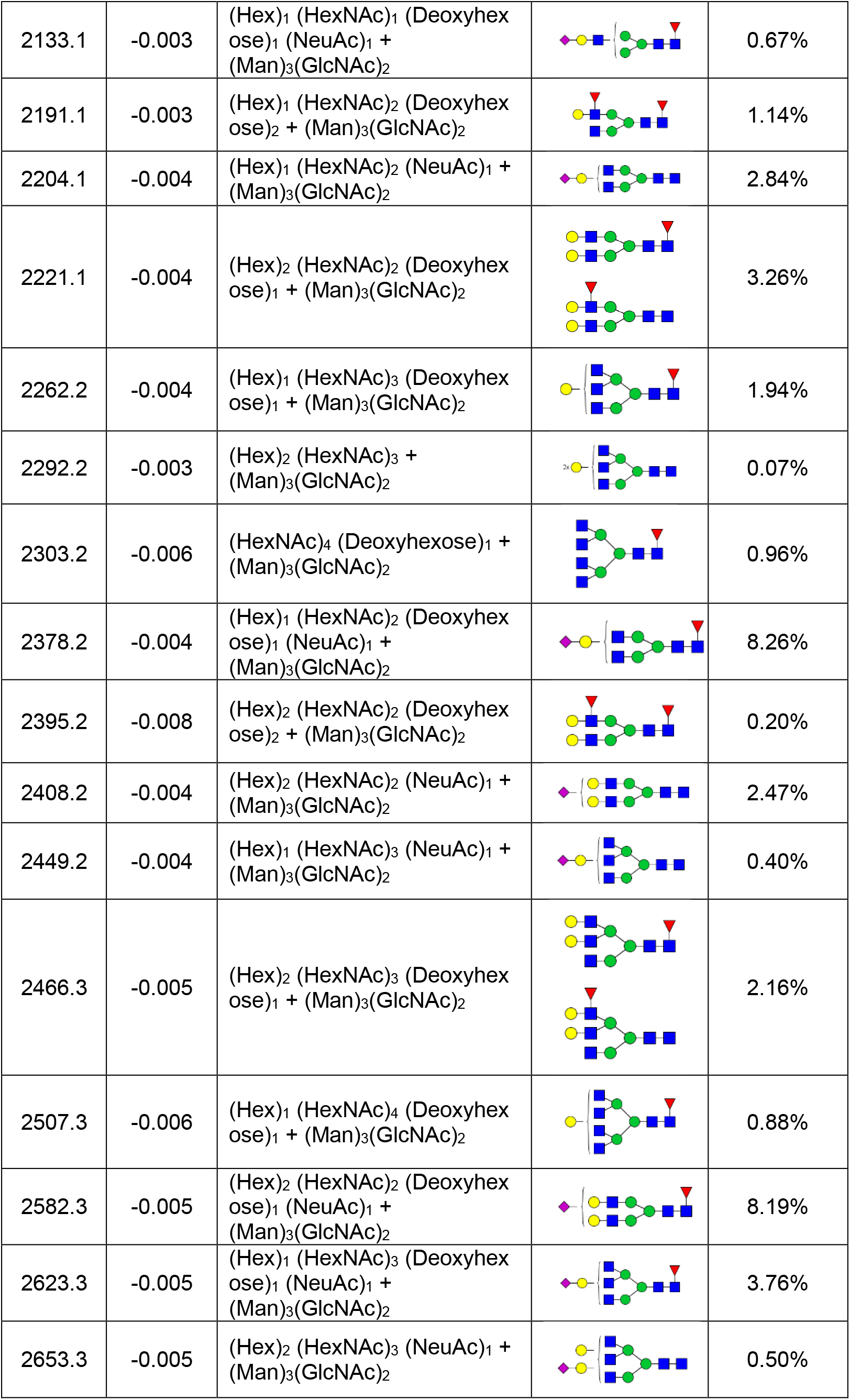
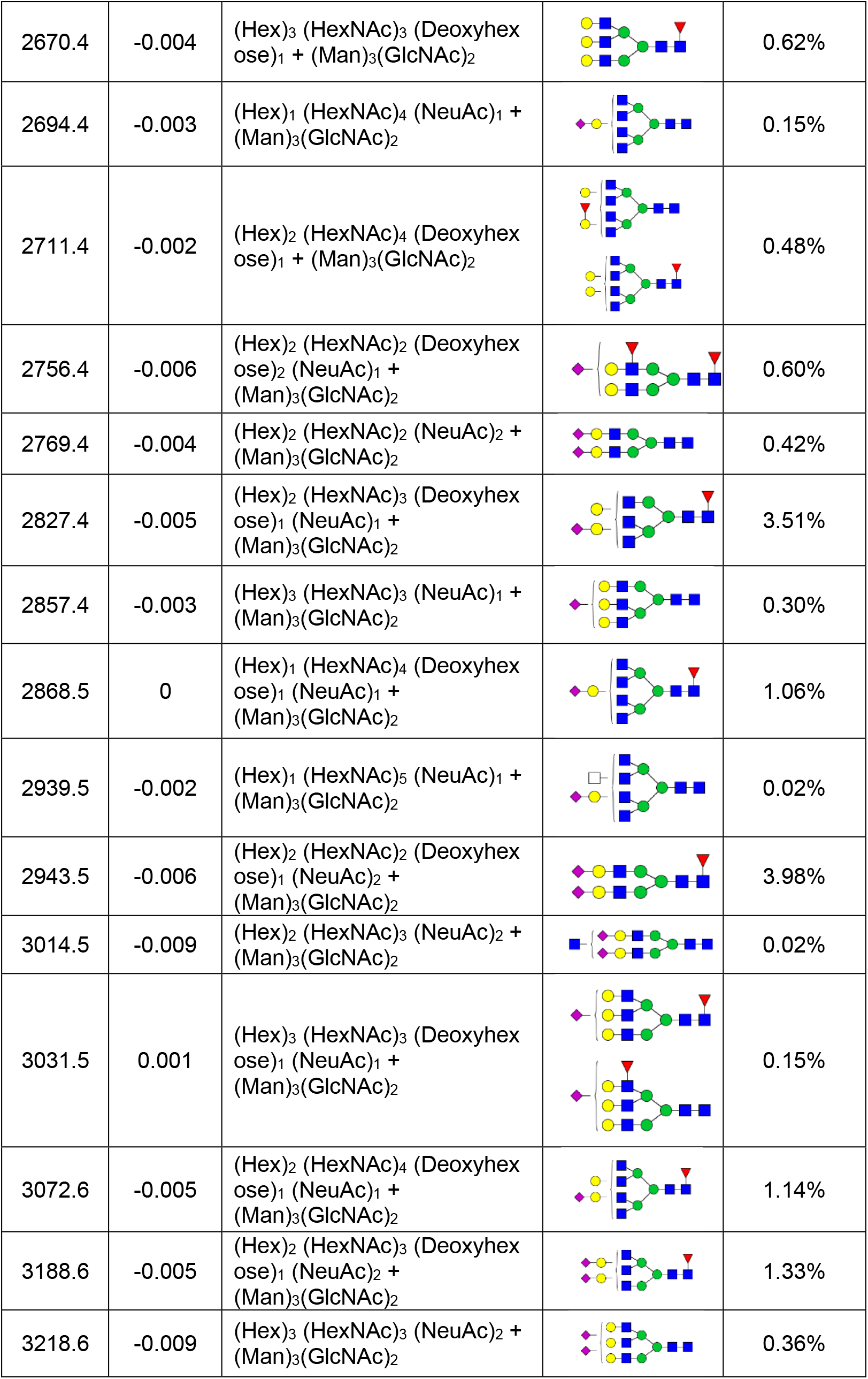
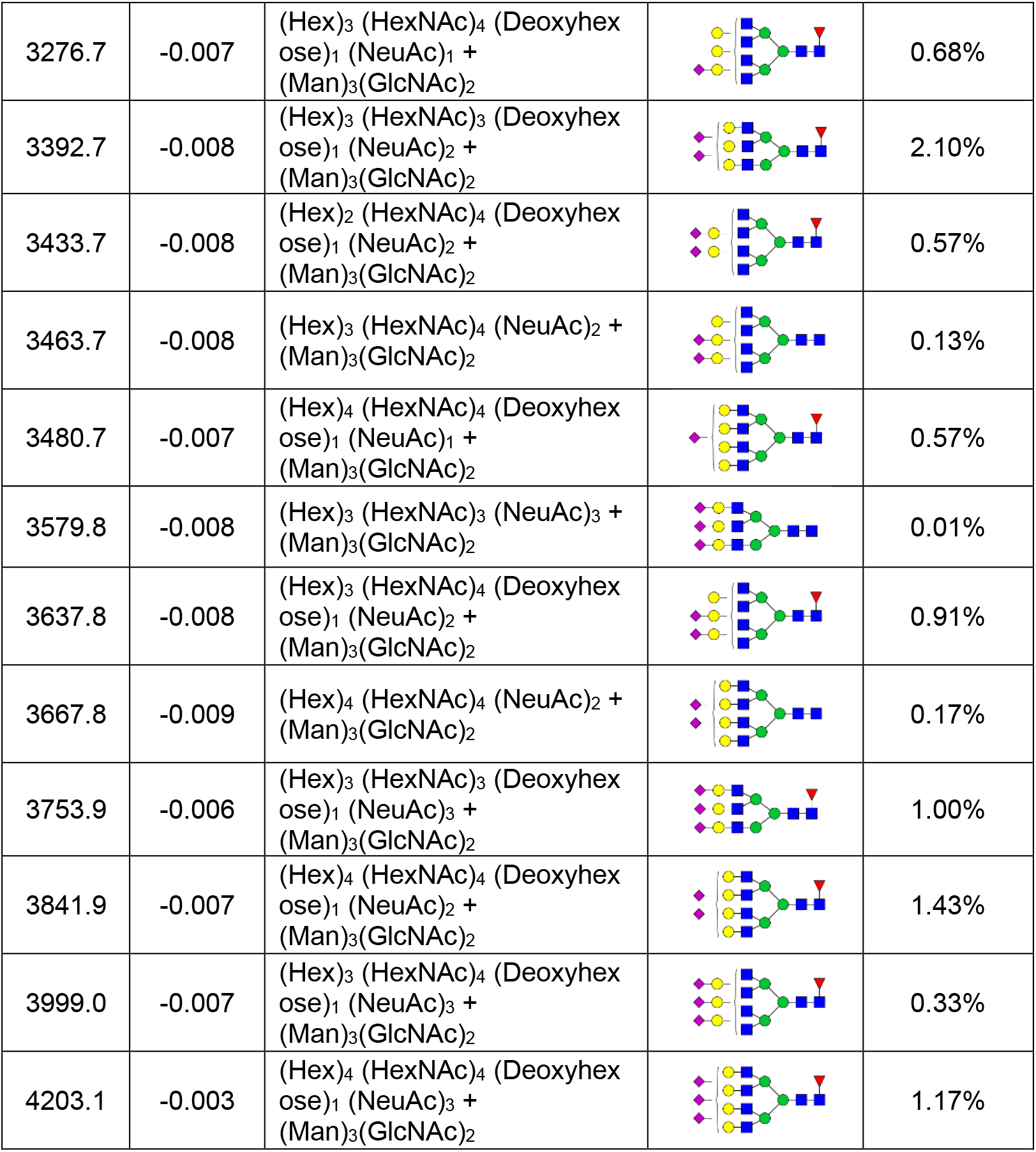
Relative abundances and structures of N-glycans detected by glycomics analysis on hACE2. Masses of permethylated glycans are represented as molecular masses [M+] and the structures were assigned based on common mammalian N-glycosylation synthesis pathway and also by ESI-MS^n^ analysis.

| Mass (M) | Mass Error | Composition | Proposed Structure | Relative Percent |
| --- | --- | --- | --- | --- |
| 1393.7 | -0.002 | (HexNAc)<sub>1</sub> + (Man)<sub>3</sub>(GlcNAc)<sub>2</sub> |  | 0.65% |
| 1556.8 | -0.001 | (Hex)<sub>2</sub> + (Man)<sub>3</sub>(GlcNAc)<sub>2</sub> |  | 0.50% |
| 1567.8 | -0.002 | (HexNAc)<sub>1</sub> (Deoxyhexose)<sub>1</sub> + (Man)<sub>3</sub>(GlcNAc)<sub>2</sub> |  | 1.68% |
| 1638.8 | -0.004 | (HexNAc)<sub>2</sub> + (Man)<sub>3</sub>(GlcNAc)<sub>2</sub> |  | 2.01% |
| 1760.9 | 0.002 | (Hex)<sub>3</sub> + (Man)<sub>3</sub>(GlcNAc)<sub>2</sub> |  | 0.23% |
| 1771.9 | -0.002 | (Hex)<sub>1</sub> (HexNAc)<sub>1</sub> (Deoxyhexose)<sub>1</sub> + (Man)<sub>3</sub>(GlcNAc)<sub>2</sub> |  | 0.88% |
| 1801.9 | -0.001 | (Hex)<sub>2</sub> (HexNAc)<sub>1</sub> + (Man)<sub>3</sub>(GlcNAc)<sub>2</sub> |  | 0.15% |
| 1812.9 | -0.004 | (HexNAc)<sub>2</sub> (Deoxyhexose)<sub>1</sub> + (Man)<sub>3</sub>(GlcNAc)<sub>2</sub> |  | 14.15% |
| 1842.9 | -0.003 | (Hex)<sub>1</sub> (HexNAc)<sub>2</sub> + (Man)<sub>3</sub>(GlcNAc)<sub>2</sub> |  | 2.08% |
| 2017.0 | -0.004 | (Hex)<sub>1</sub> (HexNAc)<sub>2</sub> (Deoxyhexose)<sub>1</sub> + (Man)<sub>3</sub>(GlcNAc)<sub>2</sub> |  | 9.62% |
| 2047.0 | -0.002 | (Hex)<sub>2</sub> (HexNAc)<sub>2</sub> + (Man)<sub>3</sub>(GlcNAc)<sub>2</sub> |  | 0.00% |
| 2058.1 | -0.004 | (HexNAc)<sub>3</sub> (Deoxyhexose)<sub>1</sub> + (Man)<sub>3</sub>(GlcNAc)<sub>2</sub> |  | 6.91% |
| 2088.1 | -0.004 | (Hex)<sub>1</sub> (HexNAc)<sub>3</sub> + (Man)<sub>3</sub>(GlcNAc)<sub>2</sub> |  | 0.24% |
| 2133.1 | -0.003 | (Hex)<sub>1</sub> (HexNAc)<sub>1</sub> (Deoxyhexose)<sub>1</sub> (NeuAc)<sub>1</sub> + (Man)<sub>3</sub>(GlcNAc)<sub>2</sub> |  | 0.67% |
| 2191.1 | -0.003 | (Hex)<sub>1</sub> (HexNAc)<sub>2</sub> (Deoxyhexose)<sub>2</sub> + (Man)<sub>3</sub>(GlcNAc)<sub>2</sub> |  | 1.14% |
| 2204.1 | -0.004 | (Hex)<sub>1</sub> (HexNAc)<sub>2</sub> (NeuAc)<sub>1</sub> + (Man)<sub>3</sub>(GlcNAc)<sub>2</sub> |  | 2.84% |
| 2221.1 | -0.004 | (Hex)<sub>2</sub> (HexNAc)<sub>2</sub> (Deoxyhexose)<sub>1</sub> + (Man)<sub>3</sub>(GlcNAc)<sub>2</sub> |  | 3.26% |
| 2262.2 | -0.004 | (Hex)<sub>1</sub> (HexNAc)<sub>3</sub> (Deoxyhexose)<sub>1</sub> + (Man)<sub>3</sub>(GlcNAc)<sub>2</sub> |  | 1.94% |
| 2292.2 | -0.003 | (Hex)<sub>2</sub> (HexNAc)<sub>3</sub> + (Man)<sub>3</sub>(GlcNAc)<sub>2</sub> |  | 0.07% |
| 2303.2 | -0.006 | (HexNAc)<sub>4</sub> (Deoxyhexose)<sub>1</sub> + (Man)<sub>3</sub>(GlcNAc)<sub>2</sub> |  | 0.96% |
| 2378.2 | -0.004 | (Hex)<sub>1</sub> (HexNAc)<sub>2</sub> (Deoxyhexose)<sub>1</sub> (NeuAc)<sub>1</sub> + (Man)<sub>3</sub>(GlcNAc)<sub>2</sub> |  | 8.26% |
| 2395.2 | -0.008 | (Hex)<sub>2</sub> (HexNAc)<sub>2</sub> (Deoxyhexose)<sub>2</sub> + (Man)<sub>3</sub>(GlcNAc)<sub>2</sub> |  | 0.20% |
| 2408.2 | -0.004 | (Hex)<sub>2</sub> (HexNAc)<sub>2</sub> (NeuAc)<sub>1</sub> + (Man)<sub>3</sub>(GlcNAc)<sub>2</sub> |  | 2.47% |
| 2449.2 | -0.004 | (Hex)<sub>1</sub> (HexNAc)<sub>3</sub> (NeuAc)<sub>1</sub> + (Man)<sub>3</sub>(GlcNAc)<sub>2</sub> |  | 0.40% |
| 2466.3 | -0.005 | (Hex)<sub>2</sub> (HexNAc)<sub>3</sub> (Deoxyhexose)<sub>1</sub> + (Man)<sub>3</sub>(GlcNAc)<sub>2</sub> |  | 2.16% |
| 2507.3 | -0.006 | (Hex)<sub>1</sub> (HexNAc)<sub>4</sub> (Deoxyhexose)<sub>1</sub> + (Man)<sub>3</sub>(GlcNAc)<sub>2</sub> |  | 0.88% |
| 2582.3 | -0.005 | (Hex)<sub>2</sub> (HexNAc)<sub>2</sub> (Deoxyhexose)<sub>1</sub> (NeuAc)<sub>1</sub> + (Man)<sub>3</sub>(GlcNAc)<sub>2</sub> |  | 8.19% |
| 2623.3 | -0.005 | (Hex)<sub>1</sub> (HexNAc)<sub>3</sub> (Deoxyhexose)<sub>1</sub> (NeuAc)<sub>1</sub> + (Man)<sub>3</sub>(GlcNAc)<sub>2</sub> |  | 3.76% |
| 2653.3 | -0.005 | (Hex)<sub>2</sub> (HexNAc)<sub>3</sub> (NeuAc)<sub>1</sub> + (Man)<sub>3</sub>(GlcNAc)<sub>2</sub> |  | 0.50% |
| 2670.4 | -0.004 | (Hex)<sub>3</sub> (HexNAc)<sub>3</sub> (Deoxyhexose)<sub>1</sub> + (Man)<sub>3</sub>(GlcNAc)<sub>2</sub> |  | 0.62% |
| 2694.4 | -0.003 | (Hex)<sub>1</sub> (HexNAc)<sub>4</sub> (NeuAc)<sub>1</sub> + (Man)<sub>3</sub>(GlcNAc)<sub>2</sub> |  | 0.15% |
| 2711.4 | -0.002 | (Hex)<sub>2</sub> (HexNAc)<sub>4</sub> (Deoxyhexose)<sub>1</sub> + (Man)<sub>3</sub>(GlcNAc)<sub>2</sub> |  | 0.48% |
| 2756.4 | -0.006 | (Hex)<sub>2</sub> (HexNAc)<sub>2</sub> (Deoxyhexose)<sub>2</sub> (NeuAc)<sub>1</sub> + (Man)<sub>3</sub>(GlcNAc)<sub>2</sub> |  | 0.60% |
| 2769.4 | -0.004 | (Hex)<sub>2</sub> (HexNAc)<sub>2</sub> (NeuAc)<sub>2</sub> + (Man)<sub>3</sub>(GlcNAc)<sub>2</sub> |  | 0.42% |
| 2827.4 | -0.005 | (Hex)<sub>2</sub> (HexNAc)<sub>3</sub> (Deoxyhexose)<sub>1</sub> (NeuAc)<sub>1</sub> + (Man)<sub>3</sub>(GlcNAc)<sub>2</sub> |  | 3.51% |
| 2857.4 | -0.003 | (Hex)<sub>3</sub> (HexNAc)<sub>3</sub> (NeuAc)<sub>1</sub> + (Man)<sub>3</sub>(GlcNAc)<sub>2</sub> |  | 0.30% |
| 2868.5 | 0 | (Hex)<sub>1</sub> (HexNAc)<sub>4</sub> (Deoxyhexose)<sub>1</sub> (NeuAc)<sub>1</sub> + (Man)<sub>3</sub>(GlcNAc)<sub>2</sub> |  | 1.06% |
| 2939.5 | -0.002 | (Hex)<sub>1</sub> (HexNAc)<sub>5</sub> (NeuAc)<sub>1</sub> + (Man)<sub>3</sub>(GlcNAc)<sub>2</sub> |  | 0.02% |
| 2943.5 | -0.006 | (Hex)<sub>2</sub> (HexNAc)<sub>2</sub> (Deoxyhexose)<sub>1</sub> (NeuAc)<sub>2</sub> + (Man)<sub>3</sub>(GlcNAc)<sub>2</sub> |  | 3.98% |
| 3014.5 | -0.009 | (Hex)<sub>2</sub> (HexNAc)<sub>3</sub> (NeuAc)<sub>2</sub> + (Man)<sub>3</sub>(GlcNAc)<sub>2</sub> |  | 0.02% |
| 3031.5 | 0.001 | (Hex)<sub>3</sub> (HexNAc)<sub>3</sub> (Deoxyhexose)<sub>1</sub> (NeuAc)<sub>1</sub> + (Man)<sub>3</sub>(GlcNAc)<sub>2</sub> |  | 0.15% |
| 3072.6 | -0.005 | (Hex)<sub>2</sub> (HexNAc)<sub>4</sub> (Deoxyhexose)<sub>1</sub> (NeuAc)<sub>1</sub> + (Man)<sub>3</sub>(GlcNAc)<sub>2</sub> |  | 1.14% |
| 3188.6 | -0.005 | (Hex)<sub>2</sub> (HexNAc)<sub>3</sub> (Deoxyhexose)<sub>1</sub> (NeuAc)<sub>2</sub> + (Man)<sub>3</sub>(GlcNAc)<sub>2</sub> |  | 1.33% |
| 3218.6 | -0.009 | (Hex)<sub>3</sub> (HexNAc)<sub>3</sub> (NeuAc)<sub>2</sub> + (Man)<sub>3</sub>(GlcNAc)<sub>2</sub> |  | 0.36% |
| 3276.7 | -0.007 | (Hex)<sub>3</sub> (HexNAc)<sub>4</sub> (Deoxyhexose)<sub>1</sub> (NeuAc)<sub>1</sub> + (Man)<sub>3</sub>(GlcNAc)<sub>2</sub> |  | 0.68% |
| 3392.7 | -0.008 | (Hex)<sub>3</sub> (HexNAc)<sub>3</sub> (Deoxyhexose)<sub>1</sub> (NeuAc)<sub>2</sub> + (Man)<sub>3</sub>(GlcNAc)<sub>2</sub> |  | 2.10% |
| 3433.7 | -0.008 | (Hex)<sub>2</sub> (HexNAc)<sub>4</sub> (Deoxyhexose)<sub>1</sub> (NeuAc)<sub>2</sub> + (Man)<sub>3</sub>(GlcNAc)<sub>2</sub> |  | 0.57% |
| 3463.7 | -0.008 | (Hex)<sub>3</sub> (HexNAc)<sub>4</sub> (NeuAc)<sub>2</sub> + (Man)<sub>3</sub>(GlcNAc)<sub>2</sub> |  | 0.13% |
| 3480.7 | -0.007 | (Hex)<sub>4</sub> (HexNAc)<sub>4</sub> (Deoxyhexose)<sub>1</sub> (NeuAc)<sub>1</sub> + (Man)<sub>3</sub>(GlcNAc)<sub>2</sub> |  | 0.57% |
| 3579.8 | -0.008 | (Hex)<sub>3</sub> (HexNAc)<sub>3</sub> (NeuAc)<sub>3</sub> + (Man)<sub>3</sub>(GlcNAc)<sub>2</sub> |  | 0.01% |
| 3637.8 | -0.008 | (Hex)<sub>3</sub> (HexNAc)<sub>4</sub> (Deoxyhexose)<sub>1</sub> (NeuAc)<sub>2</sub> + (Man)<sub>3</sub>(GlcNAc)<sub>2</sub> |  | 0.91% |
| 3667.8 | -0.009 | (Hex)<sub>4</sub> (HexNAc)<sub>4</sub> (NeuAc)<sub>2</sub> + (Man)<sub>3</sub>(GlcNAc)<sub>2</sub> |  | 0.17% |
| 3753.9 | -0.006 | (Hex)<sub>3</sub> (HexNAc)<sub>3</sub> (Deoxyhexose)<sub>1</sub> (NeuAc)<sub>3</sub> + (Man)<sub>3</sub>(GlcNAc)<sub>2</sub> |  | 1.00% |
| 3841.9 | -0.007 | (Hex)<sub>4</sub> (HexNAc)<sub>4</sub> (Deoxyhexose)<sub>1</sub> (NeuAc)<sub>2</sub> + (Man)<sub>3</sub>(GlcNAc)<sub>2</sub> |  | 1.43% |
| 3999.0 | -0.007 | (Hex)<sub>3</sub> (HexNAc)<sub>4</sub> (Deoxyhexose)<sub>1</sub> (NeuAc)<sub>3</sub> + (Man)<sub>3</sub>(GlcNAc)<sub>2</sub> |  | 0.33% |
| 4203.1 | -0.003 | (Hex)<sub>4</sub> (HexNAc)<sub>4</sub> (Deoxyhexose)<sub>1</sub> (NeuAc)<sub>3</sub> + (Man)<sub>3</sub>(GlcNAc)<sub>2</sub> |  | 1.17% |

### Core versus antennal fucosylation on hACE2 N-glycans

MS/MS sequencing of the observed N-glycans was conducted with an automated top-down program, collecting MS^2^ spectra of the highest intensity peaks with CID. This helped confirm overall structure (for example hybrid forms versus complex type) and determine placement of the fucose. We observed that, while most of the fucosylated structures were primarily core-fucose, small amounts of the same glycoform appeared to be fucosylated on the antennae (Figure 7A). This can be determined by a diagnostic terminal GalGlcNAcFuc fragment of m/z 660 [M+Na^+^], which corresponds to the fragmentation of the antenna (Figure 7A) appearing as a b-ion. The corresponding y-ion m/z 1402 [M+Na^+^] is also found, confirming this structure. Additional glycoforms displaying two fucoses (core and antennal) were also observed among these structures (Figure 6, Table 1).

### Identifying the sialic acid linkages on hACE2 N-glycans by ESI-MS^n^

A separate MS^n^ procedure was used to determine the linkages of sialic acids in the complex-type N-glycans in hACE2. The method, which uses direct infusion of the permethylated glycans in a lithium carbonate and methanol solution, causes sequential fragmentation of the sialylated arms of a complex-type N-glycan, down to the galactose monosaccharide residue, which carries different methyl substitution, depending on the position through which it was linked to sialic acid in the glycan. The methyl substitution is then determined by analysis of the cross-ring fragments produced from the galactose monosaccharide ion. The cross-ring fragments obtained from the fragmentation of hACE2 glycans using this MS^5^ method (Anthony, R.M., Nimmerjahn, F., et al. 2008, Shajahan, A., Supekar, N.T., et al. 2020a) provided diagnostic ions corresponding to 2→3 or 2→6 linked sialic acids (Figures 7B and S9). While the spectra demonstrated a predominance of fragments indicating 2→3 linked sialylated N-glycans, most structures also showed evidence for minor presence of 2→6 linked sialic acid. No N-glycan was found to solely contain 2→6 linked sialic acids.

### Determination of bisecting GlcNAc on hACE2 N-glycans by ESI-MS^n^

To determine presence of bisecting GlcNAc, an MS^n^ strategy to trim down the permethylated N-glycan to the trimannose core was utilized (Ashline, D.J., Duk, M., et al. 2015, Shajahan, A., Supekar, N.T., et al. 2020a). Fragmentation to the core yields a 3-substituted structure at m/z 852 [M+Na^+^], that could correspond to either a triantennary structure or a biantennary structure with a bisecting GlcNAc. Fragmentation of this ion yields an m/z of 444 [M+Na^+^] for bisecting structures or m/z 458 [M+Na^+^] for triantennary glycans. As shown in Figure 7C, both m/zs of 444 and 458 were observed, indicating that both triantennary and bisecting structures were present in this sample. This supports earlier findings by Zhao et al. who reported both bisecting and higher-order complex-type glycans (Zhao, X., Guo, F., et al. 2015).

### Detection and confirmation of O-glycans by MALDI-MS and ESI-MS^n^

The released O-glycans from hACE2 were sequenced by both MALDI-MS and ESI-MS^n^ after permethylation (Figures 7D and S10). The signal intensity was very low, and mostly the disialylated Core-1 O-glycan structure was observed. This is supported by our O-glycoproteomics findings, which showed that O-glycosylation occurred only on one site, disialylated Core-1 was the major glycoform. The MS^2^ fragmentation of the O-glycan confirms its structure as a sialylated Core-1 O-glycan (Figure 7D).

## Discussion

hACE2 acts as a receptor for human coronaviruses SARS-CoV-1 and SARS-CoV-2, as well as human coronavirus NL63/HCoV-NL63 (Hoffmann, M., Kleine-Weber, H., et al. 2020, Hofmann, H., Pyrc, K., et al. 2005, Li, *W*., Moore, M.J., et al. 2003). Recent evidence indicates that the molecular and structural features of SARS-CoV-2 RBD result in stronger hACE2 binding compared to the earlier virus SARS-CoV-1 (Andersen, K.G., Rambaut, A., et al. 2020, Hoffmann, M., Kleine-Weber, H., et al. 2020).

According to recent cryo-EM studies on SARS-CoV-2, the binding of S protein to the hACE2 receptor primarily involves extensive polar residue interactions between RBD and the peptidase domain of hACE2 (Hoffmann, M., Kleine-Weber, H., et al. 2020, Walls, A.C., Park, Y.J., et al. 2020). The RBD located in the S1 subunit of SARS-CoV-2 S protein undergoes a hinge-like dynamic movement that enhances the capture of the spike protein RBD with hACE2 (Wrapp, D., Wang, N., et al. 2020). This enhanced affinity for the human ACE2 receptor is predicted to be 10-20-fold higher for SARS-CoV-2 than SARS-CoV-1, which may be responsible for the increased transmissibility of the new virus (Wrapp, D., Wang, N., et al. 2020, Yan, R., Zhang, Y., et al. 2020). The protease domain of ACE2 interacts with the RBD of coronaviruses, and thus soluble ACE2 (sACE2), which is devoid of neck and transmembrane domains are capable of binding with RBD, neutralizing infection (Hofmann, H., Geier, M., et al. 2004, Yan, R., Zhang, Y., et al. 2020).

ACE2 is expressed in most vertebrates. The ACE2 variants from human, bat, domestic pig, domestic cat, mouse, rabbit, and pangolin are all composed of 805 amino acid residues (Figure 5), whereas chicken ACE2 contains 808 amino acids. SARS-CoV-1 and SARS-CoV-2 can infect a wide variety of organisms, including but not limited to humans, palm civets, cats and bats. *In vitro* virus infectivity studies conducted by Zhou *el al* indicated that SARS-CoV-2 is able to exploit the ACE2 proteins from humans, bats, pigs and civets to infect the cell cultures expressing the respective receptor. However, a cell culture expressing murine ACE2 was not infected (Zhou, P., Yang, X.L., et al. 2020). This is based on varying degrees of receptor recognition. Receptor recognition is largely determined by two factors, (i) the binding specificity and (ii) the binding affinity of the RBD of the virus’s spike protein to the cell entry receptor ACE2. This attachment step has been identified as a crucial limiting step for infection, as well as cross-species infection (Hou, Y., Peng, C., et al. 2010, Li, F. 2013). Our comparison of N-glycosylation sites across species indicated that human ACE shares several glycosylation sites with other species. Studying the correlation between glycosylation sites on ACE2 of different species and their susceptibility to viral binding can help in understanding the key sites involved (Figure 5). According to the data published by Qiu *et al*, the overall sequence identity of the ACE2 between different organisms and the hACE2 cannot be directly translated into prediction of transmissibility (Qiu, Y., Zhao, Y.B., et al. 2020). For example, mouse ACE2 matches with a higher sequence similarity than bat and pig, but the murine ACE2 cannot be exploited by the coronavirus, the study found (Qiu, Y., Zhao, Y.B., et al. 2020). This indicates the possibility of involvement of glycosylation sites, glycans and glycan terminal epitopes in dictating the binding affinity of coronavirus RBD with ACE2. Our sequence analysis of ACE2 among different species draws attention to the correlation of glycosylation at site N90 with susceptibility, N90 being a site proximal to viral RBD binding (Figures 1, 2B, and 5).

A recent *in silico* study by Stawiski *et al* (Stawiski, E.W., Diwanji, D., et al. 2020) predicted that glycosylation at N90 is an important modification that partly disrupts the interaction of the coronavirus with the ACE2 receptor (Figures 1 and 2B). Therefore, mutation at either N90 or T92 removes the glycosylation motif and makes the unglycosylated variant prone to interactions with SARS-CoV-2. The deleterious effect of N90 glycosylation upon binding to S protein has been experimentally confirmed by Procko (Procko, E. 2020). Also, the elucidation of the structure of murine ACE2 would be beneficial for the investigation of insusceptible organisms. It might, in turn, provide valuable insight into the molecular structures that make humans prone to infection with SARS-CoV-2 via hACE2. A previous co-crystallization study of hACE2 with its inhibitor indicates that the spike protein RBD binding region of hACE2 is distal to the surface that binds with the ACE2 inhibitors, suggesting that the use of ACE2 inhibitors may not be beneficial in preventing viral binding (Figure 1) (Towler, P., Staker, B., et al. 2004). Binding of ACE2 receptor inhibitors ultimately upregulates ACE2 receptor synthesis, resulting in an increase of available binding sites for SARS-CoV-2, leading to increased chances of COVID-19 infection (Rico-Mesa, J.S., White, A., et al. 2020).

O-glycosylation is initiated by the α-glycosidic attachment of N-acetylgalactosamine (GalNAc) to the hydroxyl group of serine or threonine, and mucin type O-glycosylation is the most common type in higher eukaryotes (Brockhausen, I. and Stanley, P. 2015, Van den Steen, P., Rudd, P.M., et al. 1998). O-glycans are involved in protein stability and function, and have been suggested to play roles in mediating pathogenic binding with human receptors (Mayr, J., Lau, K., et al. 2018). Our analysis confirmed the presence of O-glycosylation at site Thr730 unambiguously (Figures 2, 3, 4, and 7D). The frequency of occurrence of proline residues is higher adjacent to O-glycosylation sites and, in accordance with this general rule, the O-glycosylation site Thr730 was adjacent to a proline at site Pro729 (Thanka Christlet, T.H. and Veluraja, K. 2001).

Our data indicate that Thr730 is fully O-glycosylated, which could have implications in the ACE2 shedding after SARS-CoV-1 and CoV-2 infection. It has been demonstrated that inhibition of ADAM17 (ACE2 sheddase) leads to decreased ACE2 shedding, which may have a protective effect against SARS-CoV infection by reducing the viral load, possibly by preventing fusion of viral particles with cytoplasmic membranes (Haga, S., Yamamoto, N., et al. 2008, Palau, V., Riera, M., et al. 2020). It was shown that TNF-α-converting enzyme (TACE), ADAM17, and the sequence of ACE2 C-terminus are involved in SARS-S-induced ACE2 shedding, and these could be targets for anti-SARS-CoV therapeutics (Haga, S., Yamamoto, N., et al. 2008). Studies indicate that the cleavage site for the ADAM17-induced ectodomain shedding of ACE2 is localized within the juxtamembrane region amino acids 716 and 741, although a later study showed that Arg708 and Ser709 is involved with ACE2 shedding (Jia, H.P., Look, D.C., et al. 2009, Lai, Z.W., Hanchapola, I., et al. 2011). Our observation of O-glycosylation at Thr730 could facilitate future studies to understand the roles of glycosylation on the juxtamembrane region of ACE2 and its shedding. One may speculate that the bulky and hydrophilic glycosylation at Thr730 in the juxtamembrane region, just outside the cell membrane, could play a role in the dimerization and presenting of ACE2 on the cell surface.

Human and avian virus hemagglutinins (HAs), including that of the 2009 pandemic H1N1, bind glycans with sulfation, fucosylation and internal sialylation (Chandrasekaran, A., Srinivasan, A., et al. 2008, Mayr, J., Lau, K., et al. 2018). The human pandemic H1N1 influenza viruses shift the preference from avian-like α2→3-linkages to α2→6-linkages during the switch to infect humans, and a characteristic structural topology of glycans is also required for the specific binding of HA to α2→6 sialylated glycans (Chandrasekaran, A., Srinivasan, A., et al. 2008). The sialylated Core-1 glycoforms are involved in the life cycle of the influenza A virus and play a crucial role during infection (Mayr, J., Lau, K., et al. 2018). The full-length S proteins of SARS-CoV-2 and SARS-CoV-1 share almost 76% identity in amino acid sequences, whereas the N-terminal domains (NTDs) show only 53.5% homology. The NTDs of different coronavirus S proteins bind with varying avidity to different sugars, and the NTD of MERS-CoV prefers α2→3-linked sialic acid over α2→6-linked sialic acid with sulfated sialyl-Lewis X (with antennal fucose) being the preferred binding motif (Li, *W*., Hulswit, R.J.G., et al. 2017, Park, Y.J., Walls, A.C., et al. 2019). Nevertheless, we observed a relatively higher level of 2→3 linked sialylated N-glycans on hACE2, and thus the sialic acid linkage could favor the binding of coronavirus S proteins (Figure 7B).

S glycoproteins of coronaviruses HCoV-OC43 and Bat CoV mediate attachment to oligosaccharide receptors by interacting with 9-O-acetyl-sialic acid (Tortorici, M.A., Walls, A.C., et al. 2019). No sialic acid binding preference of SARS-CoV-1 or SARS-CoV-2 has been reported, and whether the sialic acid linkages on the hACE2 receptor affect virus entry remains to be determined. We have searched for 9-O-acetyl-sialic acid on the hACE2 glycans by extracting the masses of its oxonium ions on the LC-MS/MS spectra but could not detect any evidence for its presence. The expression of 9-O-acetyl-sialic acid depends on the expression level of the sialate O-acetyltransferase gene (CasD1), and only 1 to 2% of total sialic acid is 9-O-acetylated in HEK293 cells (Barnard, K.N., Wasik, B.R., et al. 2019). This could be the reason for the lack of 9-O-acetyl-sialic acid on the glycans from hACE2 we used for this study, as it is produced from HEK293 cells. We have detected both core fucosylation and antennal fucosylation on hACE2 N-glycans (Figure 7A). Moreover, we found evidence of bisecting GlcNAc on the hACE2 N-glycans (Figure 7C). The discovery of such glycan epitopes of hACE2 provides a better understanding of viral binding preferences and can guide the research for the development of suitable therapeutics.

Our N- and O- glycosylation characterization of hACE2 expressed in a human cell system through both glycoproteomics and glycomics can help future studies in understanding the roles glycans play in the function and pathogen binding of hACE2. We have conducted extensive manual interpretation for the assignment of each glycopeptide, glycan and linkage structures in order to eliminate false detections. We are currently exploring the protein polymorphism on hACE2 and how the glycosylation profile varies in the variants which alter the glycosylation sites as it is reported that natural ACE2 variants that are predicted to alter the virus-host interaction and thereby potentially alter host susceptibility (Stawiski, E.W., Diwanji, D., et al. 2020).

Detailed glycan analysis is important for the development of hACE2 or sACE2-based therapeutics which are suggested as a therapeutic measure to neutralize the viral pathogens (Hofmann, H., Geier, M., et al. 2004, Yan, R., Zhang, Y., et al. 2020). Evaluation of glycosylation on glycoprotein therapeutics produced from various human and non-human expression systems is critical from the point of view of immunogenicity, stability, as well as therapeutic efficacy (Beck, A., Cochet, O., et al. 2010, Sola, R.J. and Griebenow, K. 2010). Studies indicate that although children generally express more ACE2 than adults, they tend to experience milder symptoms of COVID-19, raising the question whether the glycosylation profile of hACE2 changes with age (Ciaglia, E., Vecchione, C., et al. 2020). Recent reports on COVID-19 infection of children, the triggering of severe cardiac problems, and protection observed in asthma patients necessitates the study on disease etiology and the contribution of hACE2 receptors (Hofmann, H., Simmons, G., et al. 2006, Jackson, D.J., Busse, *W.W*., et al. 2020). The understanding of complex sialylated N-glycans and sialylated mucin type O-glycans, on hACE2, along with their linkage and structural isomerism provides the basic structural knowledge that is useful for elucidating their interaction with viral surface protein and can aid in future therapeutic possibilities.

## Materials and methods

### Materials

Dithiothreitol (DTT), iodoacetamide (IAA), and iodomethane were purchased from Sigma Aldrich (St. Louis, MO). Sequencing-grade modified trypsin, chymotrypsin and GluC were purchased from Promega (Madison, WI). Peptide-N-Glycosidase F (PNGase F) was purchased from New England Biolabs (Ipswich, MA). All other reagents were purchased from Sigma Aldrich unless indicated otherwise. Data analysis was performed using Byonic 2.3.5 software and manually using Thermo Fisher Xcalibur 4.2. The purified human angiotensin converting enzyme hACE2 (Cat. No. 230-30165) was purchased from RayBiotech (Atlanta, GA).

### Experimental Design and Statistical Rationale

We have procured recombinant hACE2 receptor expressed on human HEK293 cells. The protein was expressed with a C-terminal His tag and comprises residues Gln18 to Ser740. We performed a combination of trypsin (1:20 enzyme protein ratio) and chymotrypsin (1:20 enzyme protein ratio) digestion on reduced and alkylated hACE2 and generated glycopeptides, each containing a single N-linked glycan site (based on *in silico* digestion). In order to map the site N53 unambiguously we conducted sequential digest of reduced/alkylated hACE2 with GluC (1:20 enzyme protein ratio) and chymotrypsin (1:20 enzyme protein ratio). The glycopeptides were directly subjected to two injections of high-resolution LC-MS/MS, using a glycan oxonium ion product dependent HCD triggered CID program. The dynamic exclusion was enabled for the data dependent scan while ensuring acquisition of at least two MS^2^ fragmentations of each peak. The LC-MS/MS data were analyzed using a Byonic (version 2.3.5) software search, each glycopeptide annotation was screened thoroughly manually for the b and y ions, glycan oxonium ions and neutral losses, and false detections were eliminated. We have quantified the relative intensities of glycans at each site by evaluating the area under the curve of each extracted glycopeptide peak in the LC-MS chromatogram.

For the glycomics analysis, the permethylated N- and O- glycans were analyzed by both MALDI-MS and high-resolution direct infusion ESI-MS^n^. Multiple scans were acquired, masses of glycans with less than 5 ppm accuracy were considered, and each structure was assigned based on MS^n^ data acquired at high resolution except for the MS^4-6^ experiments, where the acquisition was conducted in the ion trap (IT). The glycomics data analysis was performed manually with the help of GlycoWorkbench software. Deconvolution of the spectra to the molecular mass was conducted by using Thermo Fisher FreeStyle 1.4 software.

### Reduction, alkylation and protease digestion of hACE2 for glycoproteomics

The purified hACE2 (20 μg) expressed on HEK293 cells in 50 mM ammonium bicarbonate solution was reduced by adding 25 mM DTT and incubating at 60 °C for 30 min. The protein was further alkylated by the addition of 90 mM IAA and incubating at RT for 30 min in the dark. Subsequently, the protein was desalted using a 10-kDa centrifuge filter and digested by sequential treatment with trypsin and chymotrypsin or GluC and chymotrypsin by incubating for 18 h at 37 °C during each digestion step. The digest was filtered through a 0.2-μm filter and directly analyzed by LC-MS/MS.

### N- and O-linked glycan release, purification, and permethylation

N- and O-linked glycans were released by following the methods described previously (Shajahan, A., Heiss, C., et al. 2017). N-linked glycans were released from about 80 μg of reduced and alkylated (as mentioned previously) hACE2 sample by treatment with PNGase F at 37°C for 16 h. The released N-glycans were isolated by passing the digest through a C18 SPE cartridge with a 5% acetic acid solution (3 mL) and dried by lyophilization.

The remaining de-N-glycosylated hACE2 protein, still containing O-glycans was then eluted from the column using 80% aqueous acetonitrile with 0.1% formic acid (3 mL). The O-glycans were released from the peptide backbone by reductive *β*-elimination (Shajahan, A., Heiss, C., et al. 2017). The eluted hACE2 was treated with a solution of 19 mg/500 μL of sodium borohydride in a solution of 50 mM sodium hydroxide. The solution was heated to 45 °C for 16 h and neutralized with a solution of 10% acetic acid. The sample was desalted on a hand-packed ion exchange resin (DOWEX H^+^) by eluting with 5% acetic acid and dried by lyophilization. The borates were removed by the addition of a solution of methanol:acetic acid (9:1) and evaporation under a steam of nitrogen.

The released N- and O-linked glycans were then permethylated using methods described elsewhere (Shajahan, A., Heiss, C., et al. 2017).

### Data acquisition of protein digest samples using nano-LC-MS/MS

The glycoprotein digests were analyzed on an Orbitrap Fusion Tribrid mass spectrometer equipped with a nanospray ion source and connected to a Ultimate 3000 RSLCnano nano-LC system (Thermo Fisher, Waltham, MA). A pre-packed nano-LC column (Cat. No. 164568, Thermo Fisher, Waltham, MA) of 15 cm length with 75 μm internal diameter (id), filled with 3-μm C18 material (reverse phase) was used for chromatographic separation of samples. The precursor ion scan was acquired at 120,000 resolution in the Orbitrap analyzer, and precursors at a time frame of 3 s were selected for subsequent MS/MS fragmentation in the Orbitrap analyzer at 15,000 resolution. The LC-MS/MS runs of each digest were conducted for 180 min in order to separate the glycopeptides. The threshold for triggering an MS/MS event was set to 1000 counts, and monoisotopic precursor selection was enabled. MS/MS fragmentation was conducted with stepped HCD (higher-energy collisional dissociation) product triggered CID (collision-induced dissociation) (HCDpdCID) program. For product triggering, the HexNAc fragment m/z 204.087 detected within 10 ppm was chosen as a criteria to trigger the fragment-dependent CID acquisition. Charge state screening was enabled, and precursors with unknown charge state or a charge state of +1 were excluded (positive ion mode). Dynamic exclusion was also enabled (exclusion duration of 30 s).

### Data analysis of glycoproteins

The LC-MS/MS spectra of the combined tryptic /chymotryptic digest of hACE2 were searched against the FASTA sequence of hACE2 (UniProt ID: Q9BYF1, modified according to expressed protein) using the Byonic software by choosing peptide cleavage sites ‘RKFWYL’ for trypsin-chymotrypsin digestion and ‘EFWYL’ for GluC-chymotrypsin digestion (semi-specific cleavage option enabled). Precursor tolerance of 5 ppm and fragment tolerance of 10 ppm was chosen for the search with three missed cleavage option enabled. Oxidation of methionine, deamidation of asparagine and glutamine, and possible common human N-glycans (common human 182 N-glycans) and O-glycan masses (common human six O-glycans) were used as variable modifications. The bad spectra were skipped, and 1 % FDR was enabled while choosing manual score cut-off value of zero. The LC-MS/MS spectra were also analyzed manually for the glycopeptides with the support of the Thermo Fisher Xcalibur 4.2 software. The HCDpdCID MS^2^ spectra of glycopeptides were evaluated for the glycan neutral loss pattern, oxonium ions and glycopeptide fragmentations to assign the sequence and the presence of glycans in the glycopeptides. The relative intensities of glycans at each site were calculated by evaluating the area under the curve of each extracted glycopeptide peak (after deconvolution using Xtract module of Xcalibur 4.2 Qual Browser) on the LC-MS chromatogram.

### N- and O-linked glycomic profiling by MALDI-MS

The permethylated N- and O-glycans were dissolved in 20 μL of methanol. A 0.5-μL portion of sample was mixed with an equal volume of DHB matrix solution (10 mg/mL in 1:1 methanol-water) and spotted on to a MALDI plate. MALDI-MS spectra were acquired in positive ion and reflector mode using an AB Sciex 5800 MALDI-TOF-TOF mass spectrometer.

### N- and O-linked glycomic profiling by DI-ESI-MS^n^

A solution of the permethylated N-glycans from hACE2 was diluted into a solution of 1 mM sodium hydroxide/50% MeOH and directly infused (0.5 μL/min) into an Orbitrap Fusion Tribrid mass spectrometer equipped with a nanospray ion source. An automated program was used to collect a full MS then MS^2^ of the highest-intensity peaks. The top 300 peaks collected from a m/z range of 800-2000 were fragmented by CID, with a dynamic exclusion of 60 s. The total run time was 20 min. The full MS was collected in the Orbitrap at a resolution of 120,000, while the MS^2^ spectra were collected in the Orbitrap at a resolution of 60,000. A similar automated program was used for the collection of O-glycan data with a m/z range adjusted to 600-1600.

Original glycoform assignments were made based on full-mass molecular weight. Additional structural details were determined by MS^n^ and modeling with the GlycoWorkbench 2 software.

### Sialic Acid Linkage Analysis by IT-MS spectrometry including MS^n^ fragmentation

A solution of the permethylated N-glycans from hACE2 was diluted into a solution of 1 mM lithium carbonate/50% MeOH and directly infused (0.5 μL/min) into an Orbitrap Fusion Tribrid mass spectrometer equipped with a nanospray ion source. The sialic acid-containing N-glycans (determined by MALDI-TOF-MS and ESI-MS^n^ experiments) were probed with MS^n^ analysis described previously (Anthony, R.M., Nimmerjahn, F., et al. 2008, Lin, N., Mascarenhas, J., et al. 2015, Shajahan, A., Supekar, N.T., et al. 2017). Isolation was conducted in the quadrupole, while detection was conducted in the IT. The isolation width of each fragmentation was 2 mass units, and the maximum injection time was 100 ms. More than 300 spectra were collected for each glycoform, which were then spectrally averaged.

### Determination of bisecting GlcNAc by IT-MS spectrometry including MS^n^ fragmentation

A solution of the permethylated N-glycans from hACE2 was diluted into a solution of 1 mM sodium hydroxide/50% MeOH and directly infused (0.5 μL/min) into an Orbitrap Fusion Tribrid mass spectrometer equipped with a nanospray ion source. The neutral complex-type N-glycans (determined by MALDI-TOF-MS and ESI-MS^n^ experiments) were probed with MS^n^ analysis described previously (Ashline, D.J., Duk, M., et al. 2015). Because of the extra fragmentation steps, sialylated N-glycans were not probed since they yielded too low of a signal. Isolation was conducted in the quadrupole while detection was conducted in the IT. The isolation width of each fragmentation was 2 mass units, and the maximum injection time was 100 ms. More than 300 spectra were collected for each glycoform, which were then spectrally averaged.

## Data Availability

The raw data files and search results can be accessed from glycopost repository - https://glycopost.glycosmos.org/preview/15928494275f039184c7229; pin: 5771.

## Acknowledgements

Financial support from the US National Institutes of Health (S10OD018530) is gratefully acknowledged. This work was also supported in part by the U.S. Department of Energy, Office of Science, Basic Energy Sciences, under Award DE-SC0015662 to DOE - Center for Plant and Microbial Complex Carbohydrates at the Complex Carbohydrate Research Center, USA. We thank Rupali Mahadik (CCRC, UGA) for help with the retrieval of hACE2 data from glygen.org. We also thank Daniel S. Rouhani for producing the graphical abstract.

## Abbreviations

S: Spike
RBD: receptor binding domain
COVID-19: coronavirus disease
hACE2: human angiotensinconverting enzyme 2
HAs: hemagglutinins
HEK: human embryonic kidney
DTT: dithiothreitol
IAA: iodoacetamide
PNGase F: Peptide-N-Glycosidase F
SPE: solid phase extraction
ACN: acetonitrile
HCD: Higher-energy Collisional Dissociation
pd: product triggered
CID: Collision-Induced Dissociation
IT-MS: ion trap-mass spectrometry.

## Contributions

P.A. and A.S. conceived of the paper; A.S. performed glycoproteomics and glycomics sample processing, S.A.H. conducted glycomics data acquisition, A.S., S.A.H., N.S. and A.G. performed data analysis; everyone contributed toward writing the paper; P.A. and C.H. monitored the project.

## Supplemental material

Annotated glycopeptide MS/MS spectra (S1 – S8), N- and O- glycan MALDI-MS data (S9, S10), and ESI-MS/MS data (S11) are incorporated as a separate supplemental material file. Supplemental Tables 1 and 2 are also included as separate files.

