## Supplementary information for "Comprehensive characterization of N- and O- glycosylation of SARS-CoV-2 human receptor angiotensin converting enzyme 2"

**Table of Contents**


| Description | Page No. |
|---|---|
| Supp. Figure S1: Spectra of intact N-glycopeptides with assigned N-glycans at N53 | S2-S6 |
| Supp. Figure S2: Spectra of intact N-glycopeptides with assigned N-glycans at N90 | S6-S9 |
| Supp. Figure S3: Spectra of intact N-glycopeptides with assigned N-glycans at N103 | S9-S11 |
| Supp. Figure S4: Spectra of intact N-glycopeptides with assigned N-glycans at N322 | S11-S16 |
| Supp. Figure S5: Spectra of intact N-glycopeptides with assigned N-glycans at N432 | S16-21 |
| Supp. Figure S6: Spectra of intact N-glycopeptides with assigned N-glycans at N546 | S21-S24 |
| Supp. Figure S7: Spectra of intact N-glycopeptides with assigned N-glycans at N690 | S24-S31 |
| Supp. Figure S8: Spectra of intact O-glycopeptides with assigned O-glycans at T730 | S32 |
| Supp. Figure S9: MALDI-MS spectrum showing the permethylated N-glycans released from hACE2 by PNGase F. | S33 |
| Supp. Figure S10: MALDI-MS spectrum showing the permethylated O-glycans released from hACE2 by β-elimination. | S33 |
| Supp. Figure S11: ESI-MSn CID spectra based sequencing of N-glycans. | S34 |

Glycopeptide Analysis:

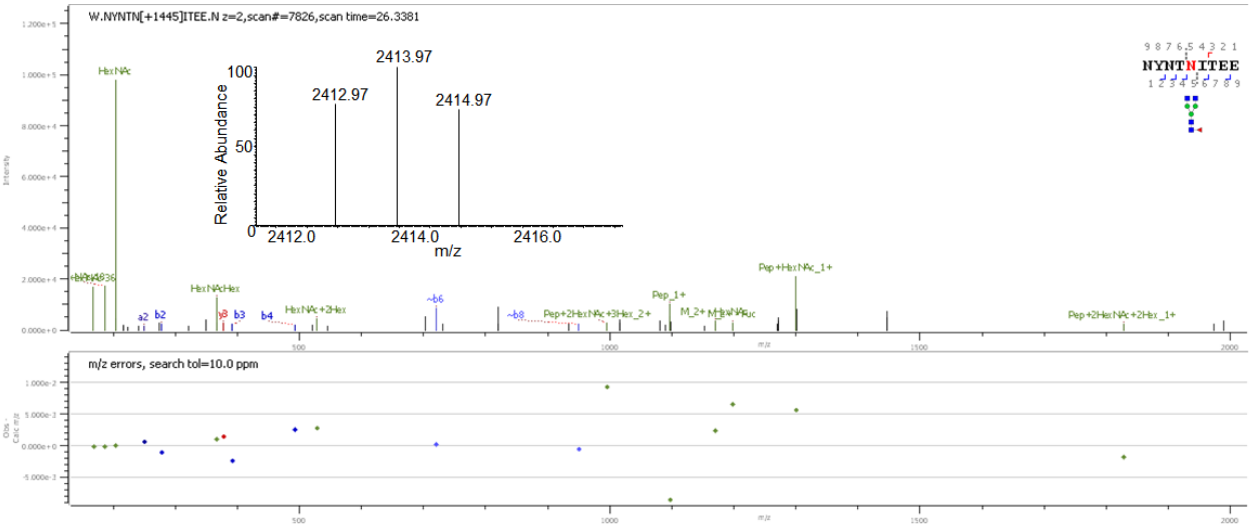


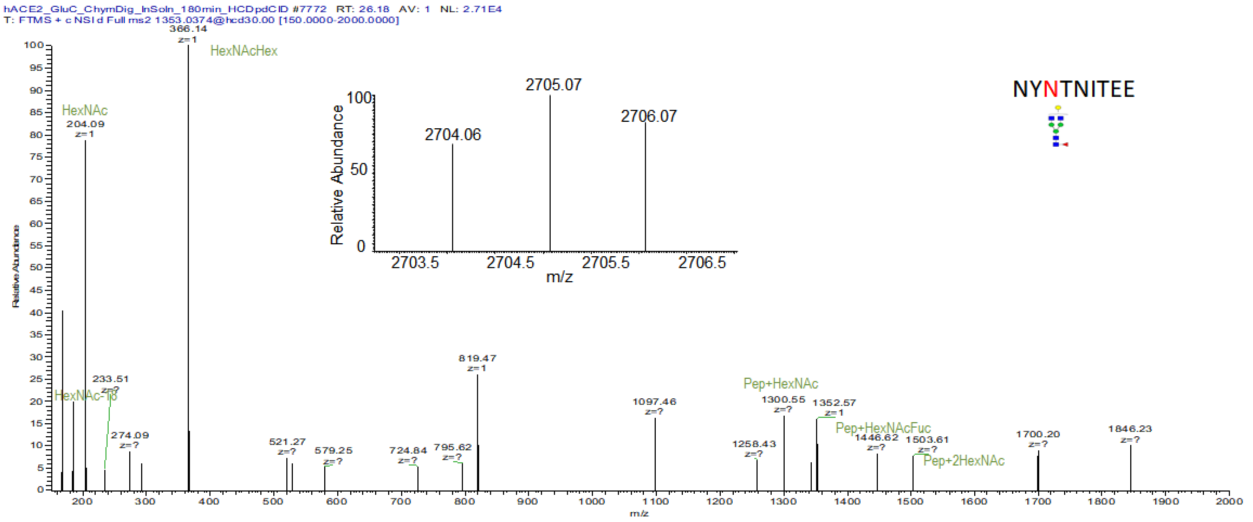


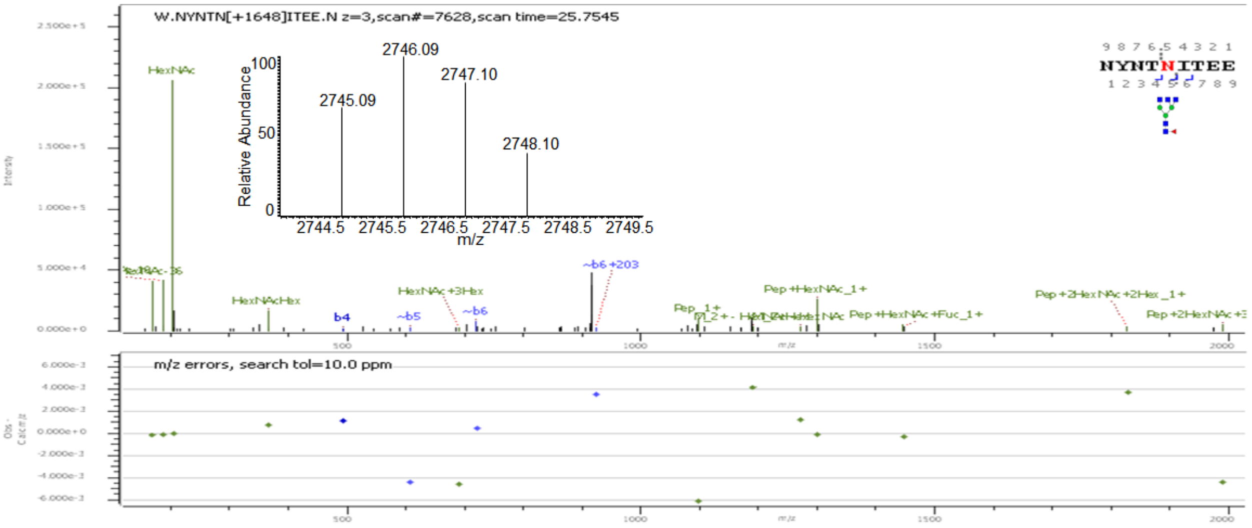

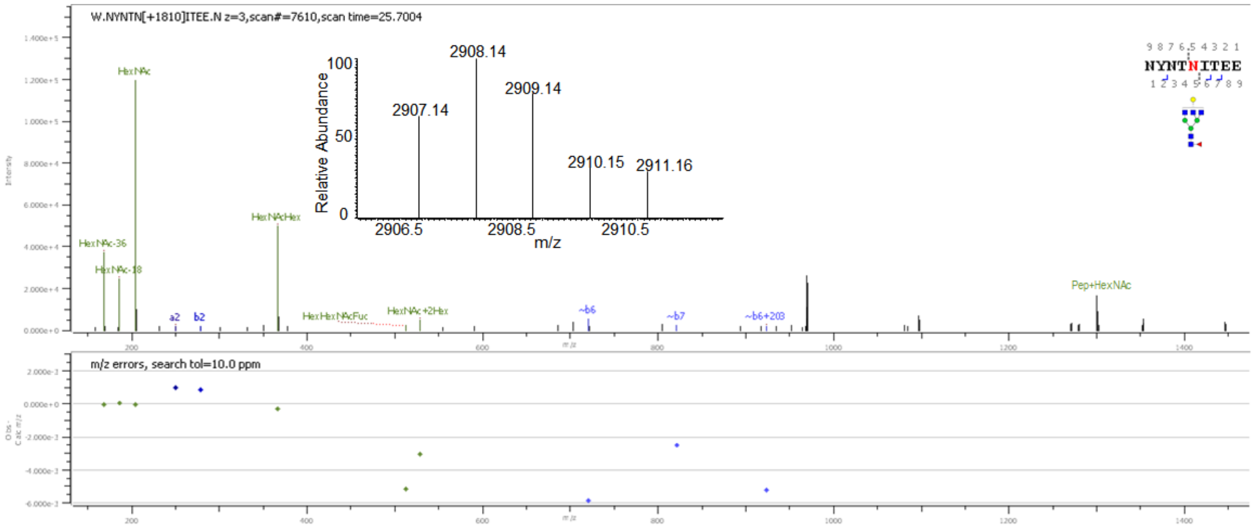
W.NYNTN[+1810]ITEE.N z=3,scan#=7610,scan time=25.7004
HexNAc
HexNAcHex
HexNAc-36
HexNAc-18
a2
b2
HexHexNAcFuc
HexNAc+2Hex
~b6
~b7
~b6+203
Pep+HexNAc
2908.14
2909.14
2907.14
2910.15
2911.16
Relative Abundance
m/z
2906.5
2908.5
2910.5
NYNTNITEE
m/z errors, search tol=10.0 ppm

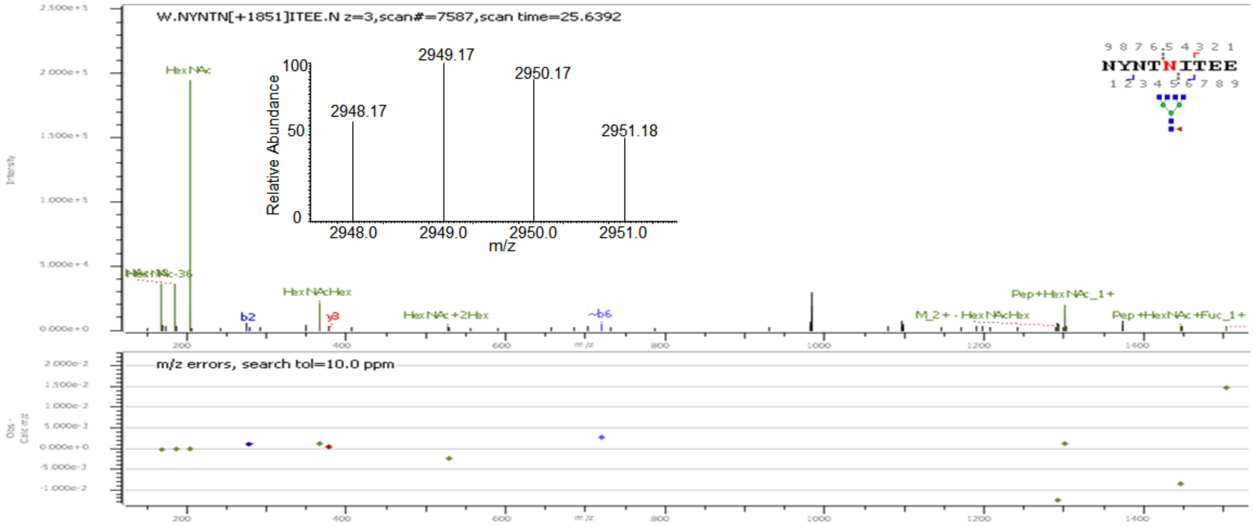
W.NYNTN[+1851]ITEE.N z=3,scan#=7587,scan time=25.6392
HexNAc
HexNAcHex
b2
y8
HexNAc+2Hex
~b6
Pep+HexNAc_1+
M_2+ - HexNAcHex
Pep+HexNAc+Fuc_1+
2949.17
2950.17
2948.17
2951.18
Relative Abundance
m/z
2948.0
2949.0
2950.0
2951.0
NYNTNITEE
m/z errors, search tol=10.0 ppm

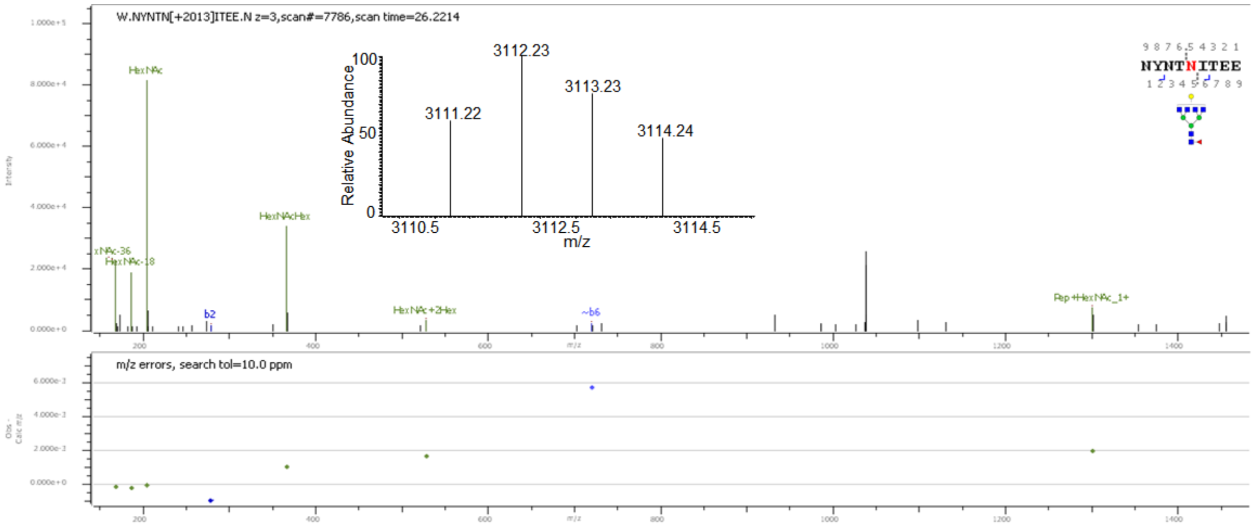
W.NYNTN[+2013]ITEE.N z=3,scan#=7786,scan time=26.2214
HexNAc
HexNAcHex
HexNAc-18
b2
HexNAc+2Hex
~b6
Pep+HexNAc_1+
3112.23
3113.23
3111.22
3114.24
Relative Abundance
m/z
3110.5
3112.5
3114.5
NYNTNITEE
m/z errors, search tol=10.0 ppm

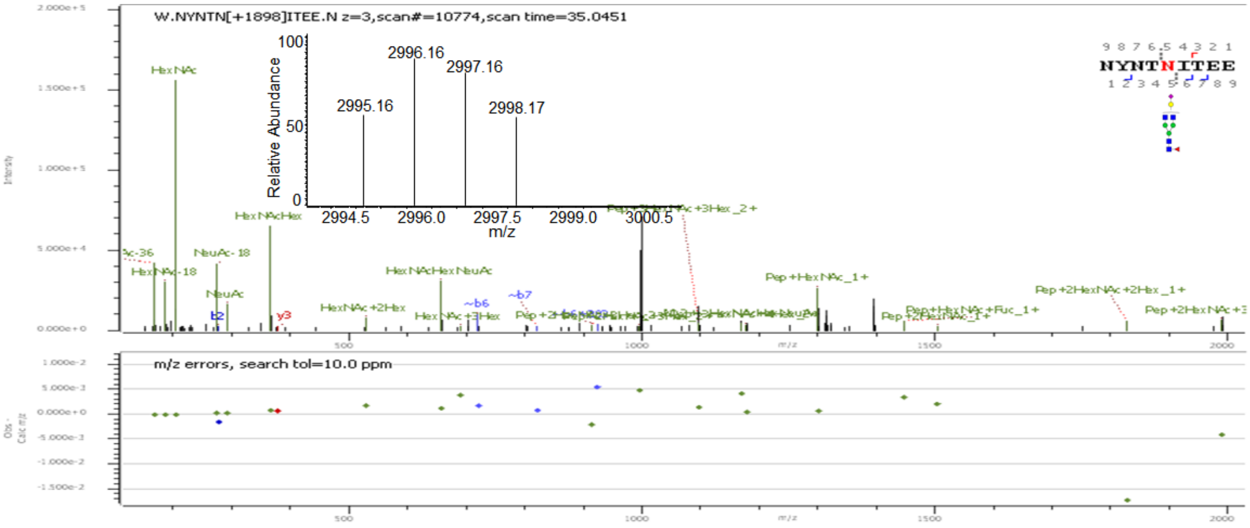
W.NYNTN[+1898]ITEE.N z=3,scan#=10774,scan time=35.0451
Relative Abundance
2995.16
2996.16
2997.16
2998.17
m/z
HexNAc
HexNAcHex
NeuAc-18
NeuAc
HexNAcHexNeuAc
HexNAc+2Hex
Pep+HexNAc_1+
Pep+2HexNAc+2Hex_1+
m/z errors, search tol=10.0 ppm

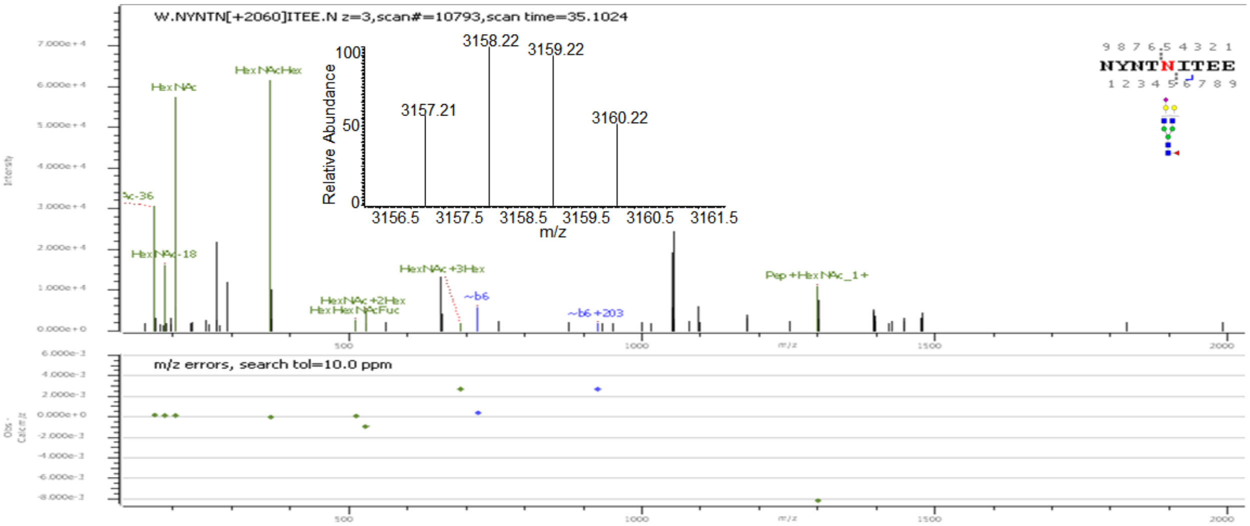
W.NYNTN[+2060]ITEE.N z=3,scan#=10793,scan time=35.1024
Relative Abundance
3157.21
3158.22
3159.22
3160.22
m/z
HexNAc
HexNAcHex
HexNAc+3Hex
HexNAc+2Hex
HexHexNAcFuc
Pep+HexNAc_1+
m/z errors, search tol=10.0 ppm

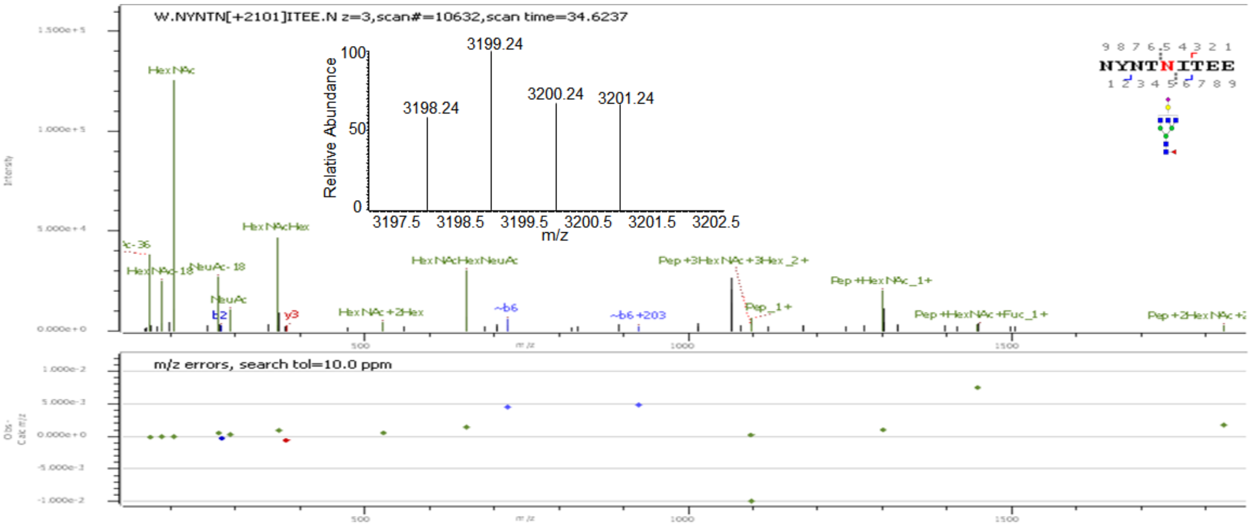
W.NYNTN[+2101]ITEE.N z=3,scan#=10632,scan time=34.6237
Relative Abundance
3198.24
3199.24
3200.24
3201.24
m/z
HexNAc
HexNAcHex
NeuAc-18
NeuAc
HexNAcHexNeuAc
HexNAc+2Hex
Pep+3HexNAc+3Hex_2+
Pep+HexNAc_1+
Pep+HexNAc+Fuc_1+
m/z errors, search tol=10.0 ppm

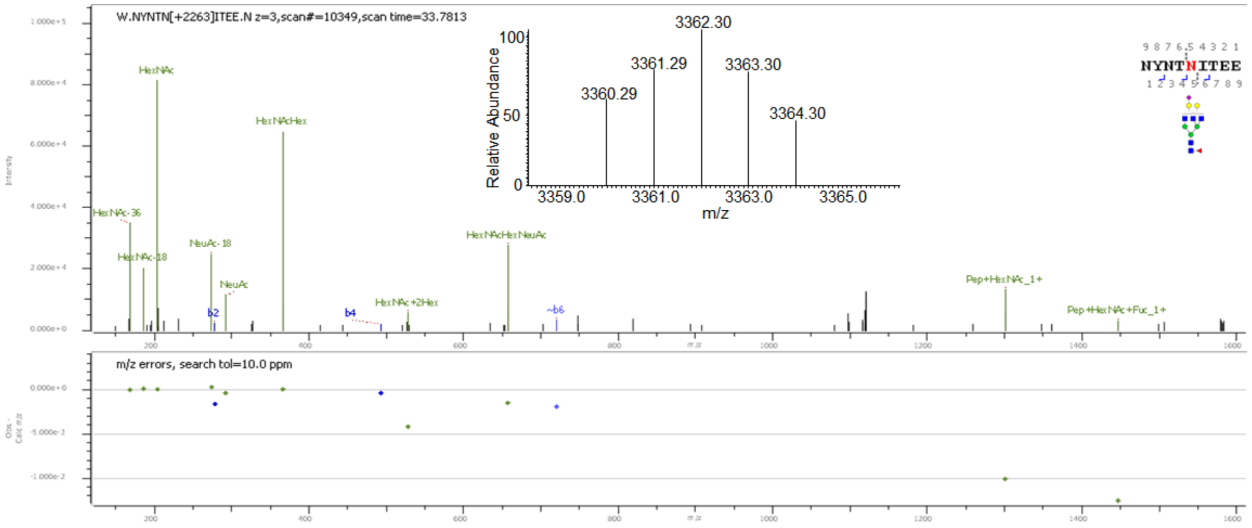
W.NYNTN[+2263]ITEE.N z=3,scan#=10349,scan time=33.7813
Relative Abundance
m/z
3362.30
3361.29
3363.30
3360.29
3364.30
NYNTNITEE
HexNAc
HexNAcHex
HexNAc-36
HexNAc-18
NeuAc-18
NeuAc
HexNAcHexNeuAc
HexNAc+2Hex
b2
b4
~b6
Pep+HexNAc_1+
Pep+HexNAc+Fuc_1+
m/z errors, search tol=10.0 ppm

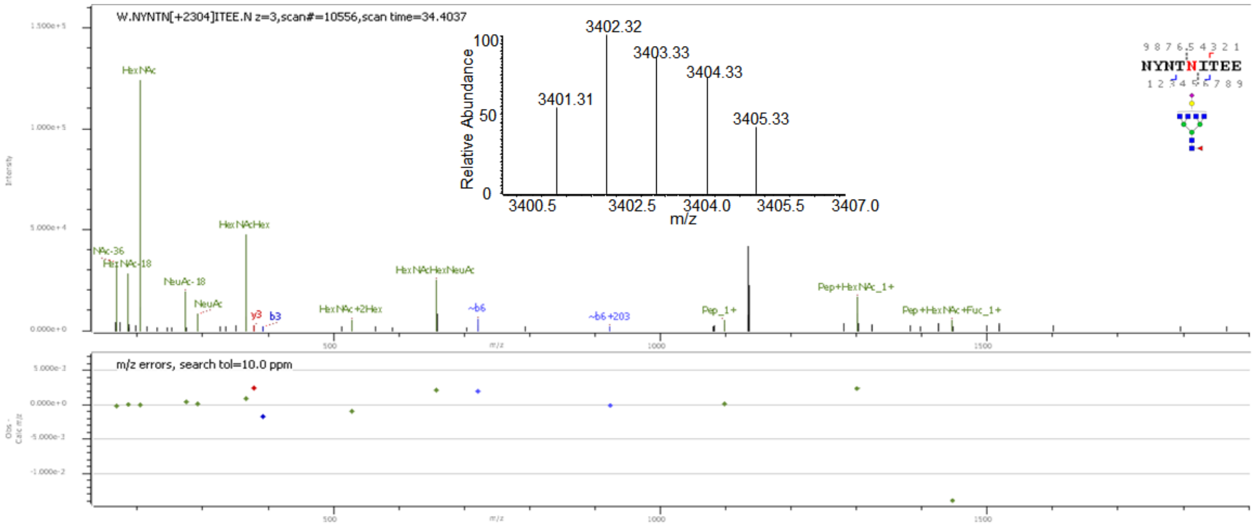
W.NYNTN[+2304]ITEE.N z=3,scan#=10556,scan time=34.4037
Relative Abundance
m/z
3402.32
3403.33
3404.33
3401.31
3405.33
NYNTNITEE
HexNAc
HexNAcHex
HexNAc-18
NeuAc-18
NeuAc
HexNAcHexNeuAc
HexNAc+2Hex
y3
b3
~b6
~b6+203
Pep_1+
Pep+HexNAc_1+
Pep+HexNAc+Fuc_1+
m/z errors, search tol=10.0 ppm

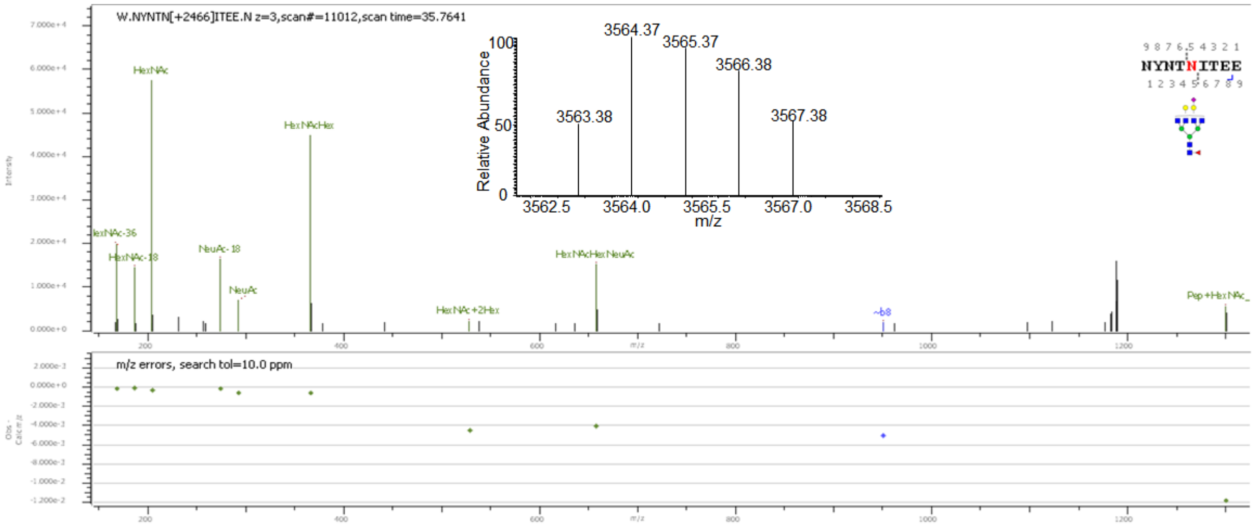
W.NYNTN[+2466]ITEE.N z=3,scan#=11012,scan time=35.7641
Relative Abundance
m/z
3564.37
3565.37
3566.38
3563.38
3567.38
NYNTNITEE
HexNAc
HexNAcHex
HexNAc-18
NeuAc-18
NeuAc
HexNAcHexNeuAc
HexNAc+2Hex
~b8
m/z errors, search tol=10.0 ppm

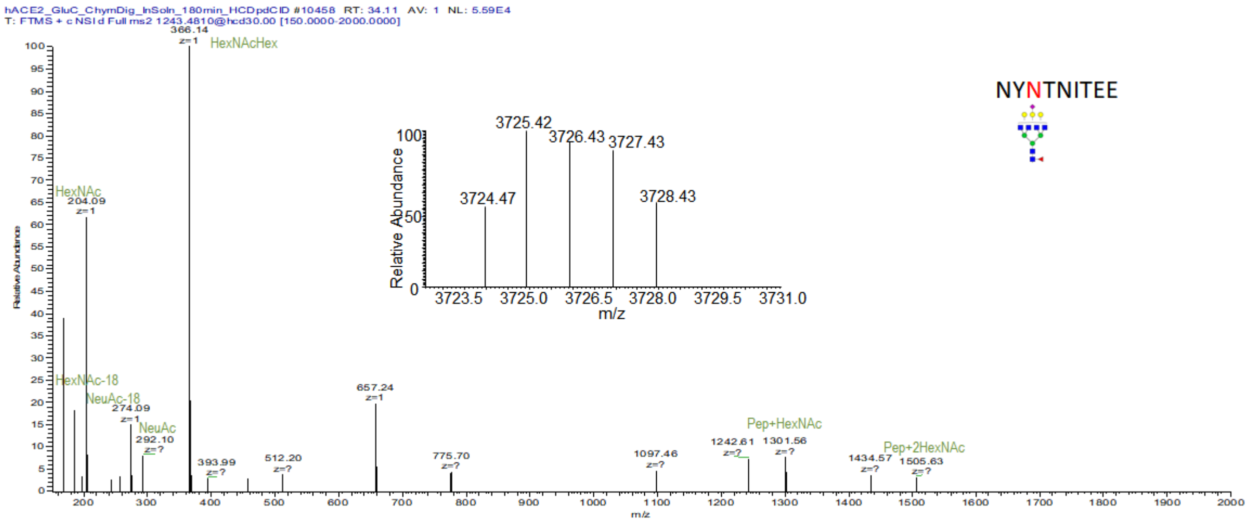


**Supp. Figure S1:** Deconvoluted precursor ion isotopic patterns (insets) and annotated HCD MS/MS spectra of intact N-glycopeptides with assigned N-glycans at N53.

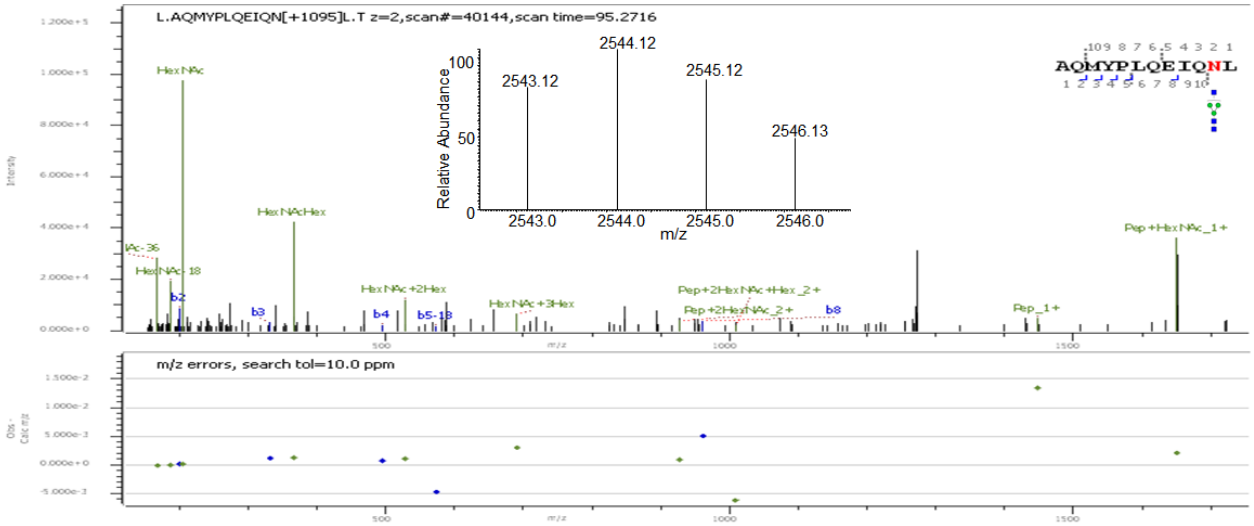


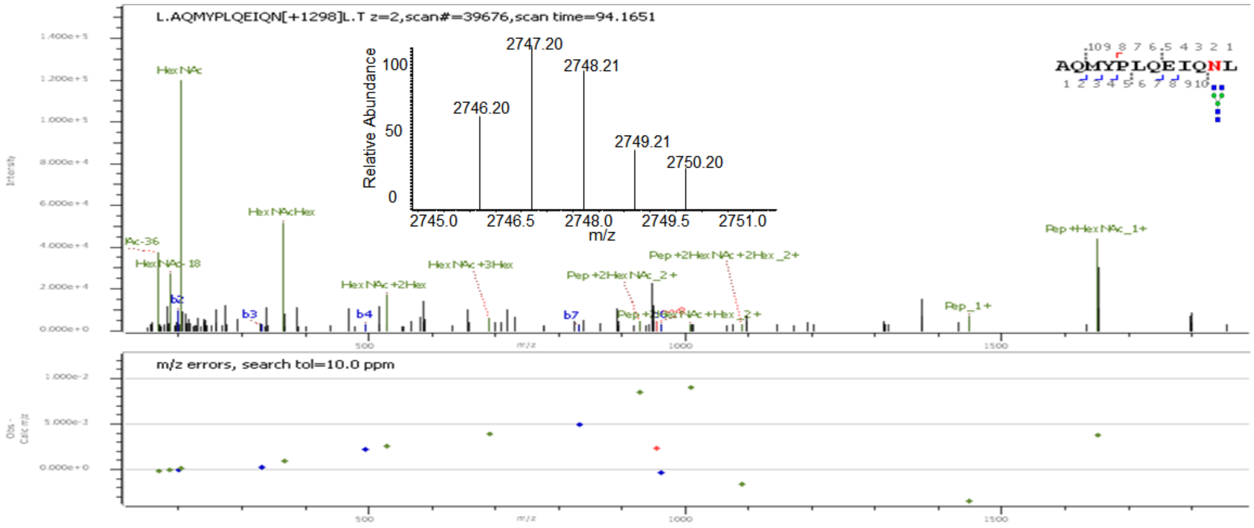

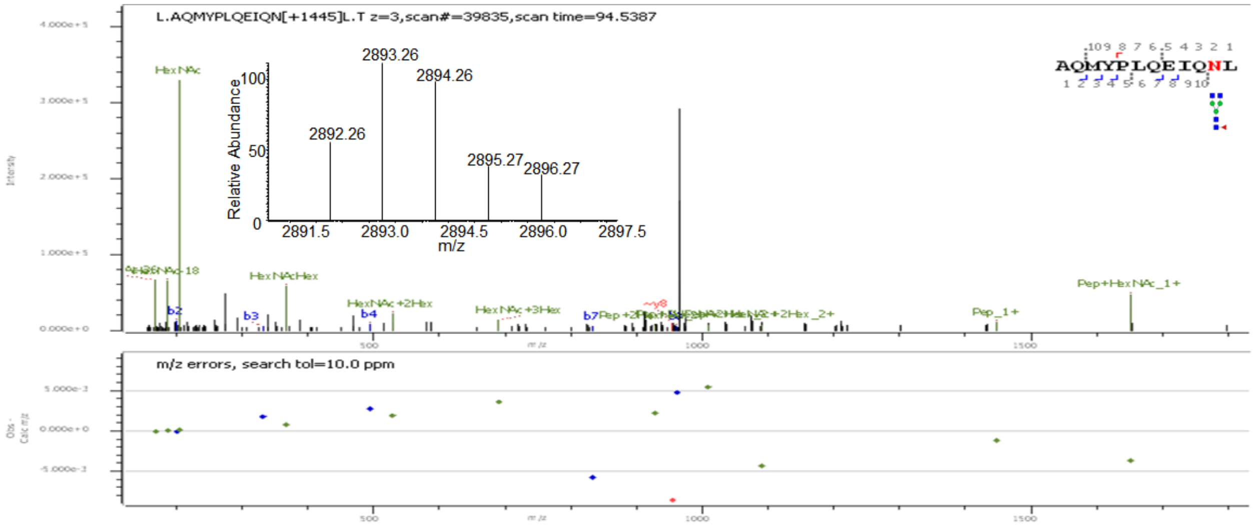
L.AQMYPLQEIQN[+1445]L.T z=3,scan#=39835,scan time=94.5387
AQMYPLQEIQNL
Relative Abundance
2892.26
2893.26
2894.26
2895.27
2896.27
m/z
HexNAc
HexNAcHex
HexNAc+2Hex
HexNAc+3Hex
Pep_1+
Pep+HexNAc_1+
m/z errors, search tol=10.0 ppm

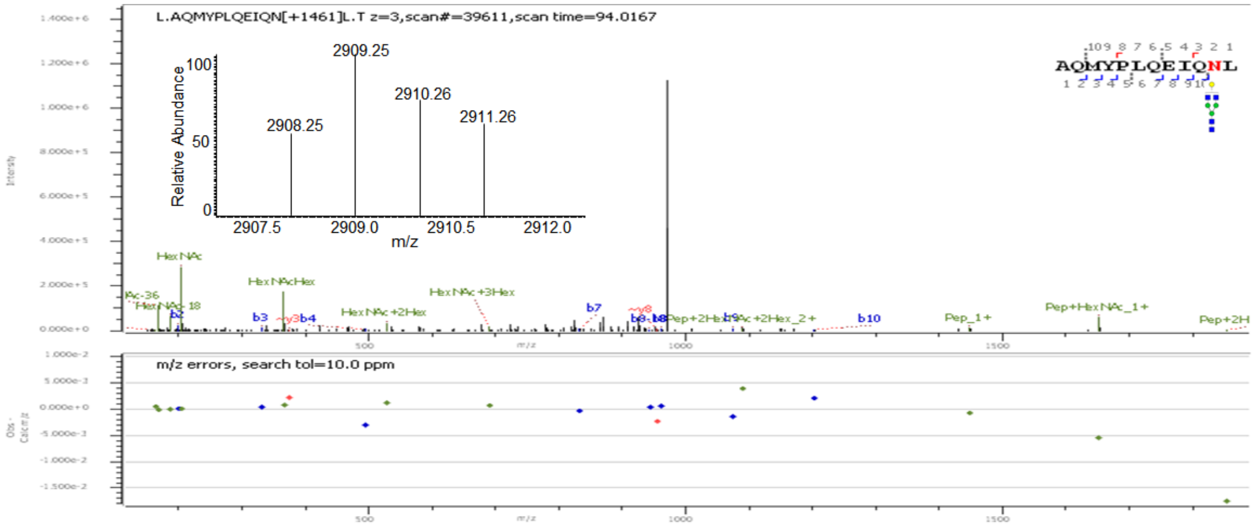
L.AQMYPLQEIQN[+1461]L.T z=3,scan#=39611,scan time=94.0167
AQMYPLQEIQNL
Relative Abundance
2908.25
2909.25
2910.26
2911.26
m/z
HexNAc
HexNAcHex
HexNAc+2Hex
HexNAc+3Hex
Pep_1+
Pep+HexNAc_1+
m/z errors, search tol=10.0 ppm

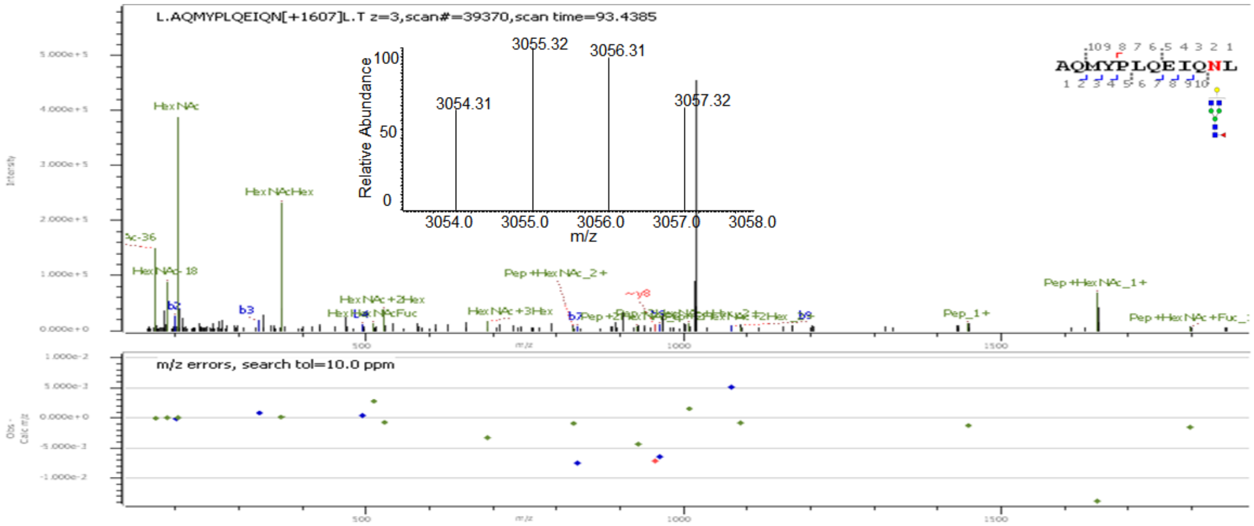
L.AQMYPLQEIQN[+1607]L.T z=3,scan#=39370,scan time=93.4385
AQMYPLQEIQNL
Relative Abundance
3054.31
3055.32
3056.31
3057.32
m/z
HexNAc
HexNAcHex
HexNAc+2Hex
HexNAc+3Hex
Pep+HexNAc_2+
Pep_1+
Pep+HexNAc_1+
m/z errors, search tol=10.0 ppm

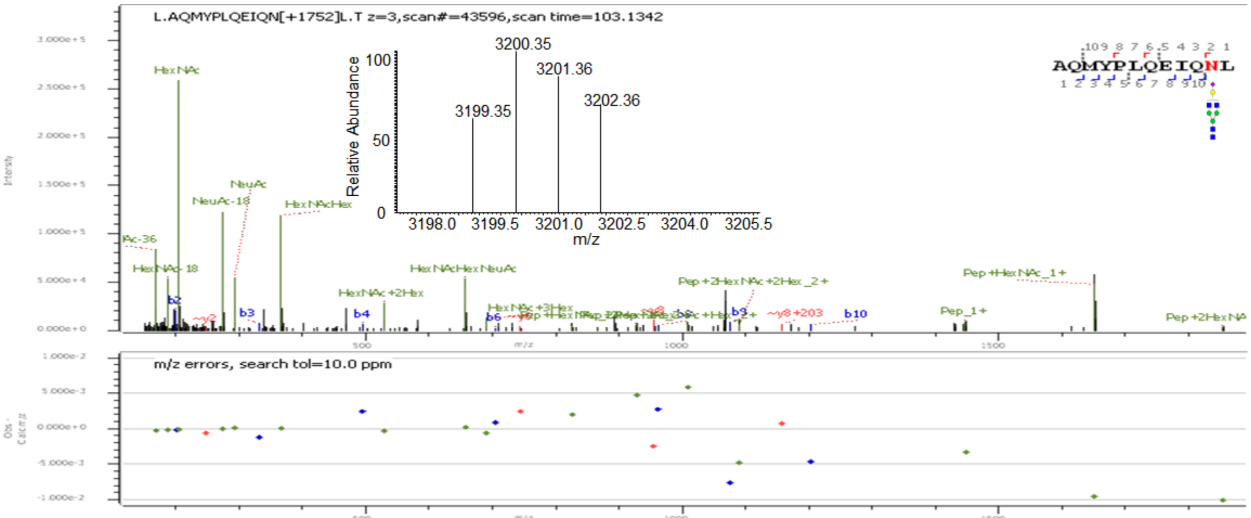
L.AQMYPLQEIQN[+1752]L.T z=3,scan#=43596,scan time=103.1342
3200.35
3201.36
3199.35
3202.36
Relative Abundance
m/z
AQMYPLQEIQNL
m/z errors, search tol=10.0 ppm

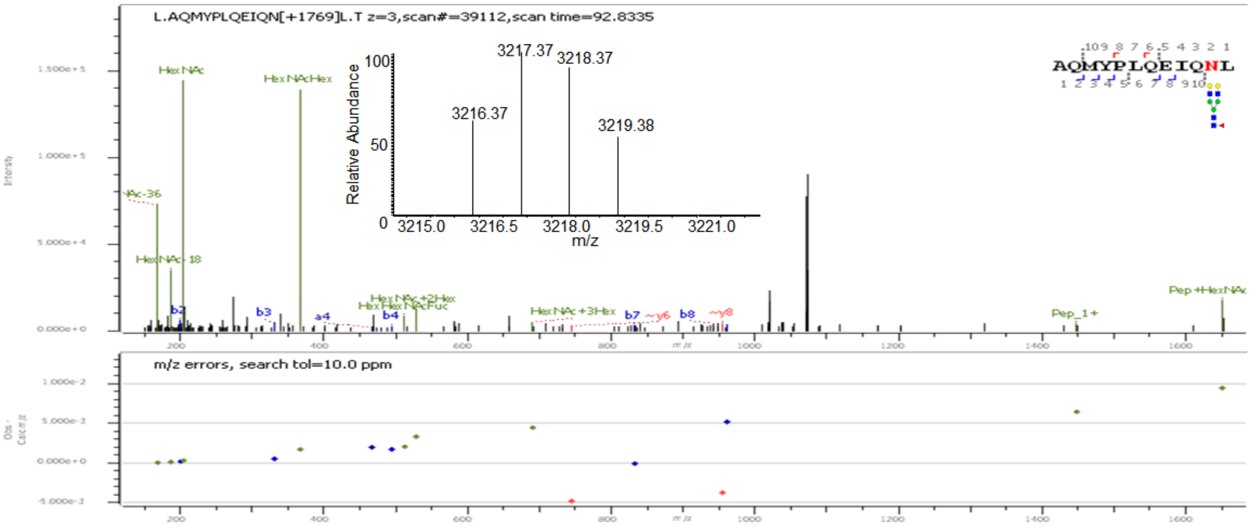
L.AQMYPLQEIQN[+1769]L.T z=3,scan#=39112,scan time=92.8335
3217.37
3218.37
3216.37
3219.38
Relative Abundance
m/z
AQMYPLQEIQNL
m/z errors, search tol=10.0 ppm

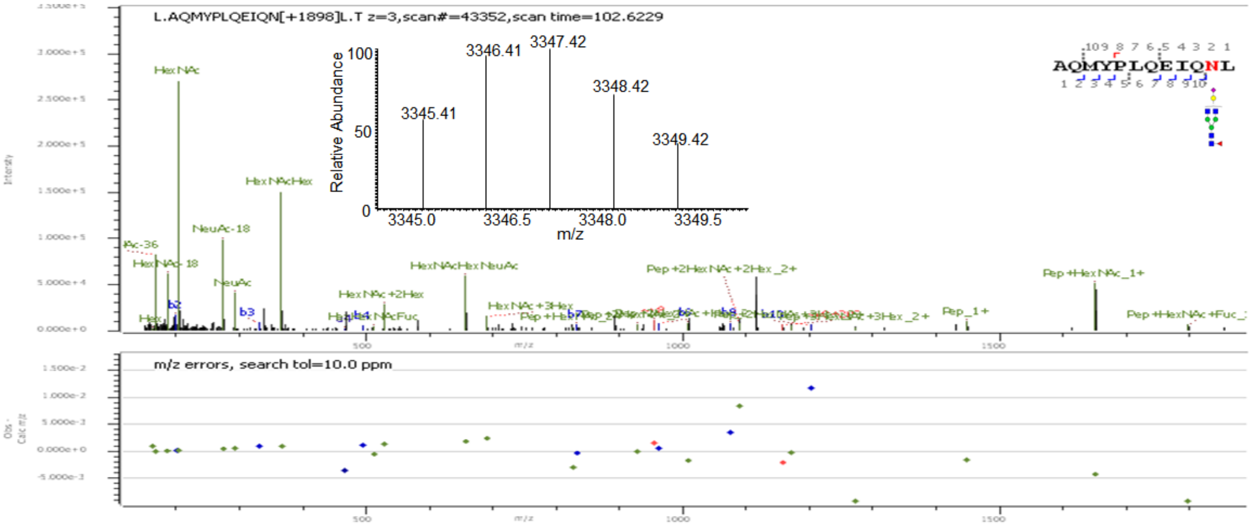
L.AQMYPLQEIQN[+1898]L.T z=3,scan#=43352,scan time=102.6229
3346.41
3347.42
3348.42
3345.41
3349.42
Relative Abundance
m/z
AQMYPLQEIQNL
m/z errors, search tol=10.0 ppm

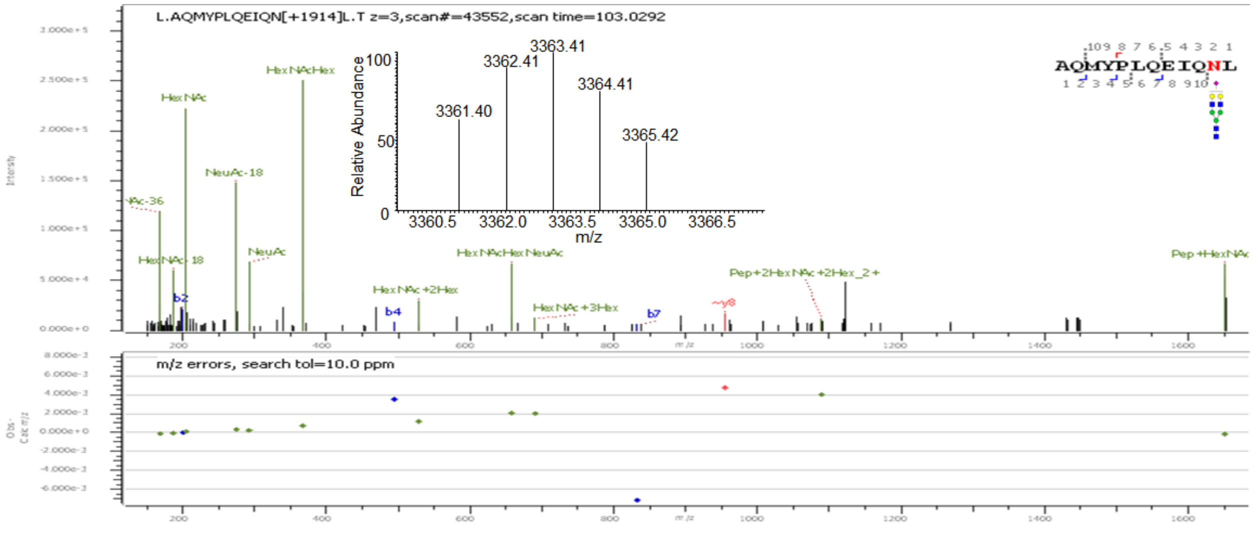


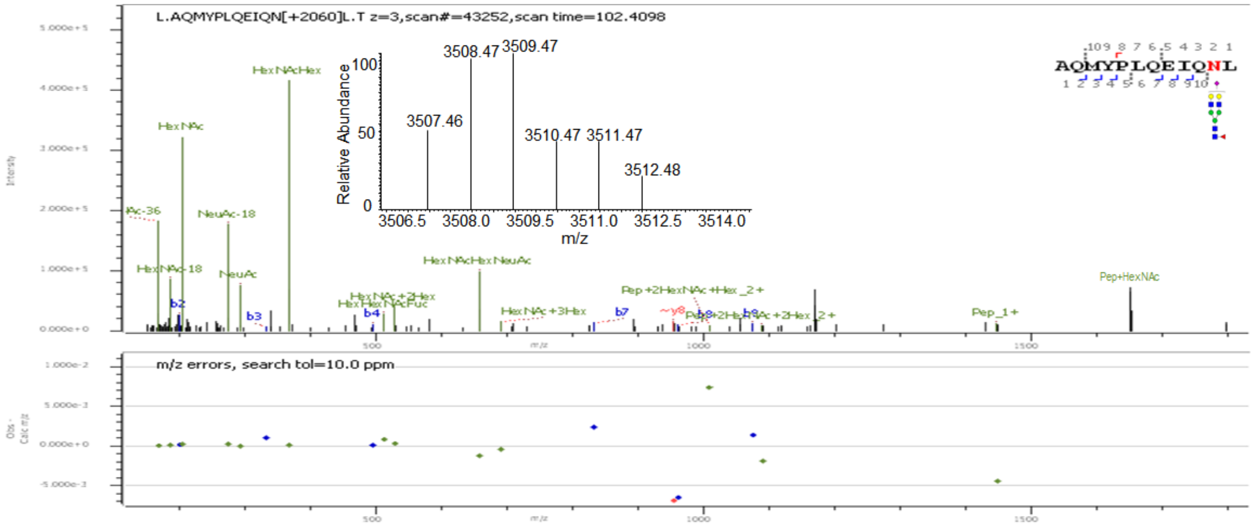


**Supp. Figure S2:** Deconvoluted precursor ion isotopic patterns (insets) and annotated HCD MS/MS spectra of intact N-glycopeptides with assigned N-glycans at N90.

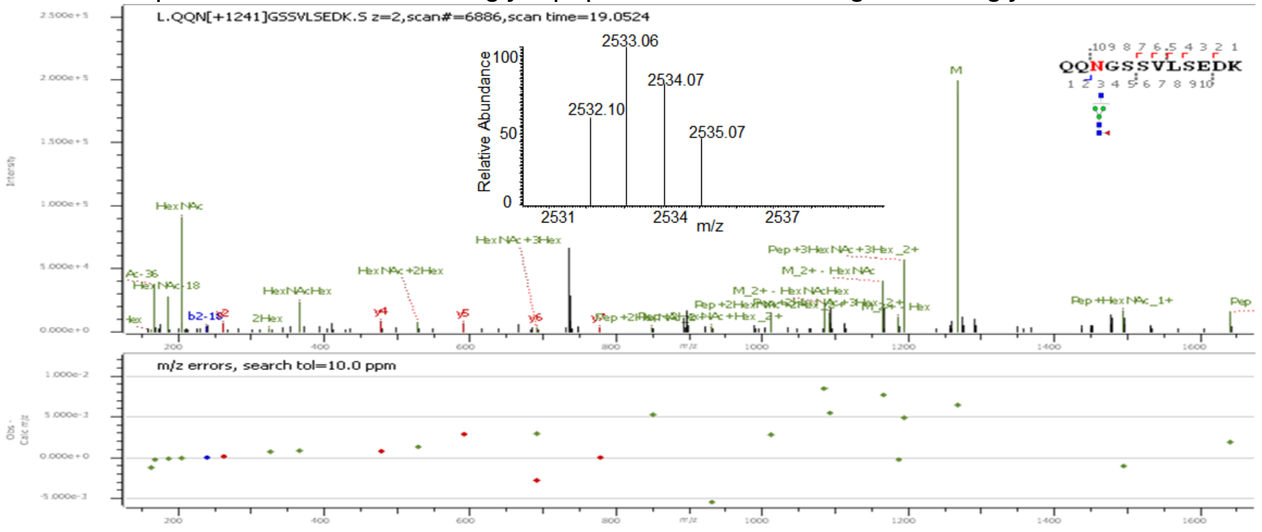

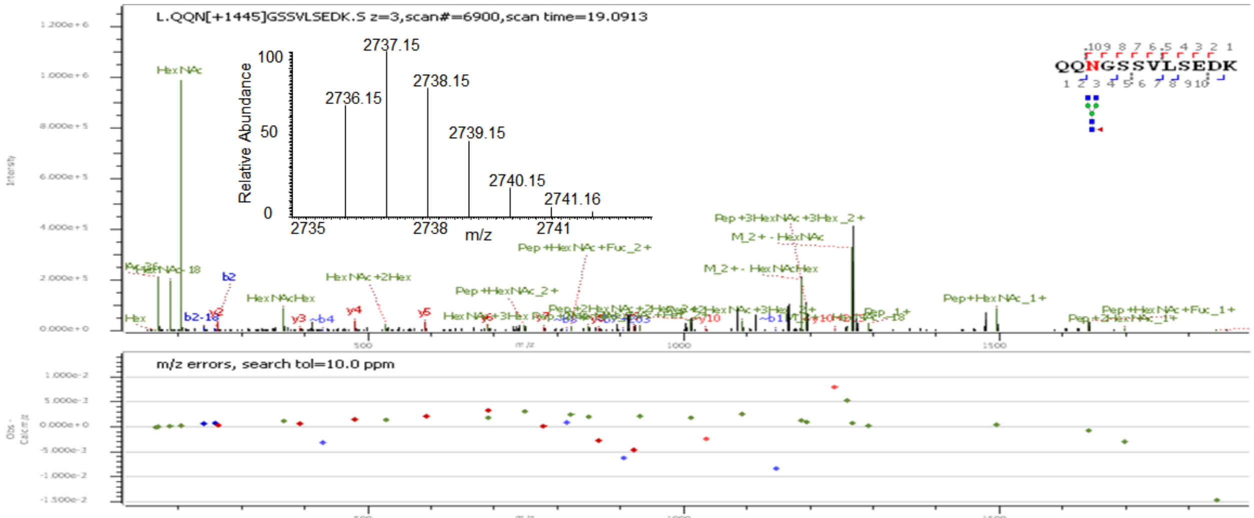
L.QQN[+1445]GSSVLSEDK.S z=3,scan#=6900,scan time=19.0913
QQNGSSVLSEDK
2737.15
2736.15
2738.15
2739.15
2740.15
2741.16
Relative Abundance
m/z
m/z errors, search tol=10.0 ppm

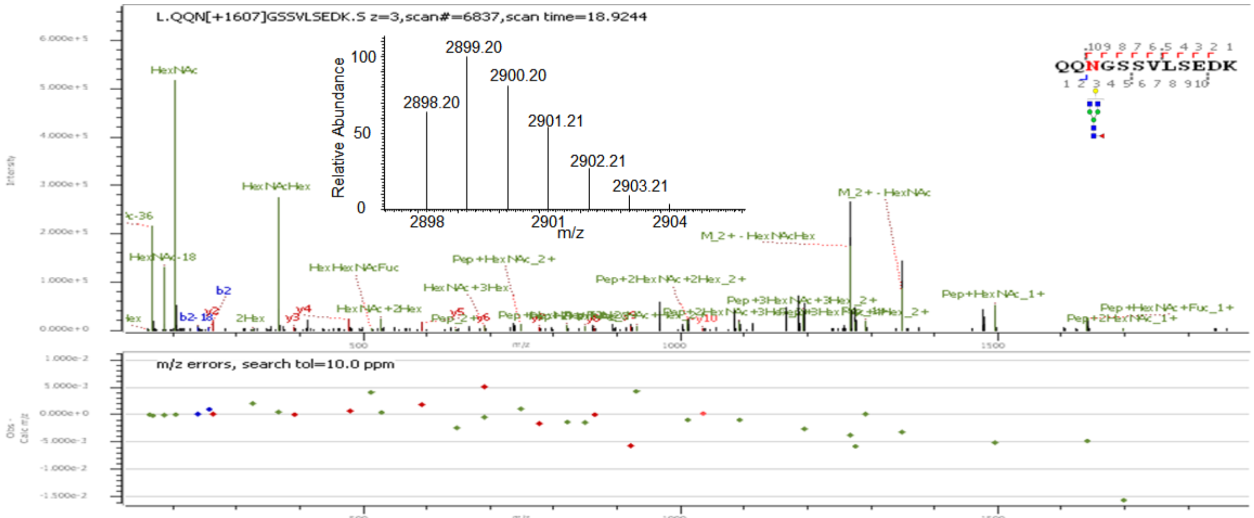
L.QQN[+1607]GSSVLSEDK.S z=3,scan#=6837,scan time=18.9244
QQNGSSVLSEDK
2899.20
2898.20
2900.20
2901.21
2902.21
2903.21
Relative Abundance
m/z
m/z errors, search tol=10.0 ppm

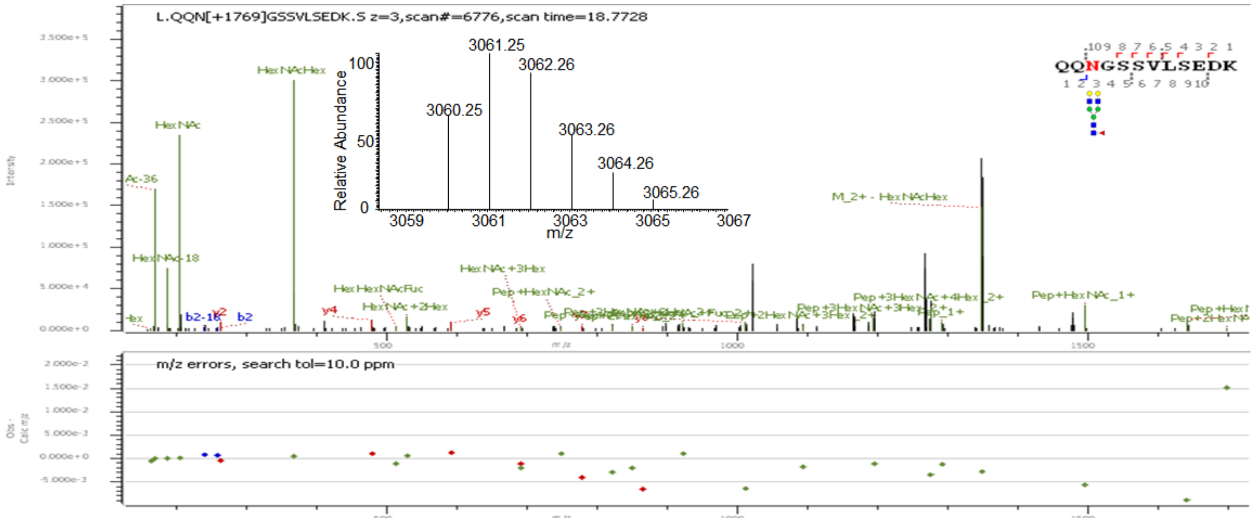
L.QQN[+1769]GSSVLSEDK.S z=3,scan#=6776,scan time=18.7728
QQNGSSVLSEDK
3061.25
3062.26
3060.25
3063.26
3064.26
3065.26
Relative Abundance
m/z
m/z errors, search tol=10.0 ppm

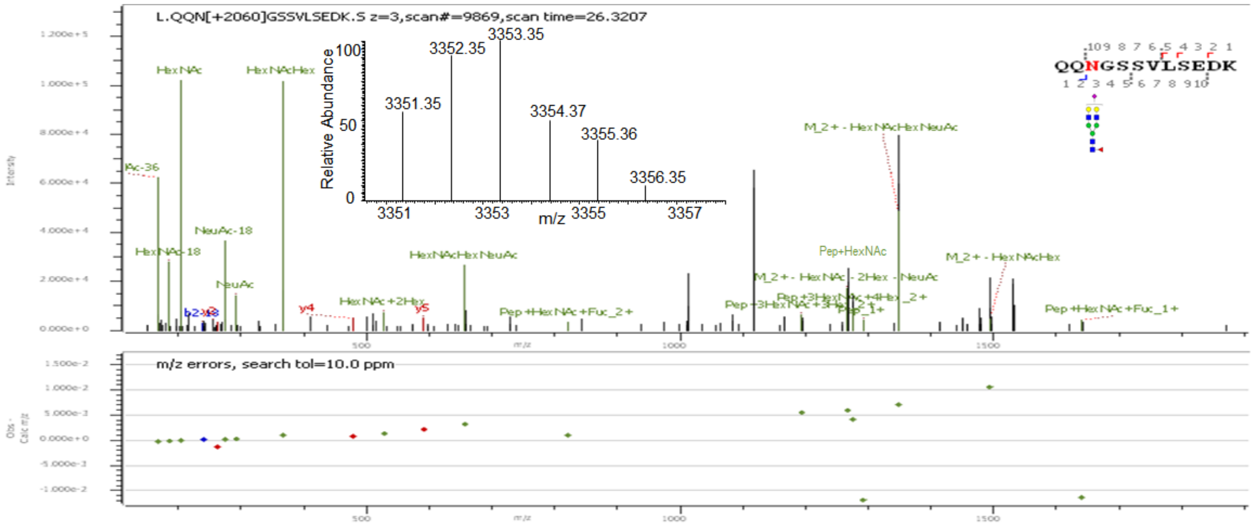


**Supp. Figure S3:** Deconvoluted precursor ion isotopic patterns (insets) and annotated HCD MS/MS spectra of intact N-glycopeptides with assigned N-glycans at N103.

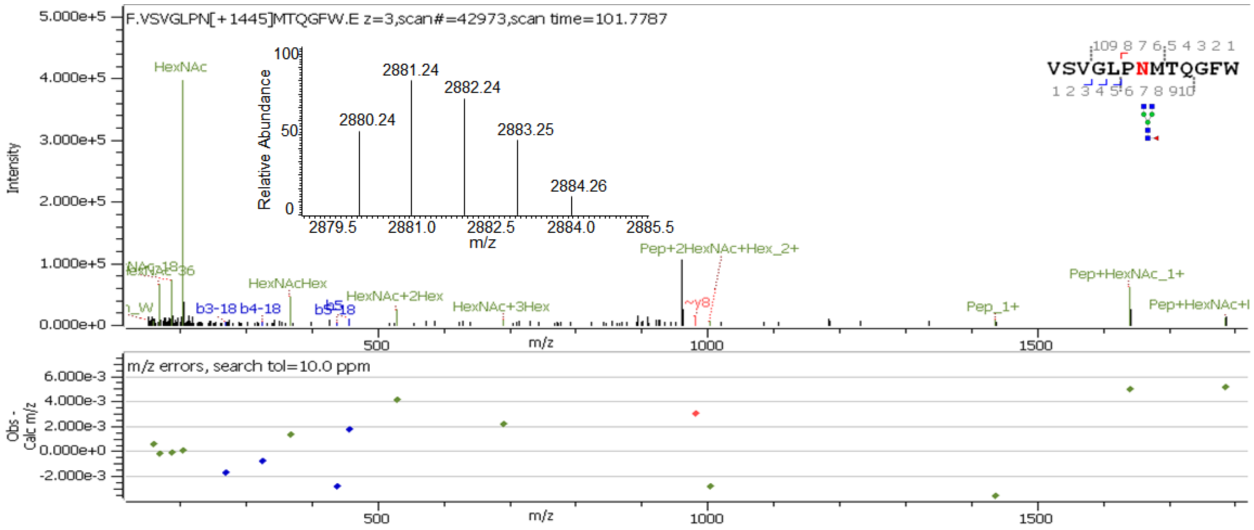


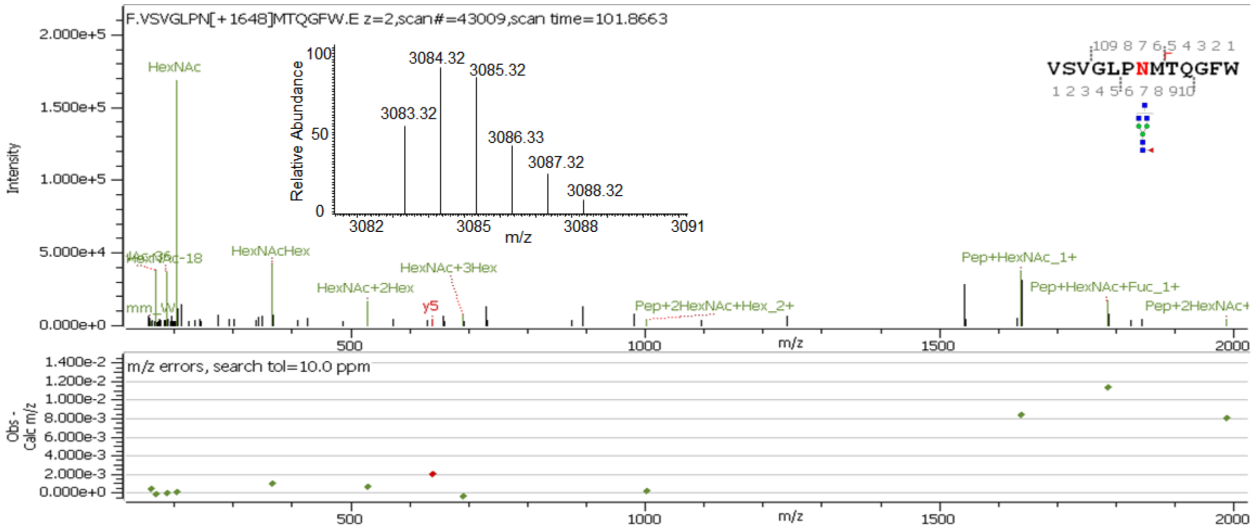

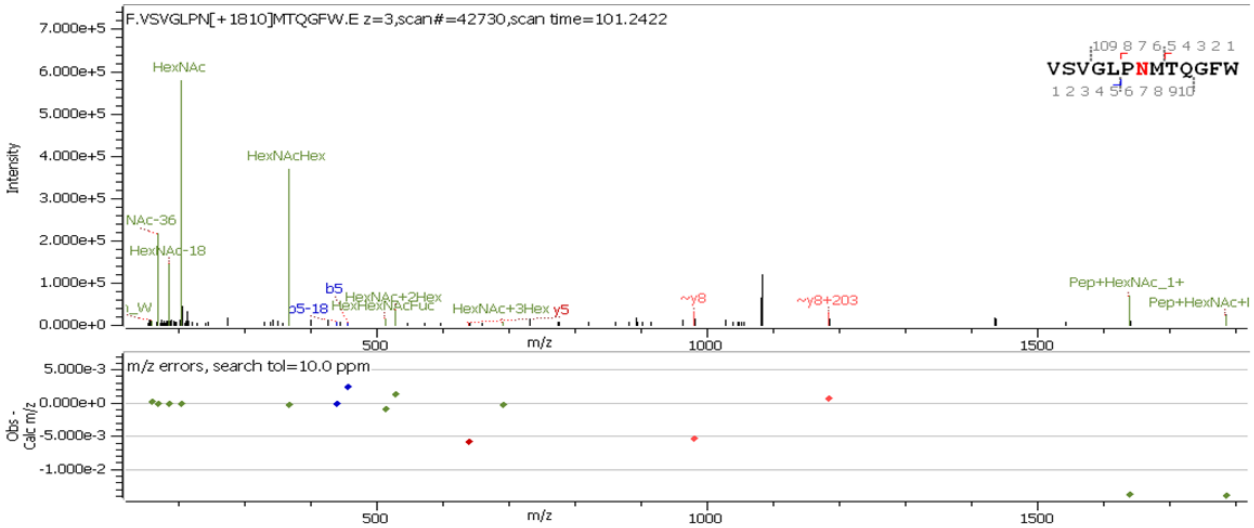
F.VSVGLPN[+1810]MTQGFW.E z=3,scan#=42730,scan time=101.2422
VSVGLPNMTQGFW
Intensity
HexNAc
HexNAcHex
HexNAc-18
b5
b5-18
HexNAc+2Hex
HexNAc+3Hex
y5
~y8
~y8+203
Pep+HexNAc_1+
m/z
m/z errors, search tol=10.0 ppm
Obs - Calc m/z

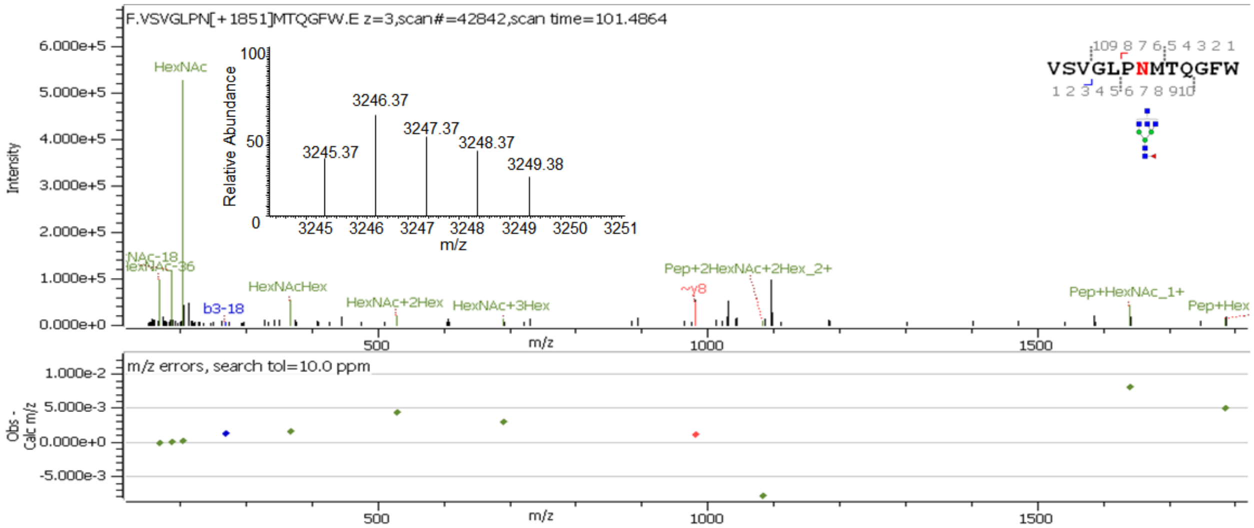
F.VSVGLPN[+1851]MTQGFW.E z=3,scan#=42842,scan time=101.4864
VSVGLPNMTQGFW
Intensity
Relative Abundance
3245.37
3246.37
3247.37
3248.37
3249.38
m/z
HexNAc
HexNAcHex
HexNAc+2Hex
HexNAc+3Hex
b3-18
Pep+2HexNAc+2Hex_2+
~y8
Pep+HexNAc_1+
m/z errors, search tol=10.0 ppm
Obs - Calc m/z

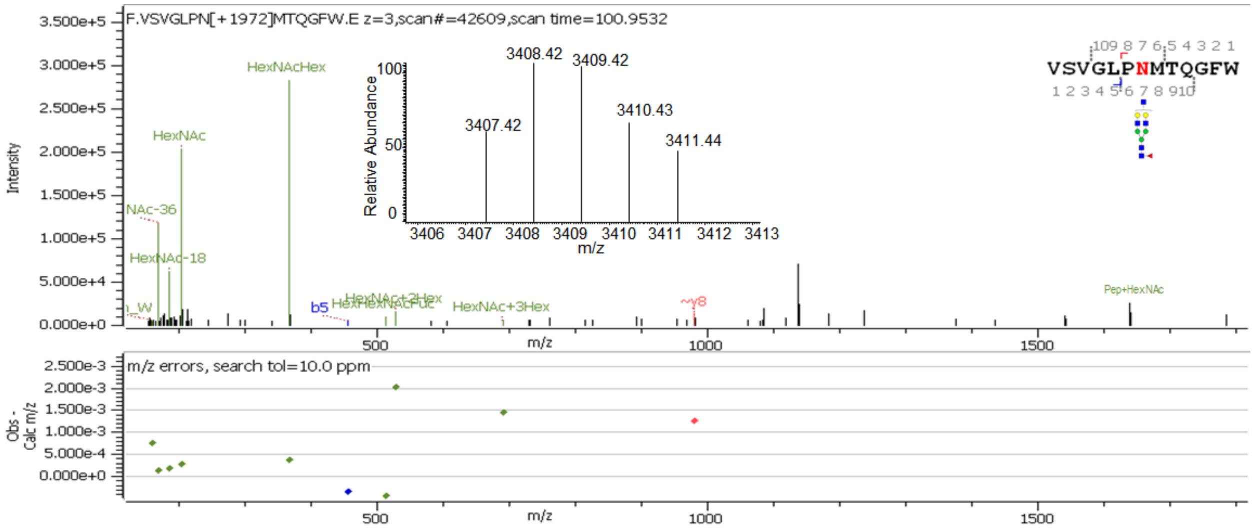
F.VSVGLPN[+1972]MTQGFW.E z=3,scan#=42609,scan time=100.9532
VSVGLPNMTQGFW
Intensity
Relative Abundance
3407.42
3408.42
3409.42
3410.43
3411.44
m/z
HexNAcHex
HexNAc
HexNAc-18
b5
HexNAc+3Hex
~y8
Pep+HexNAc
m/z errors, search tol=10.0 ppm
Obs - Calc m/z

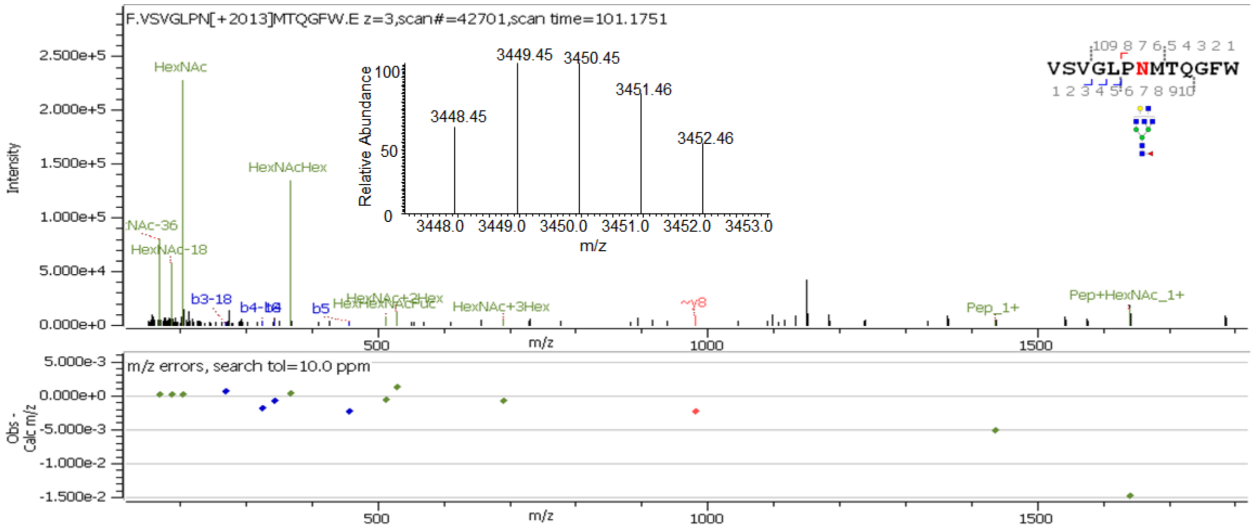
F.VSVGLPN[+2013]MTQGFW.E z=3,scan#=42701,scan time=101.1751
VSVGLPNMTQGFW
HexNAc
HexNAcHex
HexNAc-18
b3-18
b5
HexNAc+3Hex
Pep_1+
Pep+HexNAc_1+
3448.45
3449.45
3450.45
3451.46
3452.46
Relative Abundance
m/z
Intensity
m/z errors, search tol=10.0 ppm
Obs - Calc m/z

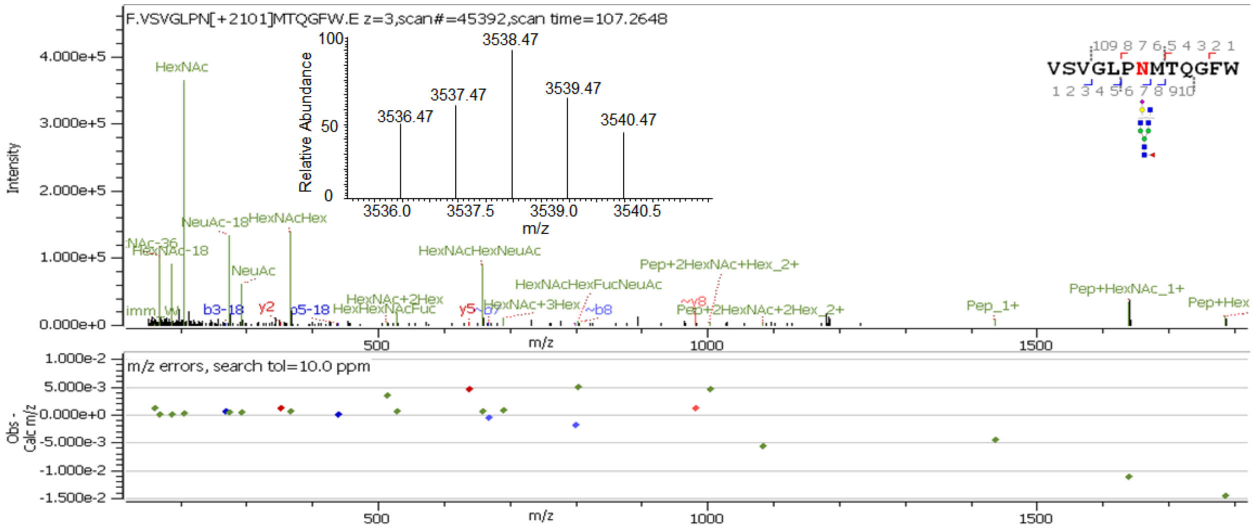
F.VSVGLPN[+2101]MTQGFW.E z=3,scan#=45392,scan time=107.2648
VSVGLPNMTQGFW
HexNAc
NeuAc-18
HexNAcHex
HexNAc-18
NeuAc
b3-18
y2
b5-18
HexNAcHexNeuAc
HexNAcHexFucNeuAc
Pep+2HexNAc+Hex_2+
Pep+2HexNAc+2Hex_2+
Pep_1+
Pep+HexNAc_1+
3536.47
3537.47
3538.47
3539.47
3540.47
Relative Abundance
m/z
Intensity
m/z errors, search tol=10.0 ppm
Obs - Calc m/z

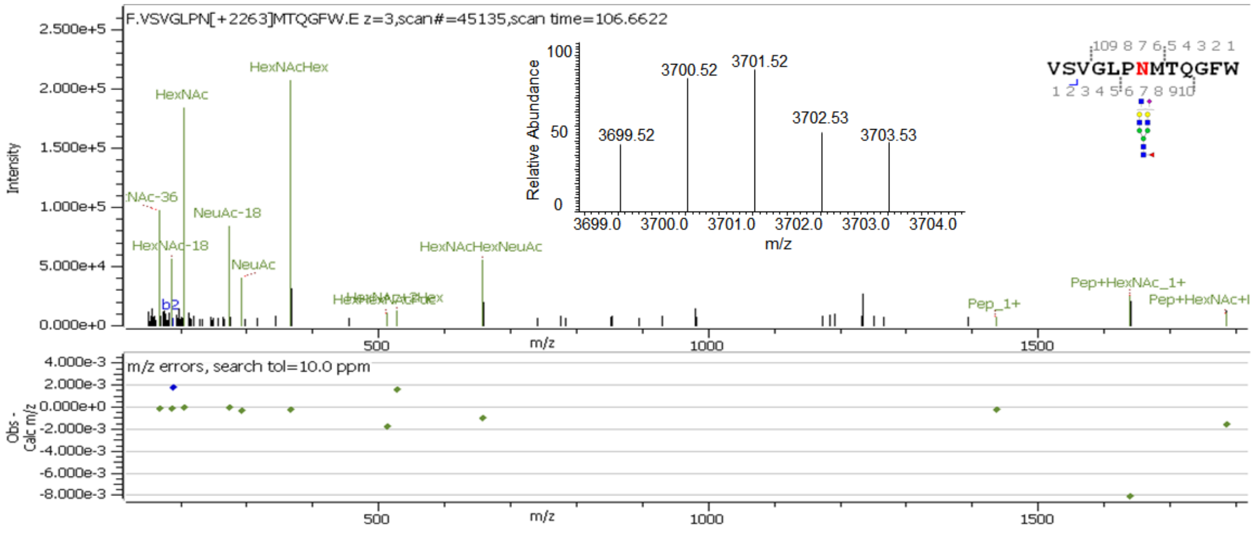
F.VSVGLPN[+2263]MTQGFW.E z=3,scan#=45135,scan time=106.6622
VSVGLPNMTQGFW
HexNAcHex
HexNAc
NeuAc-18
HexNAc-18
NeuAc
b2
HexNAcHexNeuAc
Pep_1+
Pep+HexNAc_1+
3699.52
3700.52
3701.52
3702.53
3703.53
Relative Abundance
m/z
Intensity
m/z errors, search tol=10.0 ppm
Obs - Calc m/z

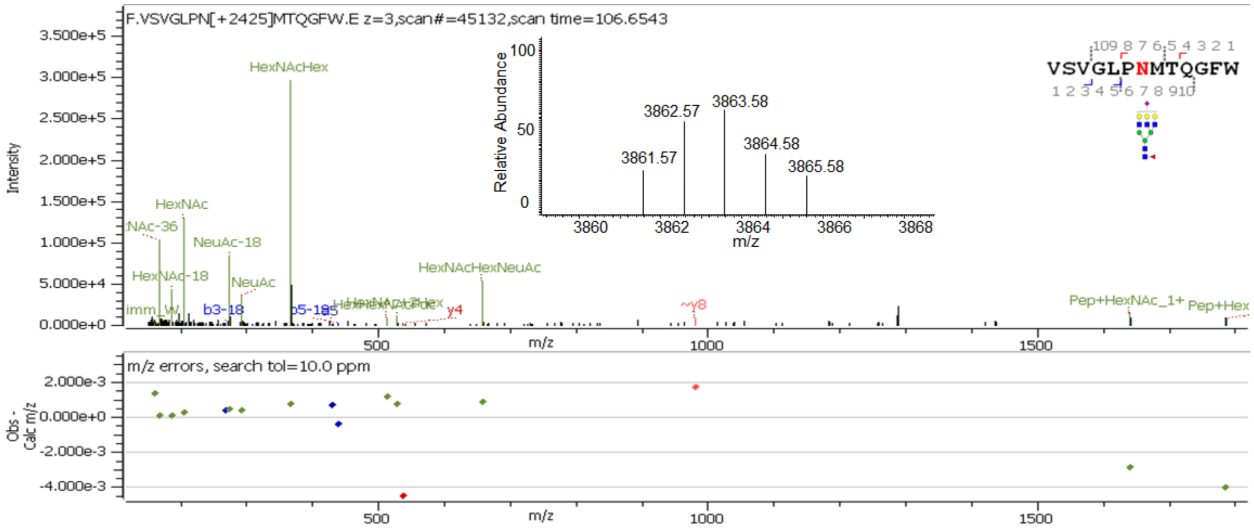
F.VSVGLPN[+2425]MTQGFW.E z=3,scan#=45132,scan time=106.6543
Intensity
Relative Abundance
3861.57
3862.57
3863.58
3864.58
3865.58
m/z
VSVGLPNMTQGFW
HexNAcHex
HexNAc
NeuAc-18
HexNAc-18
NeuAc
HexNAcHexNeuAc
b3-18
y4
~y8
Pep+HexNAc_1+
m/z errors, search tol=10.0 ppm
Obs - Calc m/z

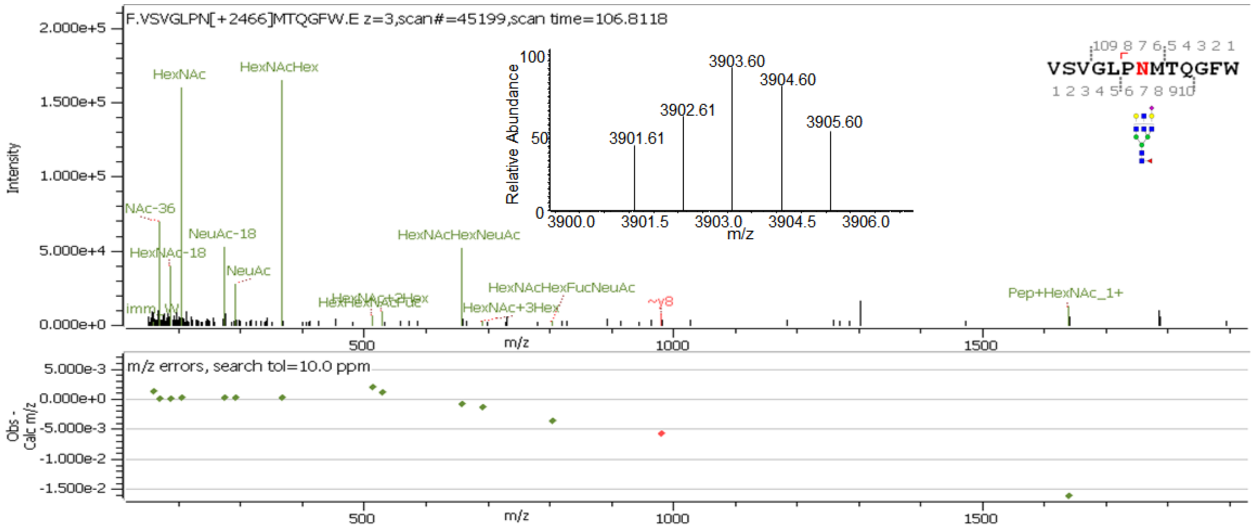
F.VSVGLPN[+2466]MTQGFW.E z=3,scan#=45199,scan time=106.8118
Intensity
Relative Abundance
3901.61
3902.61
3903.60
3904.60
3905.60
m/z
VSVGLPNMTQGFW
HexNAc
HexNAcHex
NeuAc-18
HexNAc-18
NeuAc
HexNAcHexNeuAc
HexNAcHexFucNeuAc
HexNAc+3Hex
~y8
Pep+HexNAc_1+
m/z errors, search tol=10.0 ppm
Obs - Calc m/z

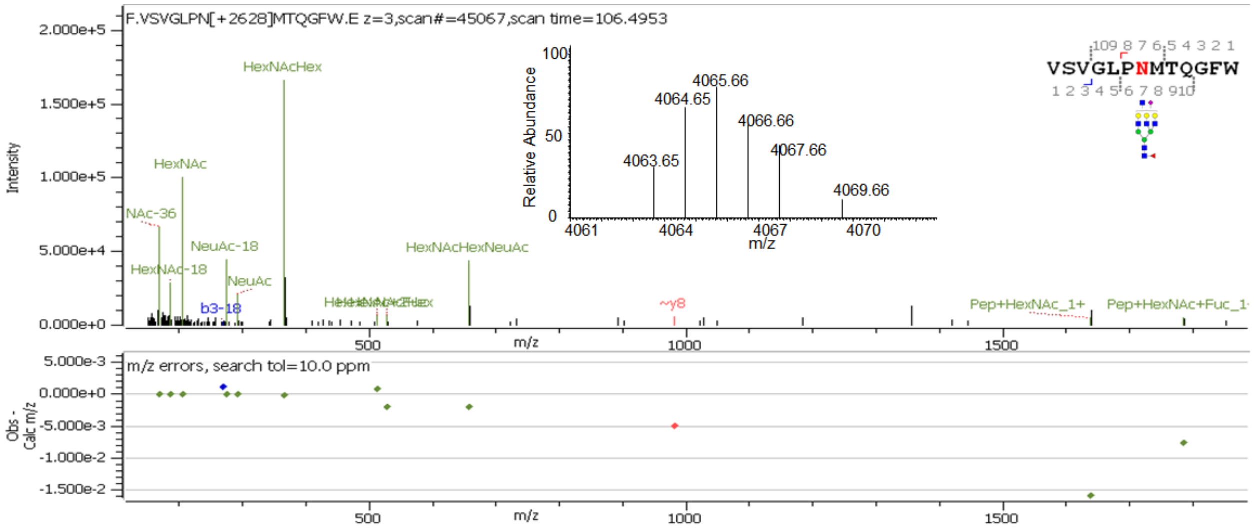
F.VSVGLPN[+2628]MTQGFW.E z=3,scan#=45067,scan time=106.4953
Intensity
Relative Abundance
4063.65
4064.65
4065.66
4066.66
4067.66
4069.66
m/z
VSVGLPNMTQGFW
HexNAcHex
HexNAc
NeuAc-18
HexNAc-18
NeuAc
b3-18
HexNAcHexNeuAc
~y8
Pep+HexNAc_1+
m/z errors, search tol=10.0 ppm
Obs - Calc m/z

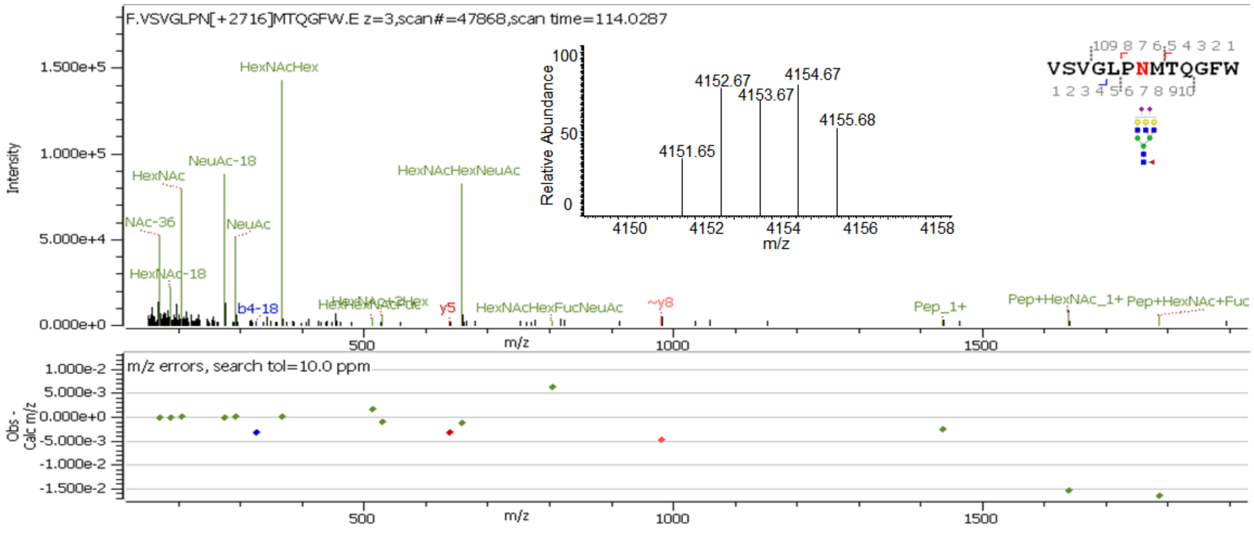
F.VSVGLPN[+2716]MTQGFW.E z=3,scan#=47868,scan time=114.0287
HexNAcHex
NeuAc-18
HexNAc
HexNAcHexNeuAc
NeuAc
HexNAc-18
b4-18
y5
HexNAcHexFucNeuAc
~y8
Pep_1+
Pep+HexNAc_1+
Pep+HexNAc+Fuc
4152.67
4153.67
4154.67
4155.68
4151.65
Relative Abundance
m/z
Intensity
m/z errors, search tol=10.0 ppm
Obs - Calc m/z
VSVGLPNMTQGFW

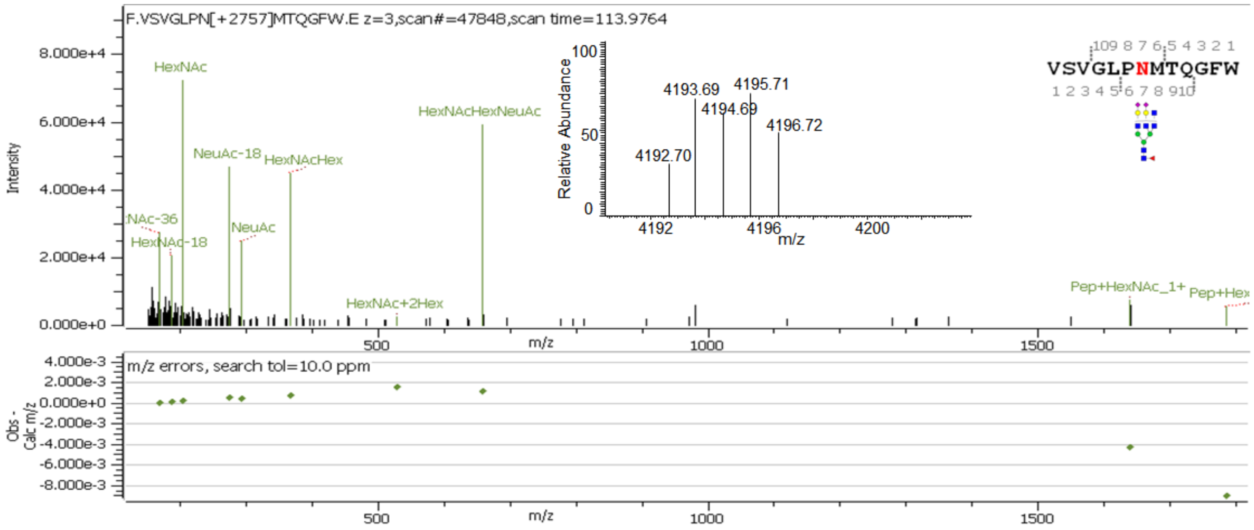
F.VSVGLPN[+2757]MTQGFW.E z=3,scan#=47848,scan time=113.9764
HexNAc
HexNAcHexNeuAc
NeuAc-18
HexNAcHex
NeuAc
HexNAc-18
HexNAc+2Hex
Pep+HexNAc_1+
4193.69
4195.71
4194.69
4196.72
4192.70
Relative Abundance
m/z
Intensity
m/z errors, search tol=10.0 ppm
Obs - Calc m/z
VSVGLPNMTQGFW

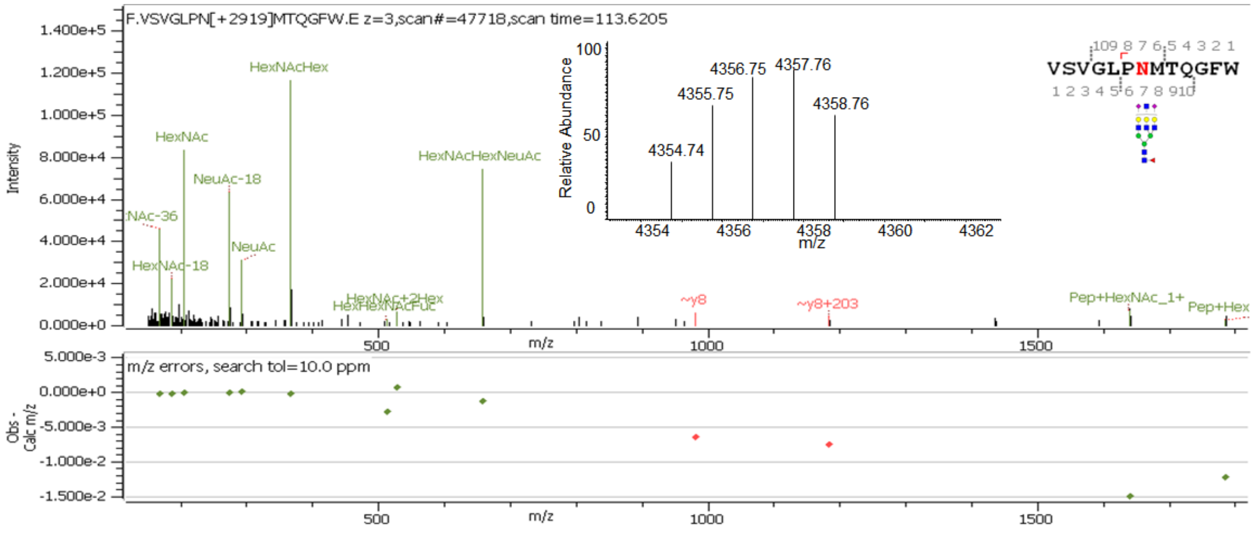
F.VSVGLPN[+2919]MTQGFW.E z=3,scan#=47718,scan time=113.6205
HexNAcHex
HexNAc
HexNAcHexNeuAc
NeuAc-18
NeuAc
HexNAc-18
HexNAc+2Hex
HexHexNAcFuc
~y8
~y8+203
Pep+HexNAc_1+
4356.75
4357.76
4355.75
4358.76
4354.74
Relative Abundance
m/z
Intensity
m/z errors, search tol=10.0 ppm
Obs - Calc m/z
VSVGLPNMTQGFW

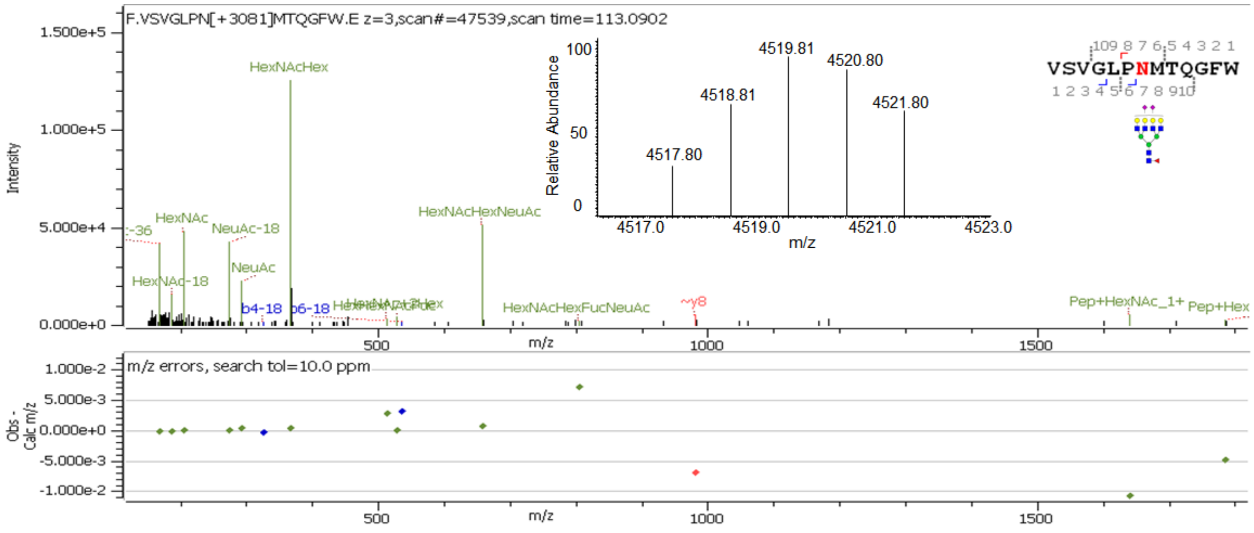


**Supp. Figure S4:** Deconvoluted precursor ion isotopic patterns (insets) and annotated HCD MS/MS spectra of intact N-glycopeptides with assigned N-glycans at N322.

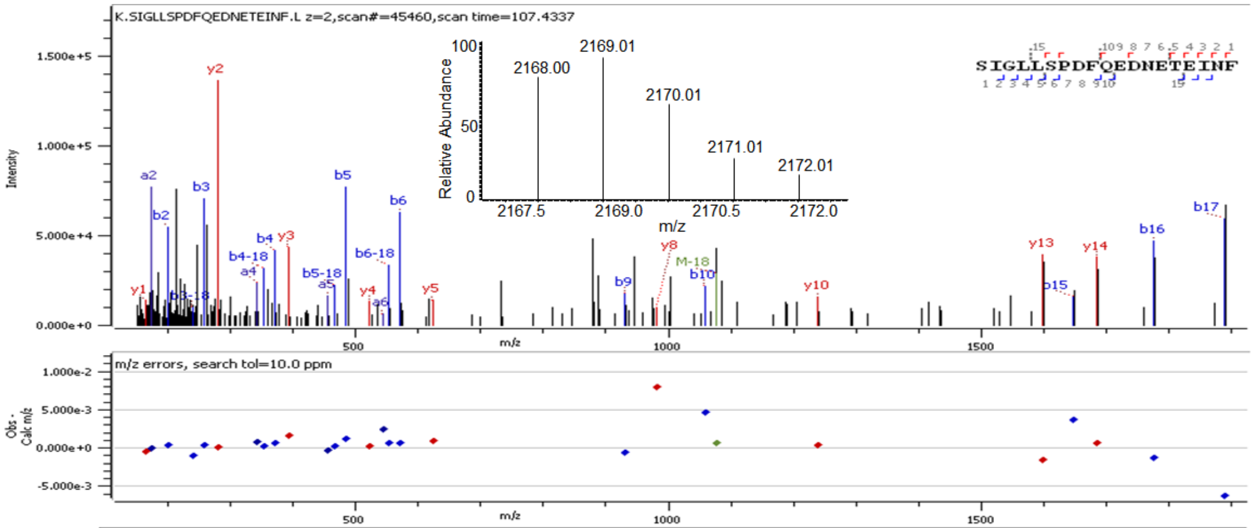


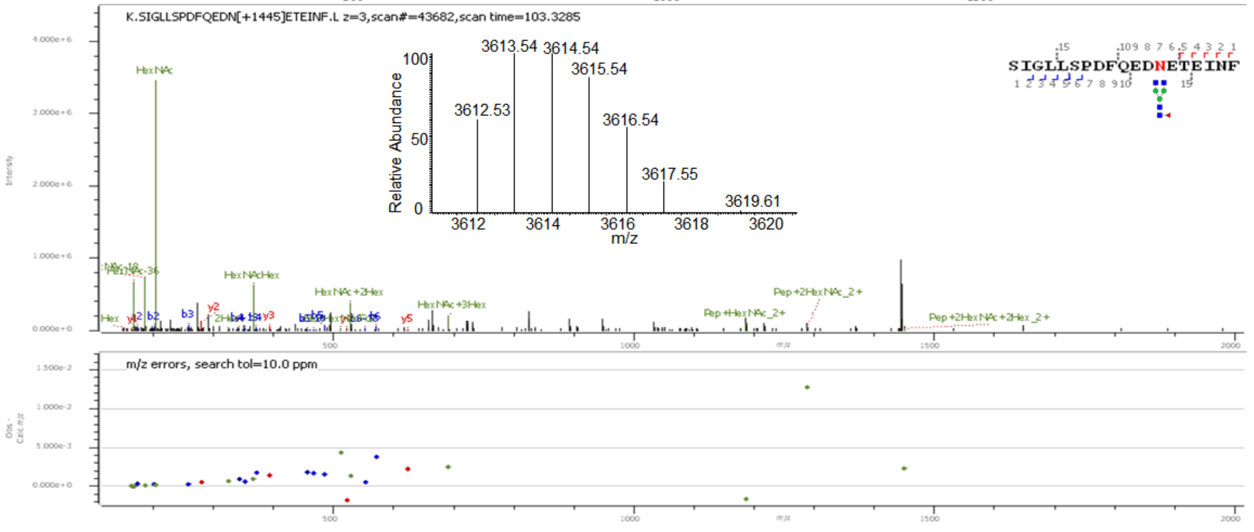

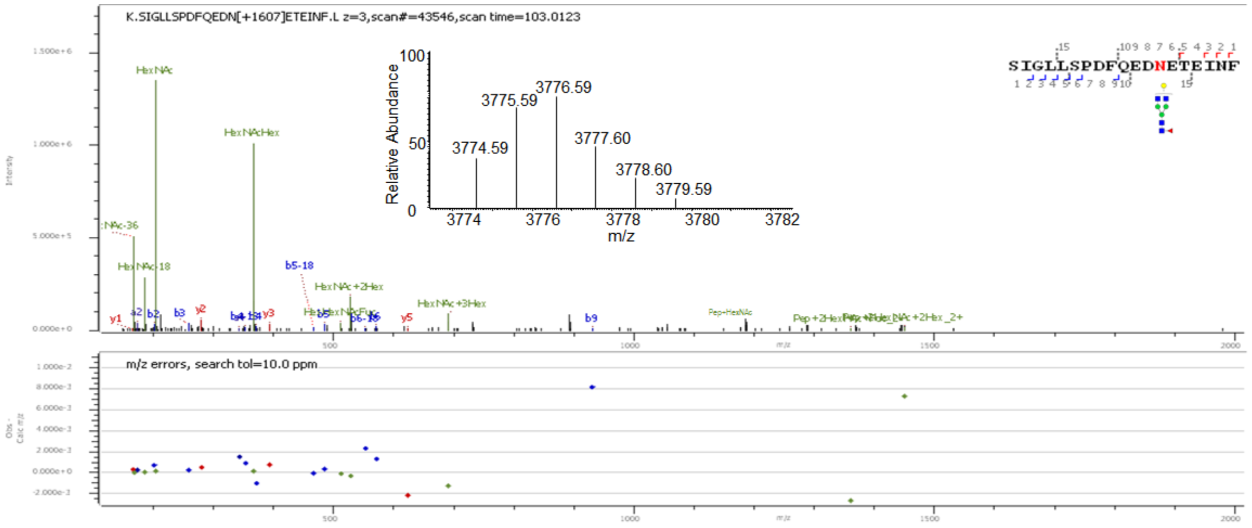
K.SIGLLSPDFQEDN[+1607]ETEINF.L z=3,scan#=43546,scan time=103.0123
SIGLLSPDFQEDNETEINF
Relative Abundance
3774.59
3775.59
3776.59
3777.60
3778.60
3779.59
m/z
HexNAc
HexNAcHex
HexNAc-18
b5-18
HexNAc+2Hex
HexNAc+3Hex
y1
b2
b3
y2
y3
y5
b9
m/z errors, search tol=10.0 ppm

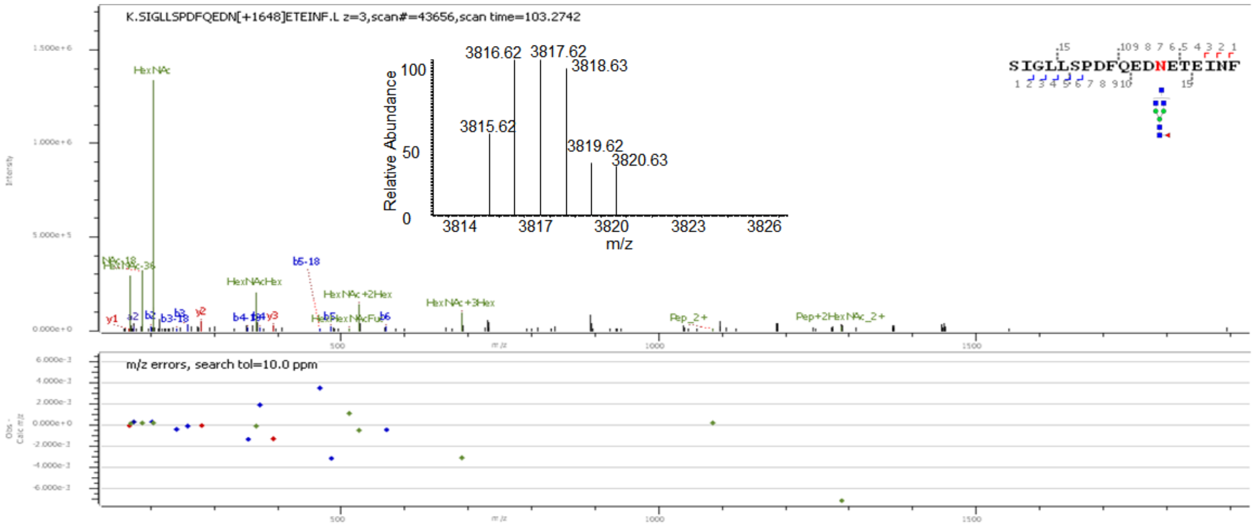
K.SIGLLSPDFQEDN[+1648]ETEINF.L z=3,scan#=43656,scan time=103.2742
SIGLLSPDFQEDNETEINF
Relative Abundance
3815.62
3816.62
3817.62
3818.63
3819.62
3820.63
m/z
HexNAc
HexNAcHex
b5-18
HexNAc+2Hex
HexNAc+3Hex
Pep_2+
Pep+2HexNAc_2+
m/z errors, search tol=10.0 ppm

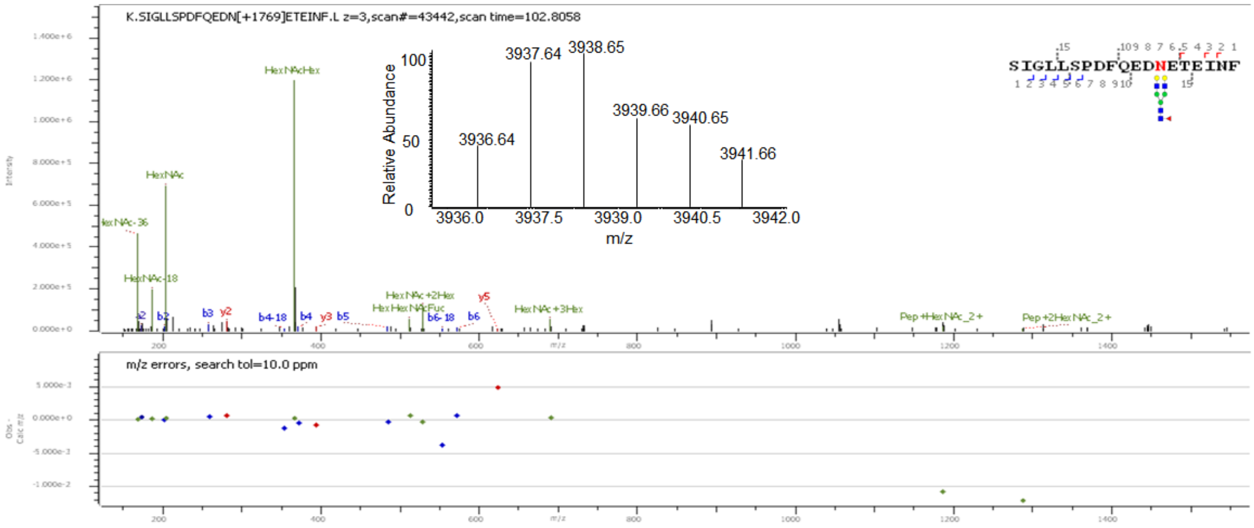
K.SIGLLSPDFQEDN[+1769]ETEINF.L z=3,scan#=43442,scan time=102.8058
SIGLLSPDFQEDNETEINF
Relative Abundance
3936.64
3937.64
3938.65
3939.66
3940.65
3941.66
m/z
HexNAcHex
HexNAc
HexNAc-36
HexNAc-18
HexNAc+2Hex
HexNAc+3Hex
Pep+HexNAc_2+
Pep+2HexNAc_2+
m/z errors, search tol=10.0 ppm

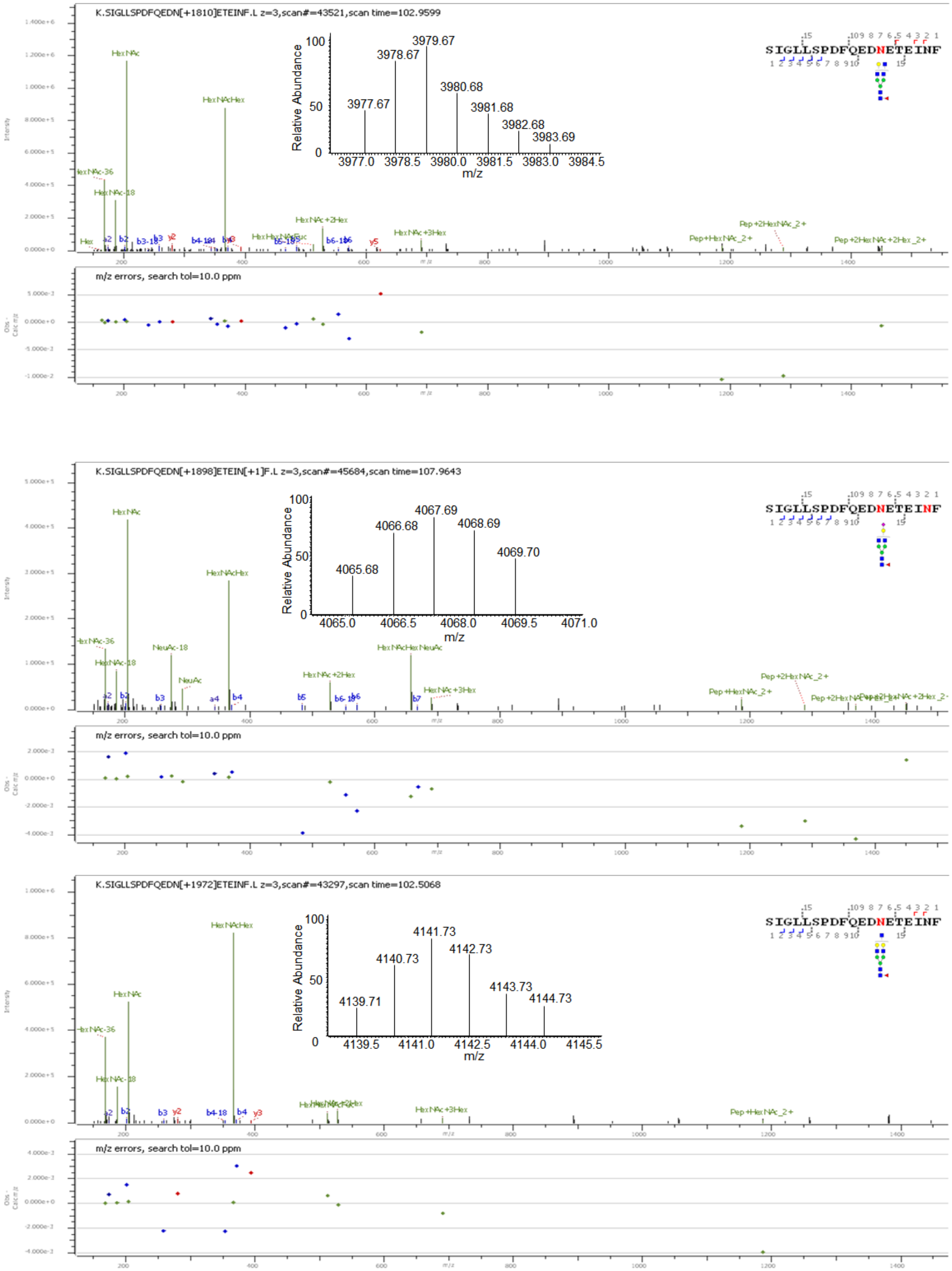
K.SIGLLSPDFQEDN[+1810]ETEINF.L z=3,scan#=43521,scan time=102.9599
m/z errors, search tol=10.0 ppm
K.SIGLLSPDFQEDN[+1898]ETEIN[+1]F.L z=3,scan#=45684,scan time=107.9643
m/z errors, search tol=10.0 ppm
K.SIGLLSPDFQEDN[+1972]ETEINF.L z=3,scan#=43297,scan time=102.5068
m/z errors, search tol=10.0 ppm
Relative Abundance
m/z
SIGLLSPDFQEDNETEINF

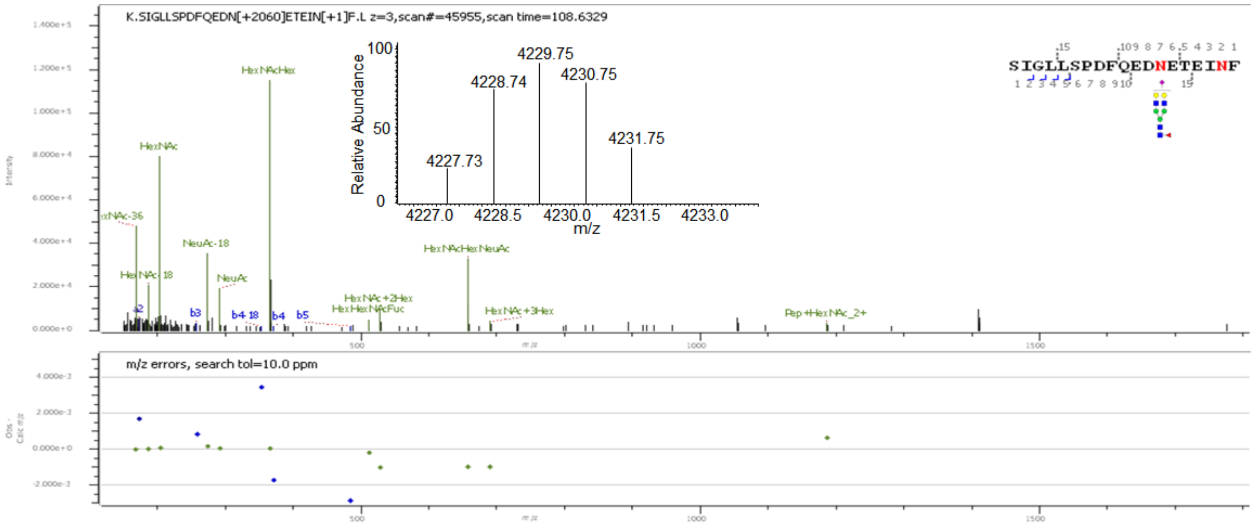
K.SIGLLSPDFQEDN[+2060]ETEIN[+1]F.L z=3,scan#=45955,scan time=108.6329
Relative Abundance
m/z
4227.73
4228.74
4229.75
4230.75
4231.75
SIGLLSPDFQEDNETEINF
HexNAcHex
HexNAc
NeuAc-18
NeuAc
HexNAcHexNeuAc
HexNAc+2Hex
HexHexNAcFuc
HexNAc+3Hex
Pep+HexNAc_2+
m/z errors, search tol=10.0 ppm

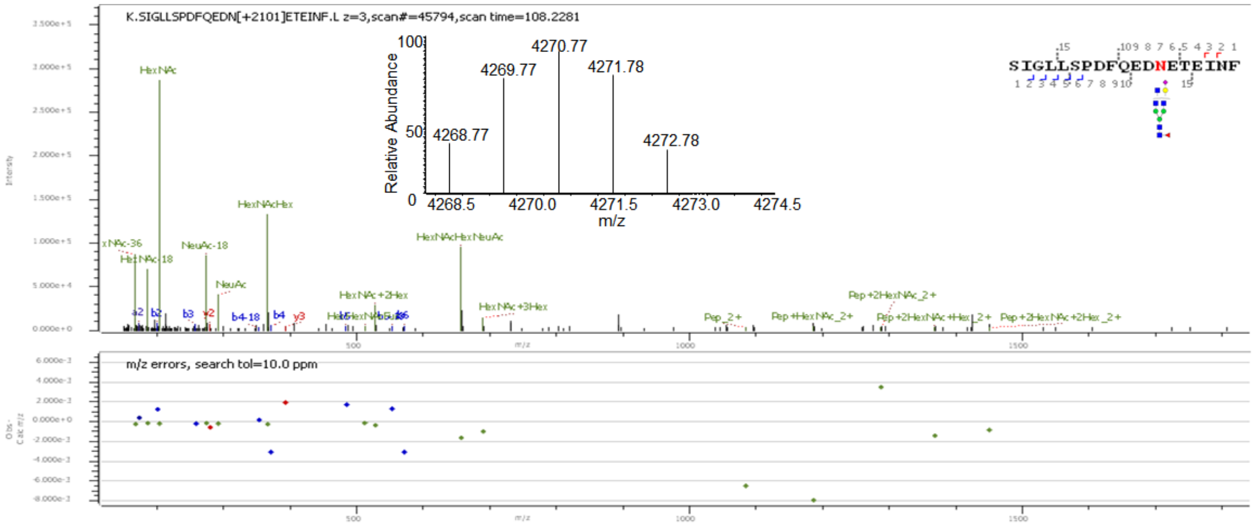
K.SIGLLSPDFQEDN[+2101]ETEINF.L z=3,scan#=45794,scan time=108.2281
Relative Abundance
m/z
4268.77
4269.77
4270.77
4271.78
4272.78
SIGLLSPDFQEDNETEINF
HexNAc
HexNAcHex
NeuAc-18
NeuAc
HexNAcHexNeuAc
HexNAc+2Hex
HexNAc+3Hex
Pep_2+
Pep+HexNAc_2+
Pep+2HexNAc_2+
m/z errors, search tol=10.0 ppm

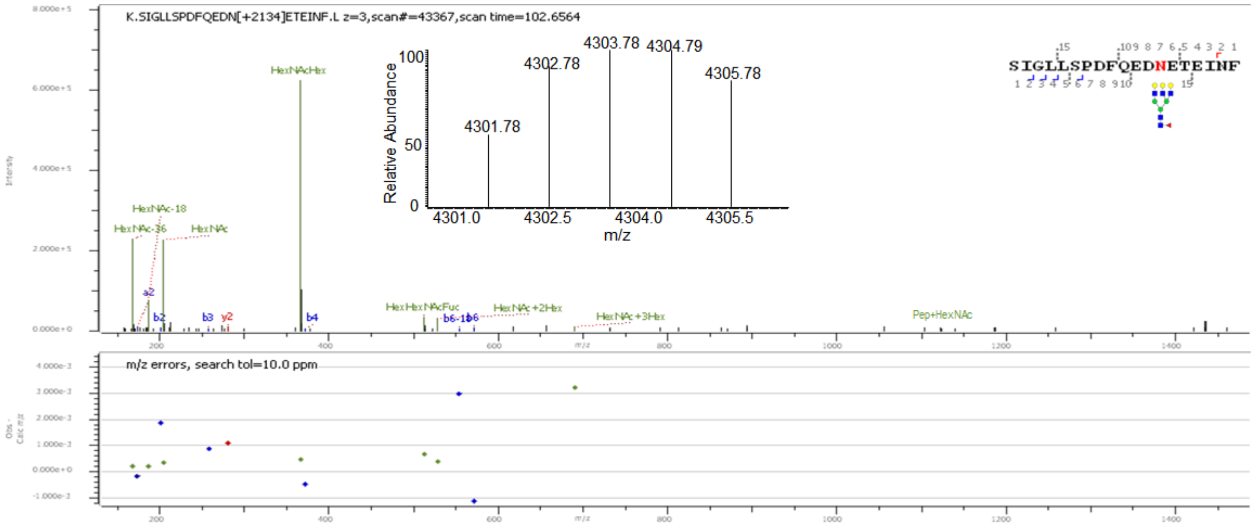
K.SIGLLSPDFQEDN[+2134]ETEINF.L z=3,scan#=43367,scan time=102.6564
Relative Abundance
m/z
4301.78
4302.78
4303.78
4304.79
4305.78
SIGLLSPDFQEDNETEINF
HexNAcHex
HexNAc-18
HexNAc-36
HexNAc
HexHexNAcFuc
HexNAc+2Hex
HexNAc+3Hex
Pep+HexNAc
m/z errors, search tol=10.0 ppm

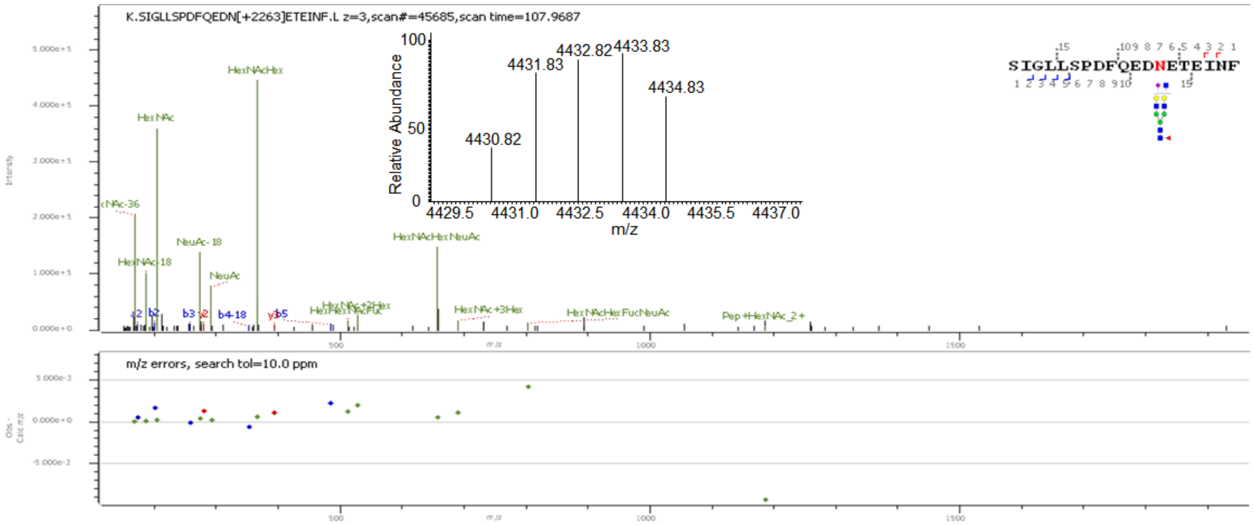
K.SIGLLSPDFQEDN[+2263]ETEINF.L z=3,scan#=45685,scan time=107.9687
Relative Abundance
4430.82
4431.83
4432.82
4433.83
4434.83
m/z
HexNAcHex
HexNAc
HexNAc-18
NeuAc-18
NeuAc
HexNAcHexNeuAc
HexNAc+3Hex
HexNAcHexFucNeuAc
Pep+HexNAc_2+
m/z errors, search tol=10.0 ppm
SIGLLSPDFQEDNETEINF

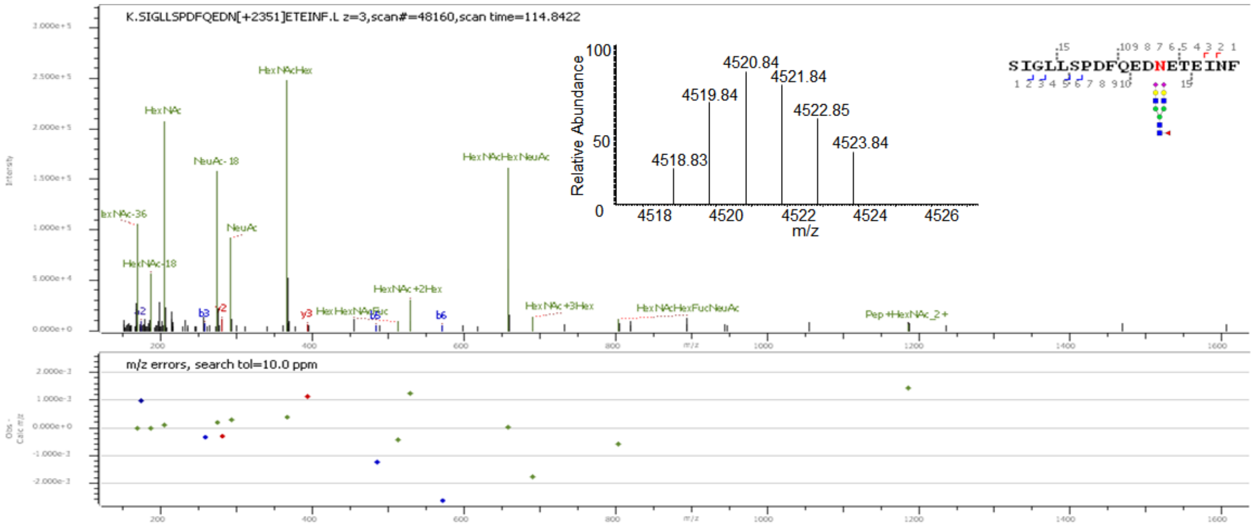
K.SIGLLSPDFQEDN[+2351]ETEINF.L z=3,scan#=48160,scan time=114.8422
Relative Abundance
4518.83
4519.84
4520.84
4521.84
4522.85
4523.84
m/z
HexNAcHex
HexNAc
NeuAc-18
NeuAc
HexNAc-18
HexNAcHexNeuAc
HexNAc+2Hex
HexNAc+3Hex
HexNAcHexFucNeuAc
Pep+HexNAc_2+
m/z errors, search tol=10.0 ppm
SIGLLSPDFQEDNETEINF

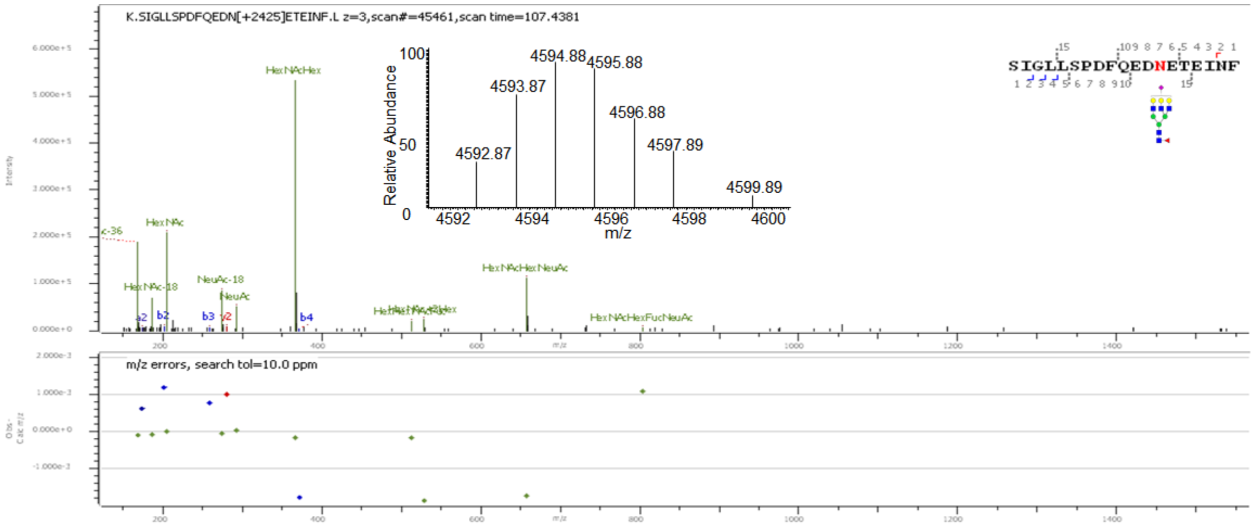
K.SIGLLSPDFQEDN[+2425]ETEINF.L z=3,scan#=45461,scan time=107.4381
Relative Abundance
4592.87
4593.87
4594.88
4595.88
4596.88
4597.89
4599.89
m/z
HexNAcHex
HexNAc
HexNAc-18
NeuAc-18
NeuAc
HexNAcHexNeuAc
HexNAcHexFucNeuAc
m/z errors, search tol=10.0 ppm
SIGLLSPDFQEDNETEINF

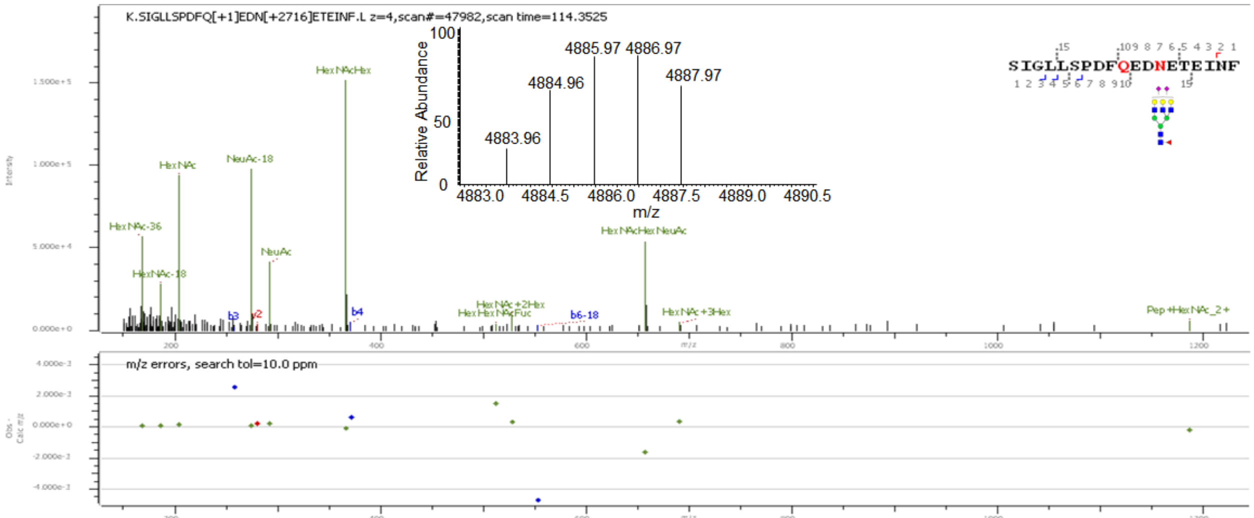


**Supp. Figure S5:** Deconvoluted precursor ion isotopic patterns (insets) and annotated HCD MS/MS spectra of intact N-glycopeptides with assigned N-glycans at N432.

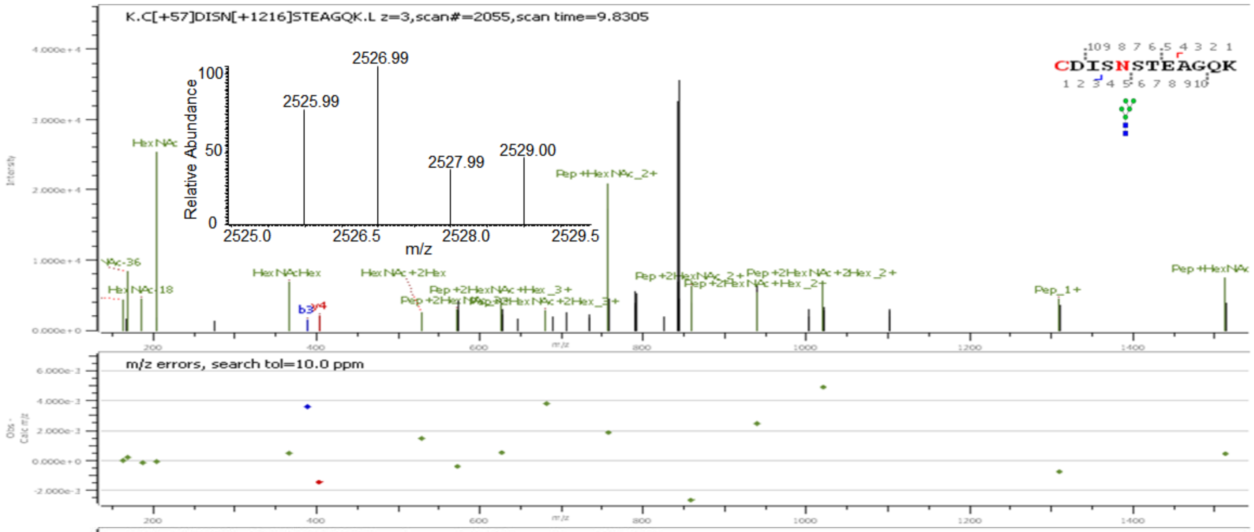


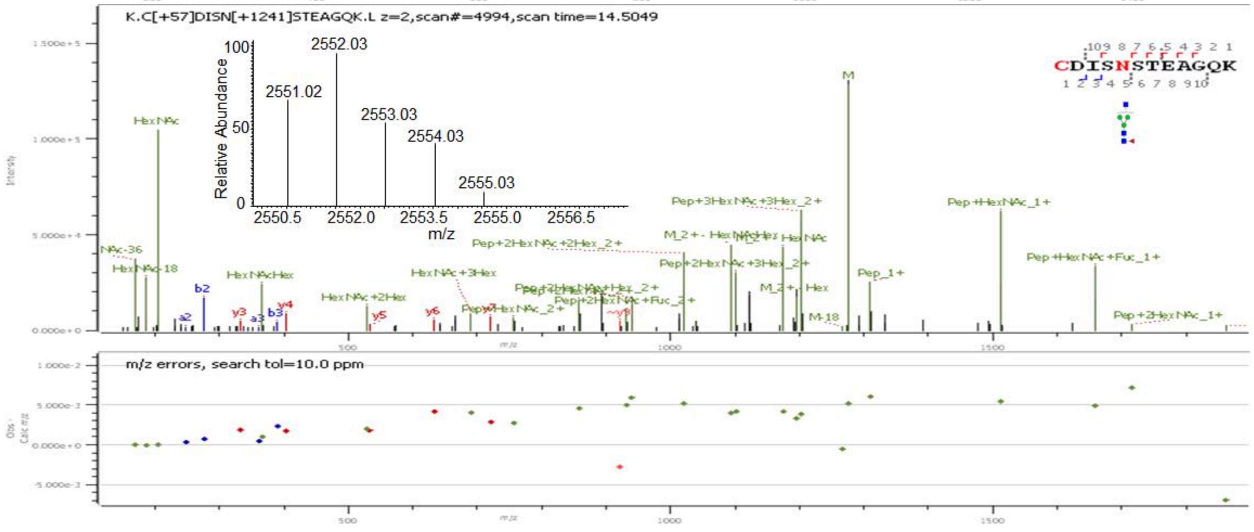

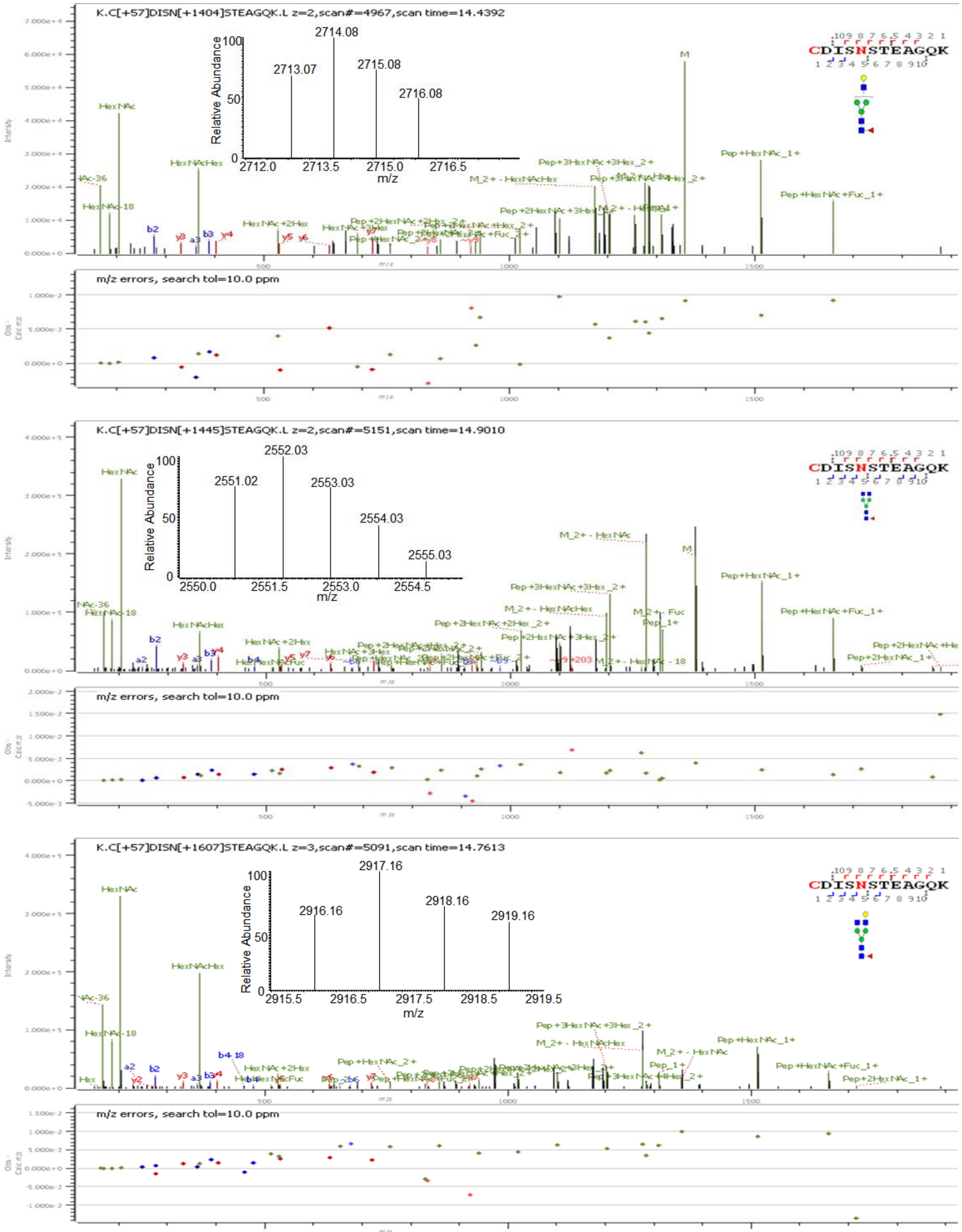
K.C[+57]DISN[+1404]STEAGQK.L z=2,scan#=4967,scan time=14.4392
Relative Abundance
m/z
2713.07
2714.08
2715.08
2716.08
CDISNSTEAGQK
m/z errors, search tol=10.0 ppm
K.C[+57]DISN[+1445]STEAGQK.L z=2,scan#=5151,scan time=14.9010
2551.02
2552.03
2553.03
2554.03
2555.03
m/z errors, search tol=10.0 ppm
K.C[+57]DISN[+1607]STEAGQK.L z=3,scan#=5091,scan time=14.7613
2916.16
2917.16
2918.16
2919.16
m/z errors, search tol=10.0 ppm

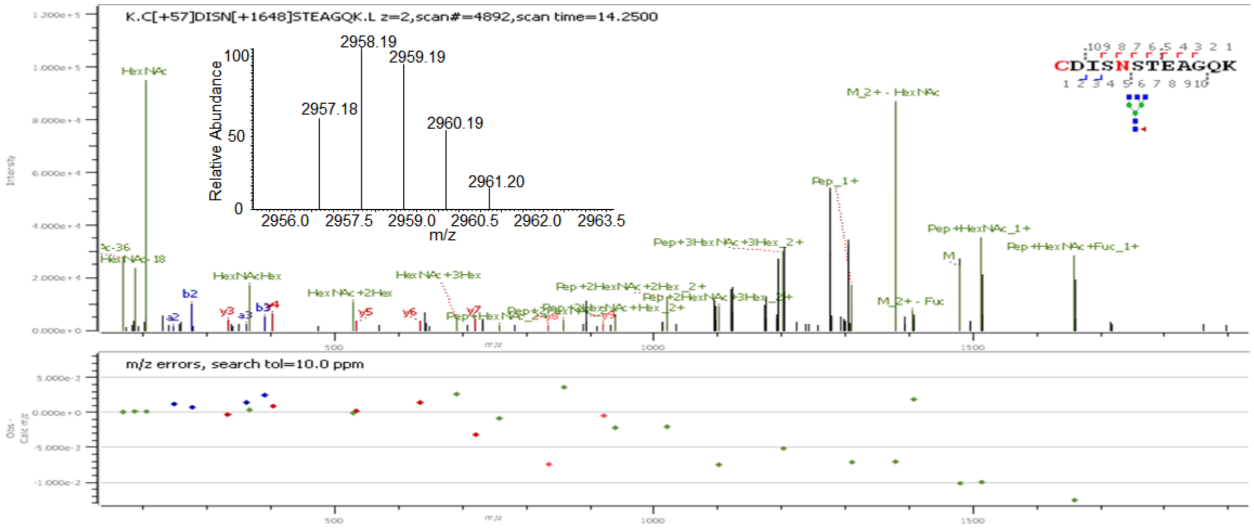
K.C[+57]DISN[+1648]STEAGQK.L z=2,scan#=4892,scan time=14.2500
Relative Abundance
m/z
2958.19
2959.19
2957.18
2960.19
2961.20
CDISNSTEAGQK
HexNAc
M_2+ - HexNAc
Pep_1+
Pep+HexNAc_1+
Pep+HexNAc+Fuc_1+
Pep+3HexNAc+3Hex_2+
HexNAcHex
HexNAc+3Hex
HexNAc+2Hex
m/z errors, search tol=10.0 ppm

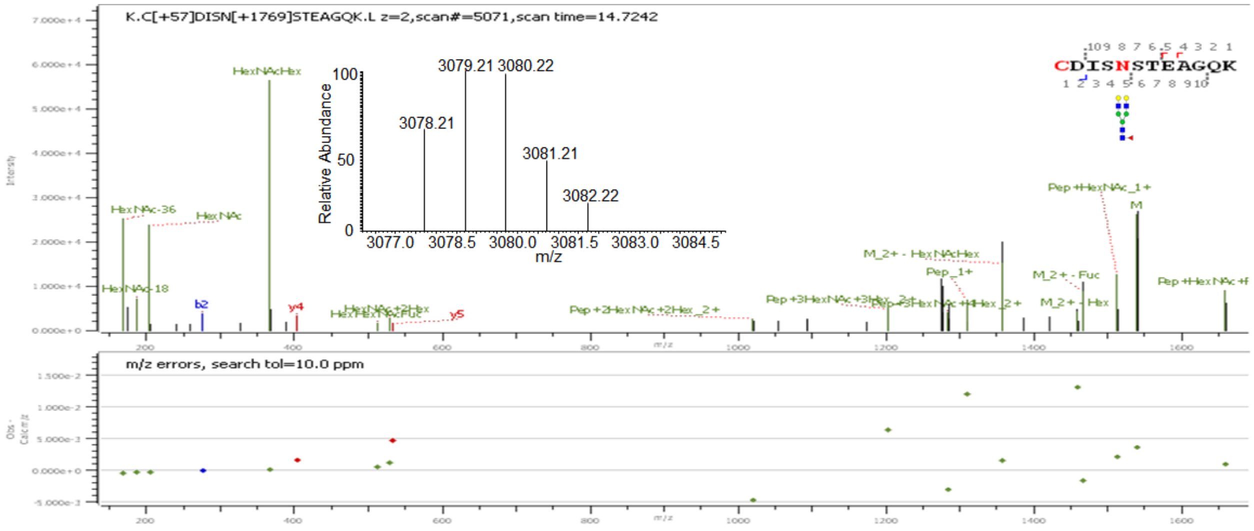
K.C[+57]DISN[+1769]STEAGQK.L z=2,scan#=5071,scan time=14.7242
Relative Abundance
m/z
3079.21
3080.22
3078.21
3081.21
3082.22
CDISNSTEAGQK
HexNAcHex
HexNAc-36
HexNAc
HexNAc-18
Pep+HexNAc_1+
M_2+ - HexNAcHex
Pep_1+
M_2+ - Fuc
M_2+ - Hex
Pep+2HexNAc+2Hex_2+
Pep+3HexNAc+3Hex_2+
m/z errors, search tol=10.0 ppm

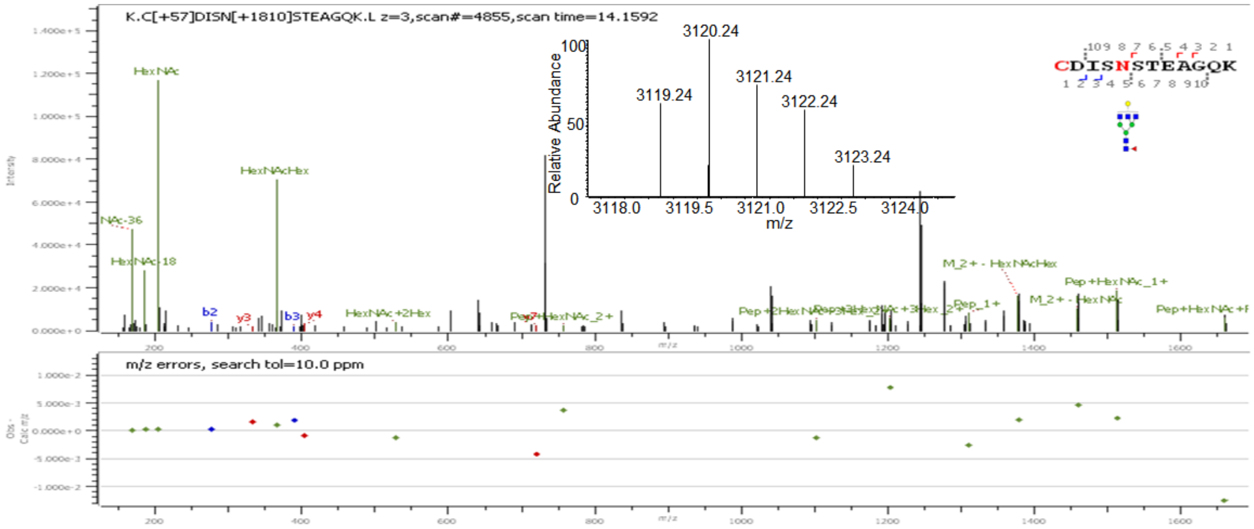
K.C[+57]DISN[+1810]STEAGQK.L z=3,scan#=4855,scan time=14.1592
Relative Abundance
m/z
3120.24
3121.24
3119.24
3122.24
3123.24
CDISNSTEAGQK
HexNAc
HexNAcHex
HexNAc-18
M_2+ - HexNAcHex
Pep+HexNAc_1+
M_2+ - HexNAc
Pep_1+
HexNAc+2Hex
m/z errors, search tol=10.0 ppm

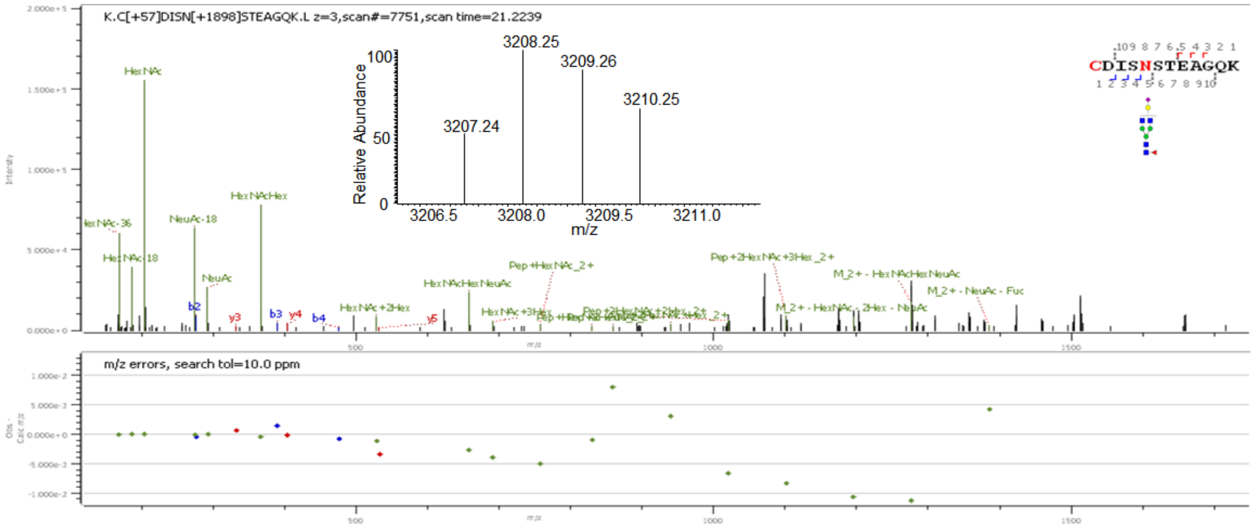


**Supp. Figure S6:** Deconvoluted precursor ion isotopic patterns (insets) and annotated HCD MS/MS spectra of intact N-glycopeptides with assigned N-glycans at N546.

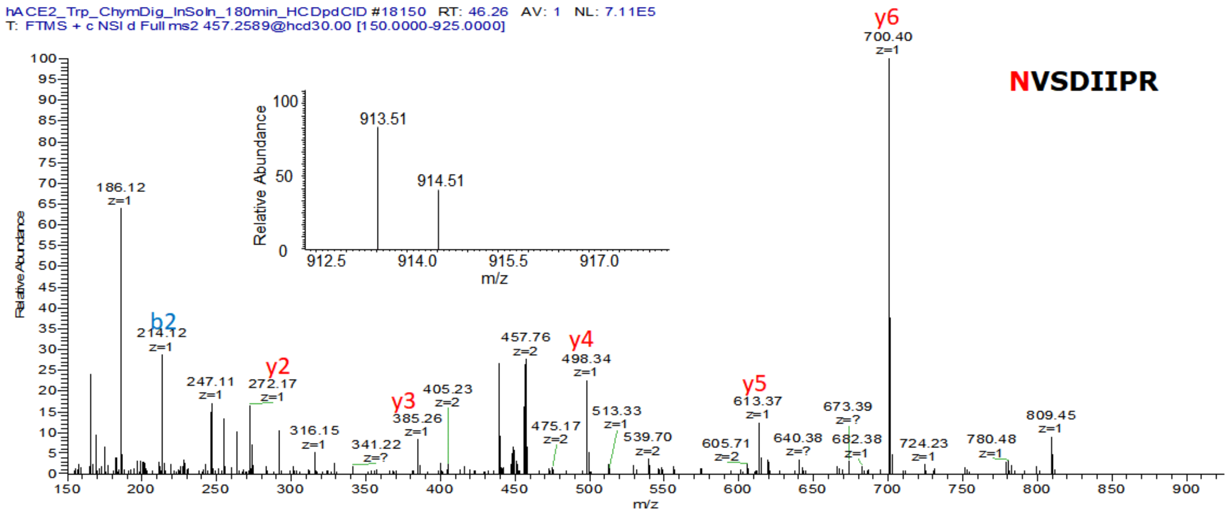


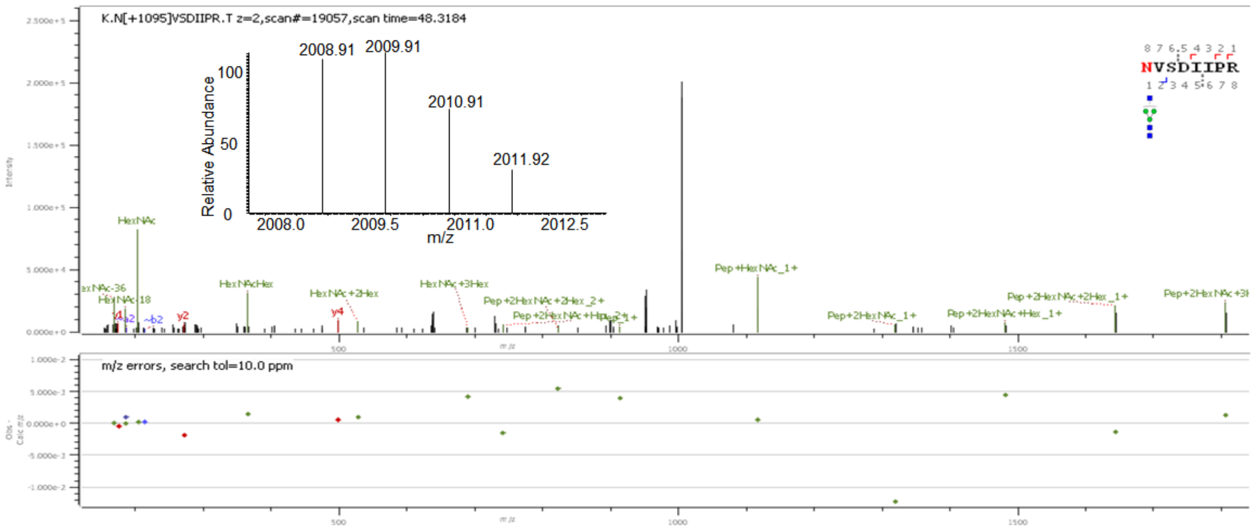

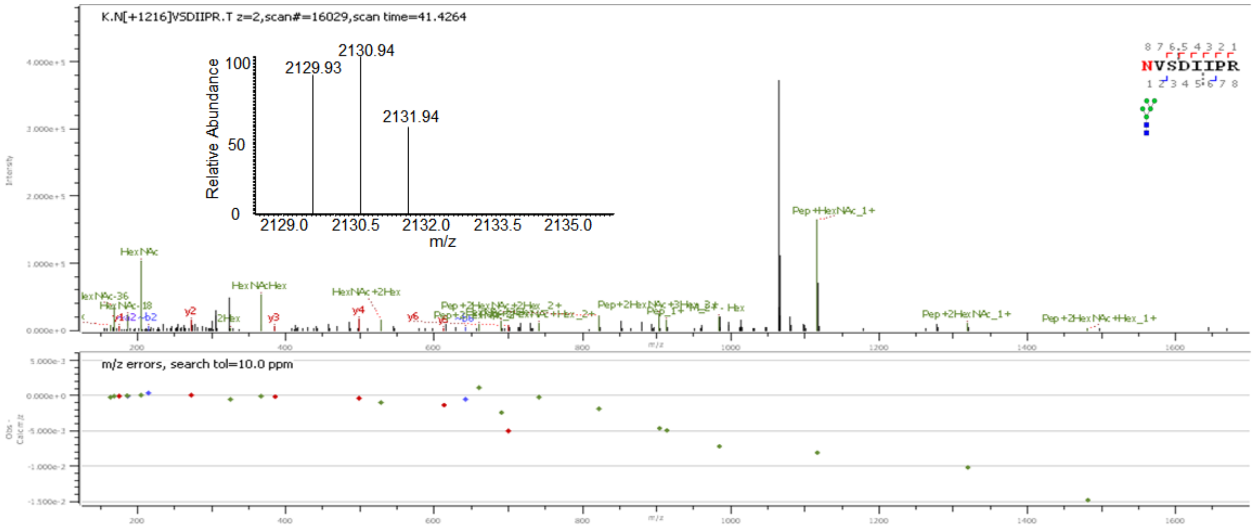
K.N[+1216]VSDIIPR.T z=2,scan#=16029,scan time=41.4264
NVSDIIPR
Relative Abundance
m/z
2129.93
2130.94
2131.94
HexNAc
HexNAcHex
HexNAc+2Hex
Pep+HexNAc_1+
Pep+2HexNAc_1+
Pep+2HexNAc+Hex_1+
m/z errors, search tol=10.0 ppm

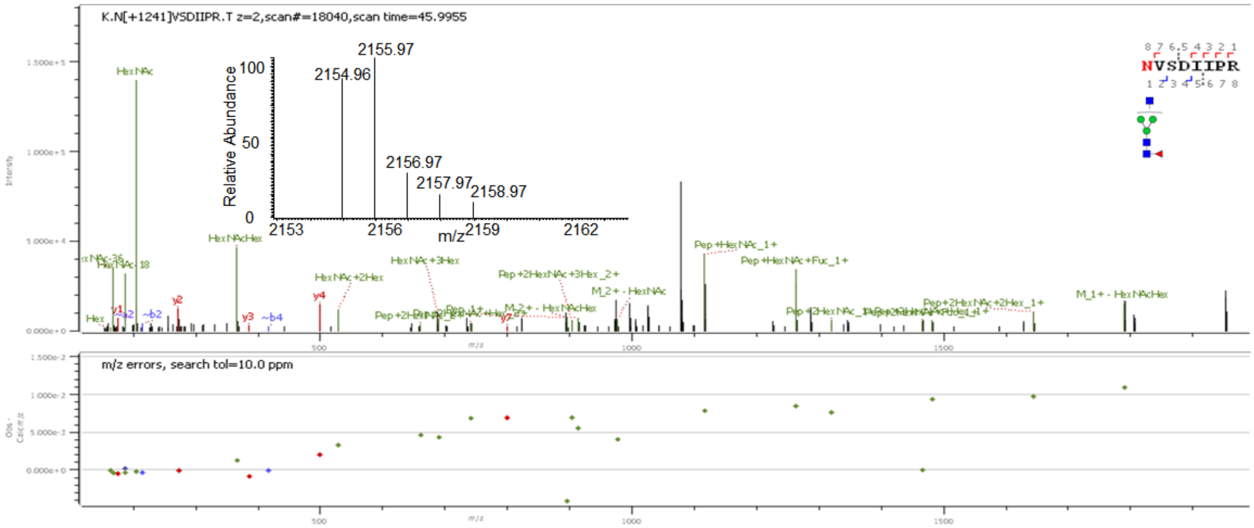
K.N[+1241]VSDIIPR.T z=2,scan#=18040,scan time=45.9955
NVSDIIPR
Relative Abundance
m/z
2154.96
2155.97
2156.97
2157.97
2158.97
HexNAc
HexNAcHex
HexNAc+2Hex
HexNAc+3Hex
Pep+2HexNAc+3Hex_2+
M_2+ - HexNAc
Pep+HexNAc_1+
Pep+HexNAc+Fuc_1+
Pep+2HexNAc+2Hex_1+
M_1+ - HexNAcHex
m/z errors, search tol=10.0 ppm

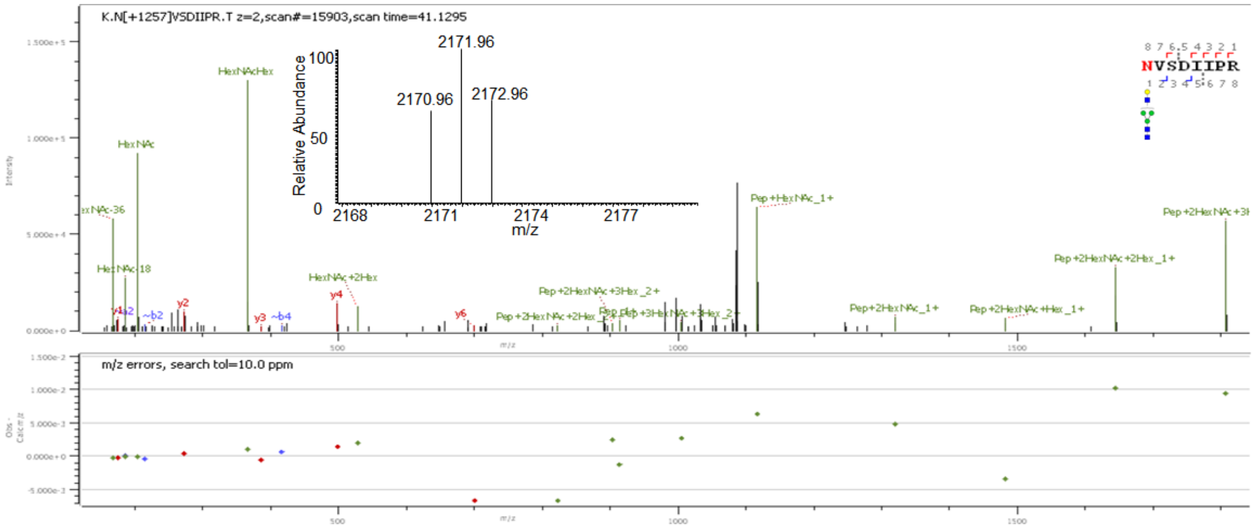
K.N[+1257]VSDIIPR.T z=2,scan#=15903,scan time=41.1295
NVSDIIPR
Relative Abundance
m/z
2170.96
2171.96
2172.96
HexNAcHex
HexNAc
HexNAc+2Hex
Pep+2HexNAc+3Hex_2+
Pep+HexNAc_1+
Pep+2HexNAc_1+
Pep+2HexNAc+Hex_1+
Pep+2HexNAc+2Hex_1+
m/z errors, search tol=10.0 ppm

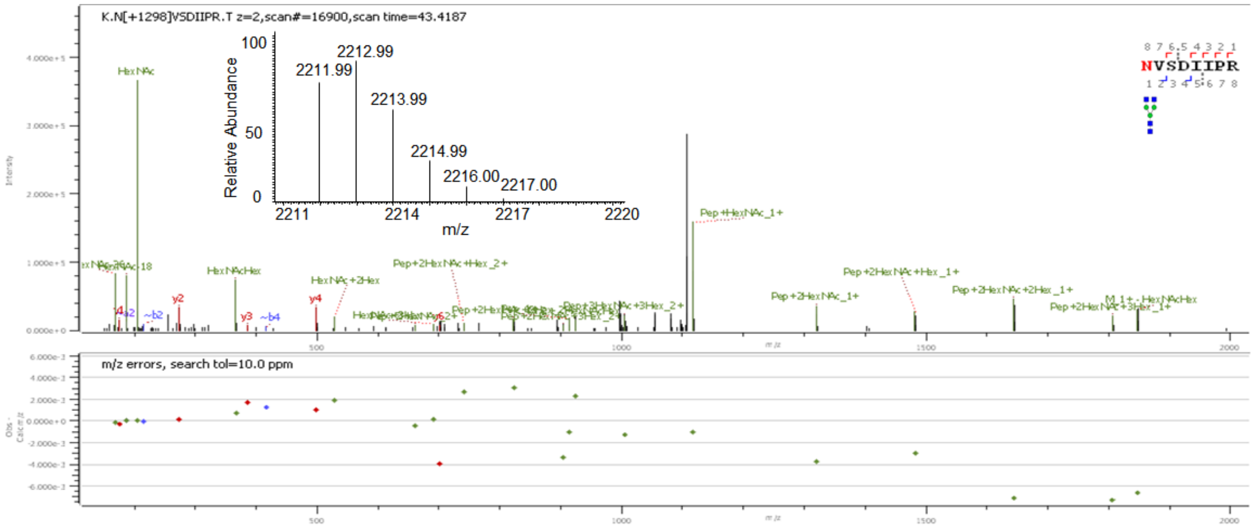
K.N[+1298]VSDIIPR.T z=2,scan#=16900,scan time=43.4187
Relative Abundance
2211.99
2212.99
2213.99
2214.99
2216.00
2217.00
m/z
Pep+HexNAc_1+
Pep+2HexNAc_1+
Pep+2HexNAc+Hex_1+
Pep+2HexNAc+2Hex_1+
m/z errors, search tol=10.0 ppm

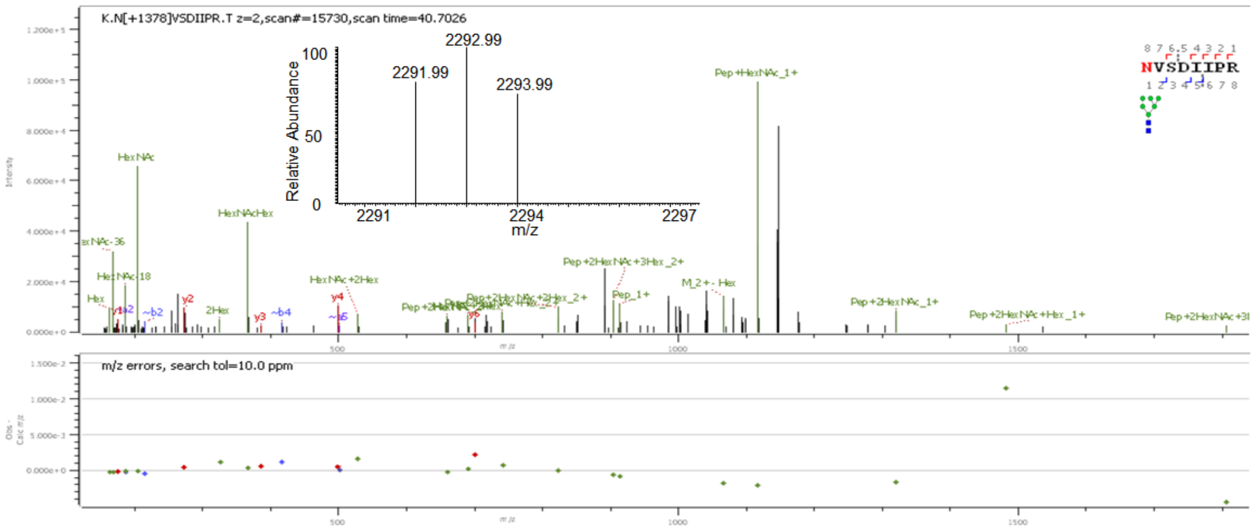
K.N[+1378]VSDIIPR.T z=2,scan#=15730,scan time=40.7026
Relative Abundance
2291.99
2292.99
2293.99
m/z
Pep+HexNAc_1+
Pep+2HexNAc_1+
Pep+2HexNAc+Hex_1+
m/z errors, search tol=10.0 ppm

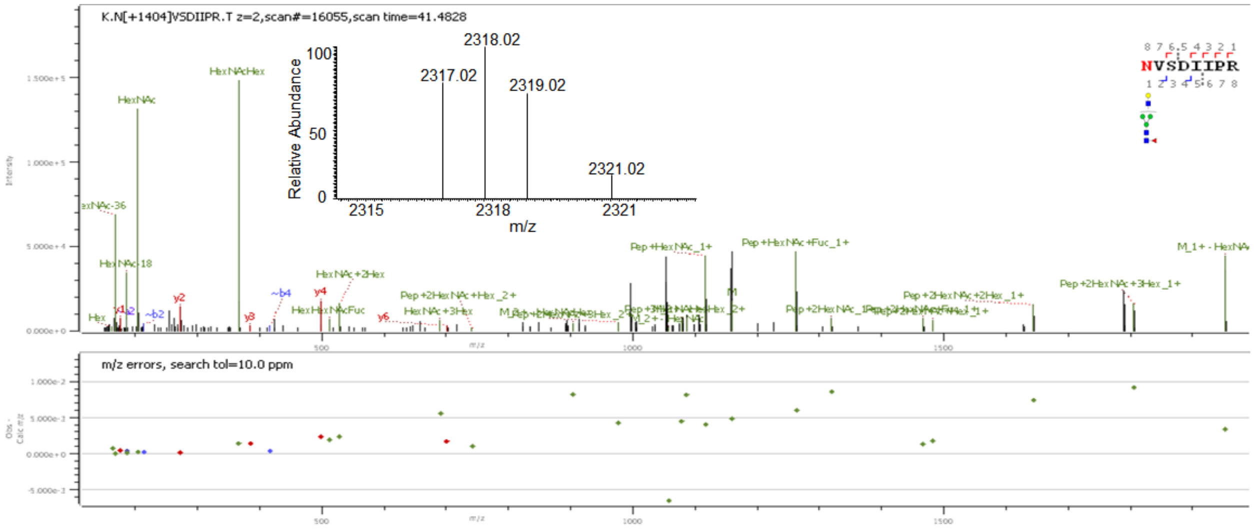
K.N[+1404]VSDIIPR.T z=2,scan#=16055,scan time=41.4828
Relative Abundance
2317.02
2318.02
2319.02
2321.02
m/z
Pep+HexNAc_1+
Pep+HexNAc+Fuc_1+
Pep+2HexNAc+3Hex_1+
m/z errors, search tol=10.0 ppm

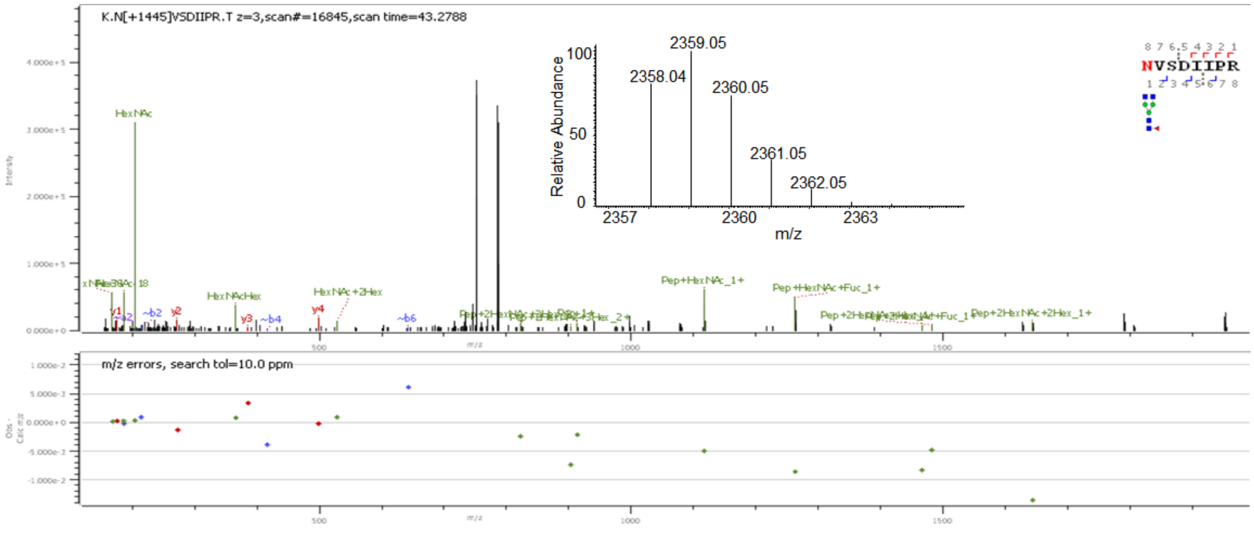
K.N[+1445]VSDIIPR.T z=3,scan#=16845,scan time=43.2788
Relative Abundance
m/z
2359.05
2358.04
2360.05
2361.05
2362.05
2357
2360
2363
NVSDIIPR
HexNAc
HexNAcHex
HexNAc+2Hex
Pep+HexNAc_1+
Pep+HexNAc+Fuc_1+
Pep+2HexNAc+2Hex_1+
m/z errors, search tol=10.0 ppm

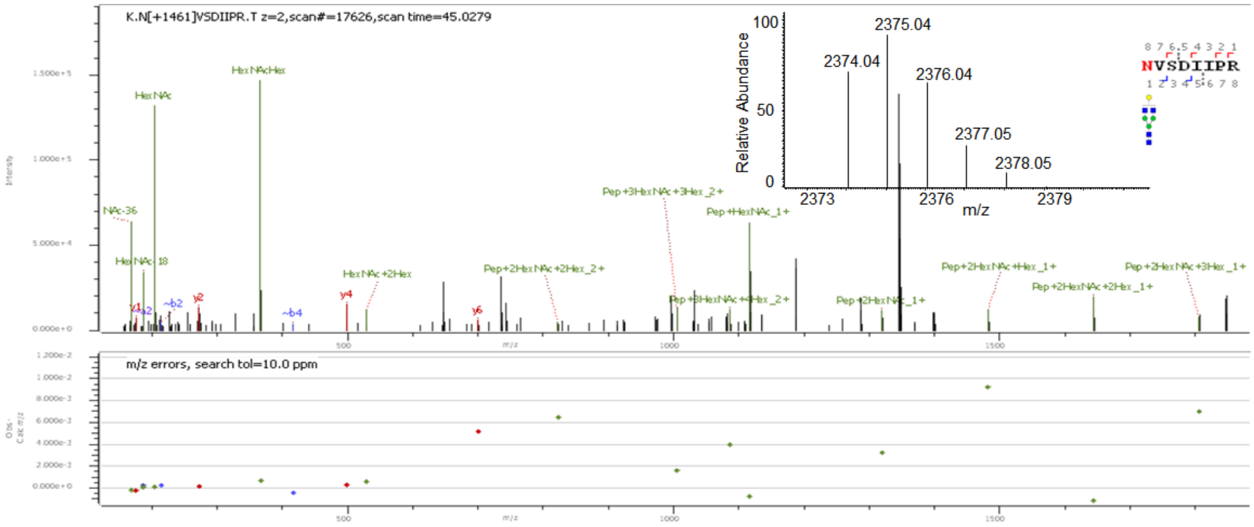
K.N[+1461]VSDIIPR.T z=2,scan#=17626,scan time=45.0279
Relative Abundance
m/z
2375.04
2374.04
2376.04
2377.05
2378.05
2373
2376
2379
NVSDIIPR
HexNAcHex
HexNAc
HexNAc+2Hex
Pep+2HexNAc+2Hex_2+
Pep+3HexNAc+3Hex_2+
Pep+HexNAc_1+
Pep+2HexNAc+Hex_1+
Pep+2HexNAc+3Hex_1+
Pep+2HexNAc+2Hex_1+
Pep+2HexNAc_1+
m/z errors, search tol=10.0 ppm

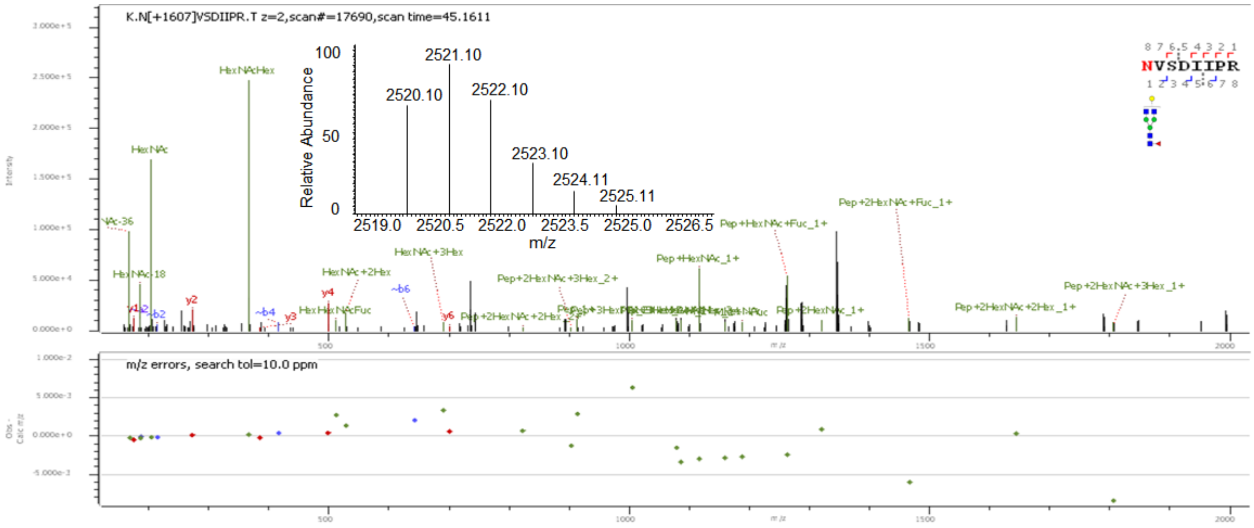
K.N[+1607]VSDIIPR.T z=2,scan#=17690,scan time=45.1611
Relative Abundance
m/z
2521.10
2520.10
2522.10
2523.10
2524.11
2525.11
2519.0
2520.5
2522.0
2523.5
2525.0
2526.5
NVSDIIPR
HexNAcHex
HexNAc
HexNAc+2Hex
HexNAc+3Hex
HexNAcFuc
Pep+HexNAc+Fuc_1+
Pep+2HexNAc+Fuc_1+
Pep+HexNAc_1+
Pep+2HexNAc+3Hex_2+
Pep+2HexNAc+3Hex_1+
Pep+2HexNAc+2Hex_1+
Pep+2HexNAc_1+
m/z errors, search tol=10.0 ppm

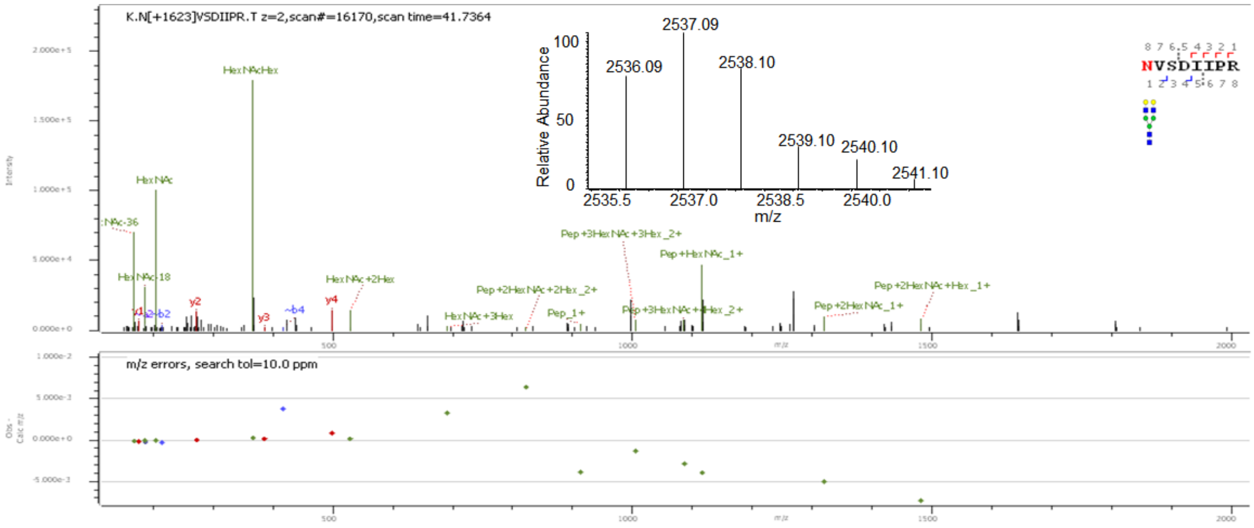
K.N[+1623]VSDIIPR.T z=2,scan#=16170,scan time=41.7364
2537.09
2536.09
2538.10
2539.10
2540.10
2541.10
Relative Abundance
m/z
NVSDIIPR
HexNAcHex
HexNAc
Pep+HexNAc_1+
Pep+2HexNAc+Hex_1+
m/z errors, search tol=10.0 ppm

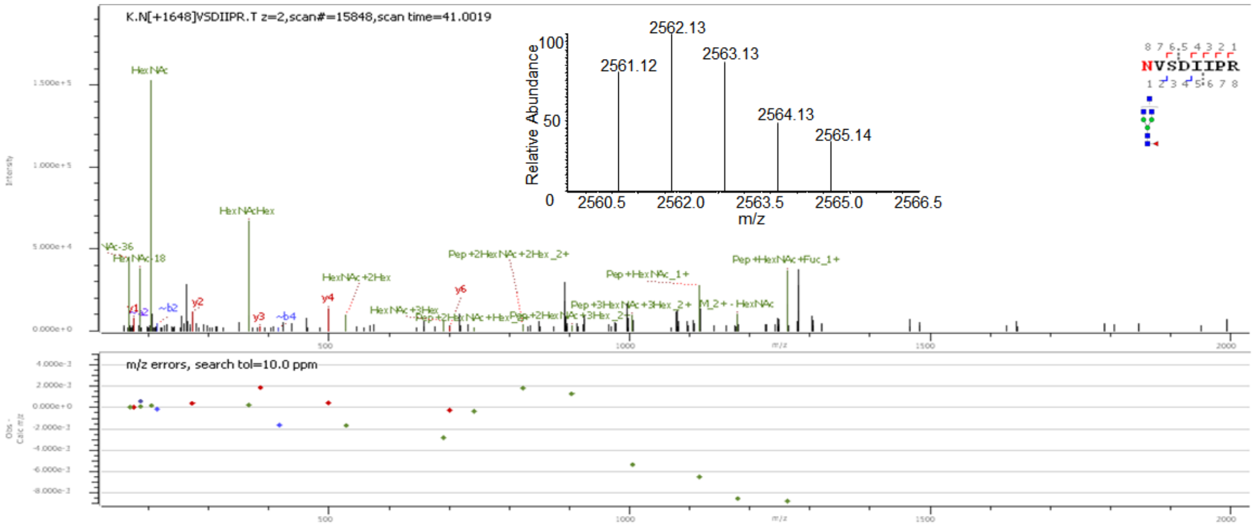
K.N[+1648]VSDIIPR.T z=2,scan#=15848,scan time=41.0019
2562.13
2561.12
2563.13
2564.13
2565.14
Relative Abundance
m/z
NVSDIIPR
HexNAc
HexNAcHex
Pep+HexNAc+Fuc_1+
m/z errors, search tol=10.0 ppm

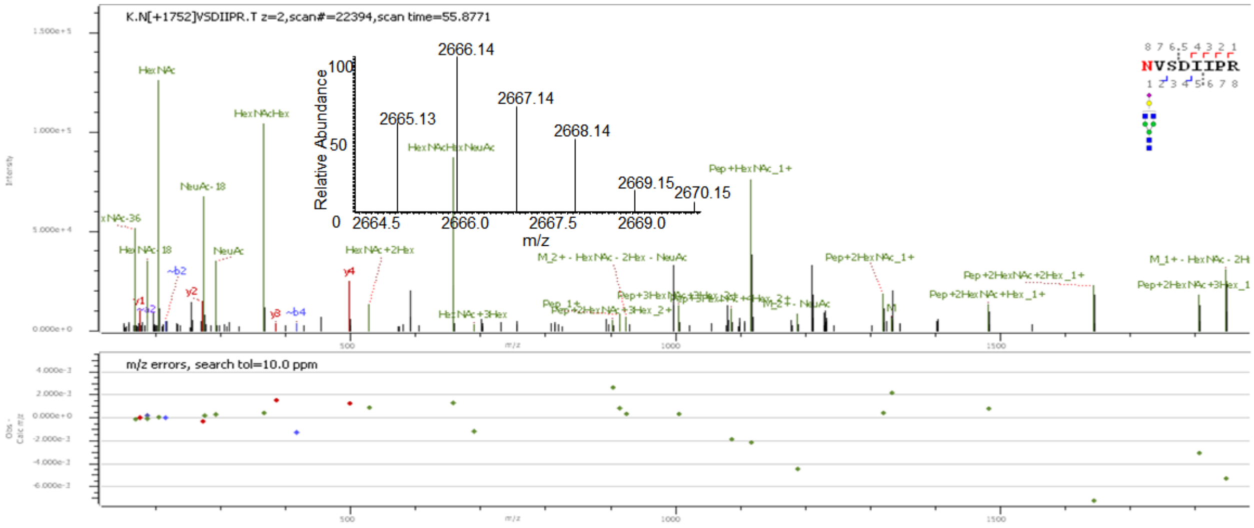
K.N[+1752]VSDIIPR.T z=2,scan#=22394,scan time=55.8771
2666.14
2665.13
2667.14
2668.14
2669.15
2670.15
Relative Abundance
m/z
NVSDIIPR
HexNAc
HexNAcHex
NeuAc-18
NeuAc
HexNAcHexNeuAc
Pep+HexNAc_1+
Pep+2HexNAc_1+
m/z errors, search tol=10.0 ppm

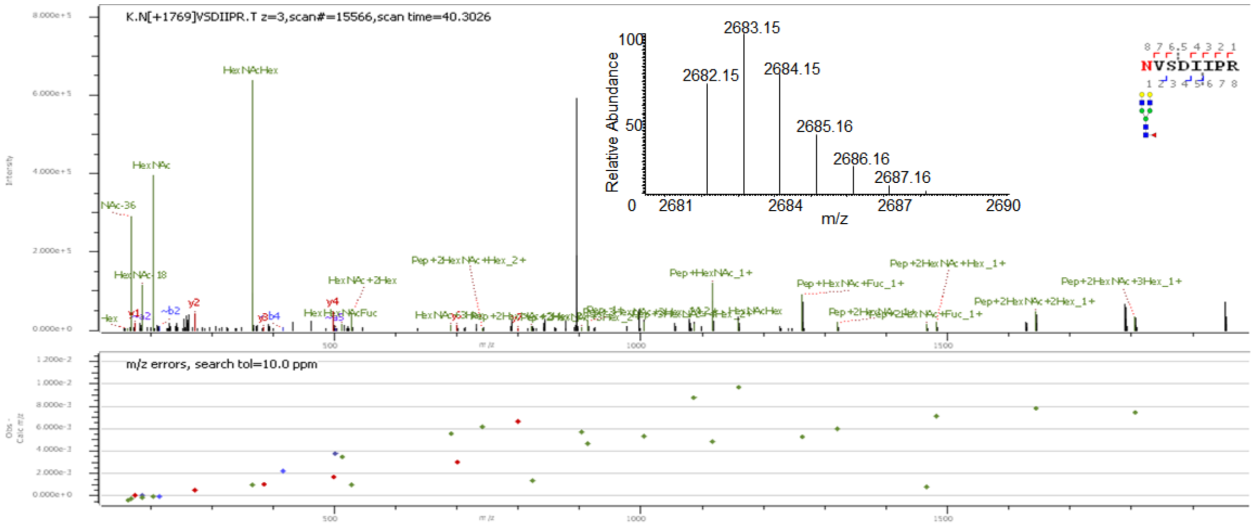
K.N[+1769]VSDIIPR.T z=3,scan#=15566,scan time=40.3026
Relative Abundance
m/z
2683.15
2682.15
2684.15
2685.16
2686.16
2687.16
HexNAcHex
HexNAc
Pep+2HexNAc+Hex_2+
Pep+HexNAc_1+
Pep+2HexNAc+Hex_1+
Pep+HexNAc+Fuc_1+
Pep+2HexNAc+3Hex_1+
Pep+2HexNAc+2Hex_1+
HexNAc+2Hex
NVSDIIPR
m/z errors, search tol=10.0 ppm

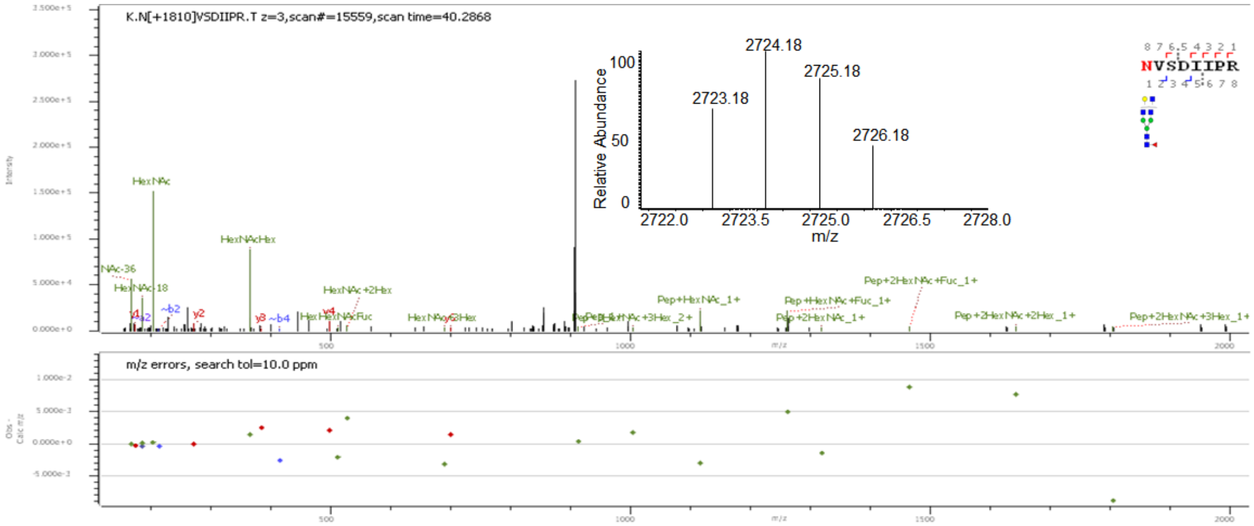
K.N[+1810]VSDIIPR.T z=3,scan#=15559,scan time=40.2868
Relative Abundance
m/z
2724.18
2723.18
2725.18
2726.18
HexNAc
HexNAcHex
HexNAc+2Hex
Pep+HexNAc_1+
Pep+HexNAc+Fuc_1+
Pep+2HexNAc_1+
Pep+2HexNAc+Fuc_1+
Pep+2HexNAc+2Hex_1+
Pep+2HexNAc+3Hex_1+
NVSDIIPR
m/z errors, search tol=10.0 ppm

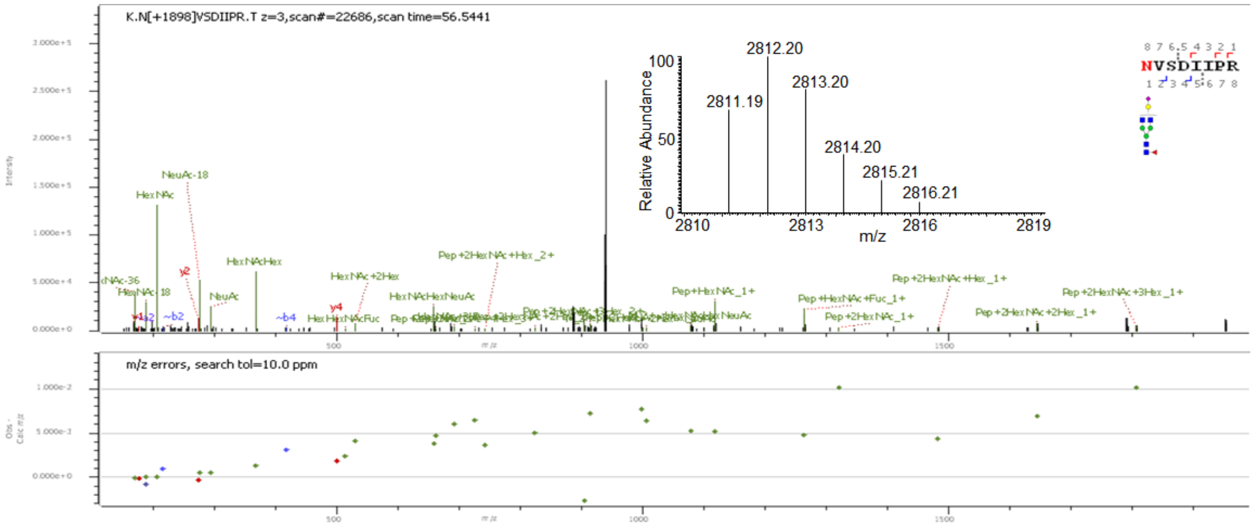
K.N[+1898]VSDIIPR.T z=3,scan#=22686,scan time=56.5441
Relative Abundance
m/z
2812.20
2811.19
2813.20
2814.20
2815.21
2816.21
NeuAc-18
HexNAc
HexNAcHex
NeuAc
HexNAc+2Hex
HexNAcHexNeuAc
Pep+2HexNAc+Hex_2+
Pep+HexNAc_1+
Pep+HexNAc+Fuc_1+
Pep+2HexNAc_1+
Pep+2HexNAc+Hex_1+
Pep+2HexNAc+2Hex_1+
Pep+2HexNAc+3Hex_1+
NVSDIIPR
m/z errors, search tol=10.0 ppm

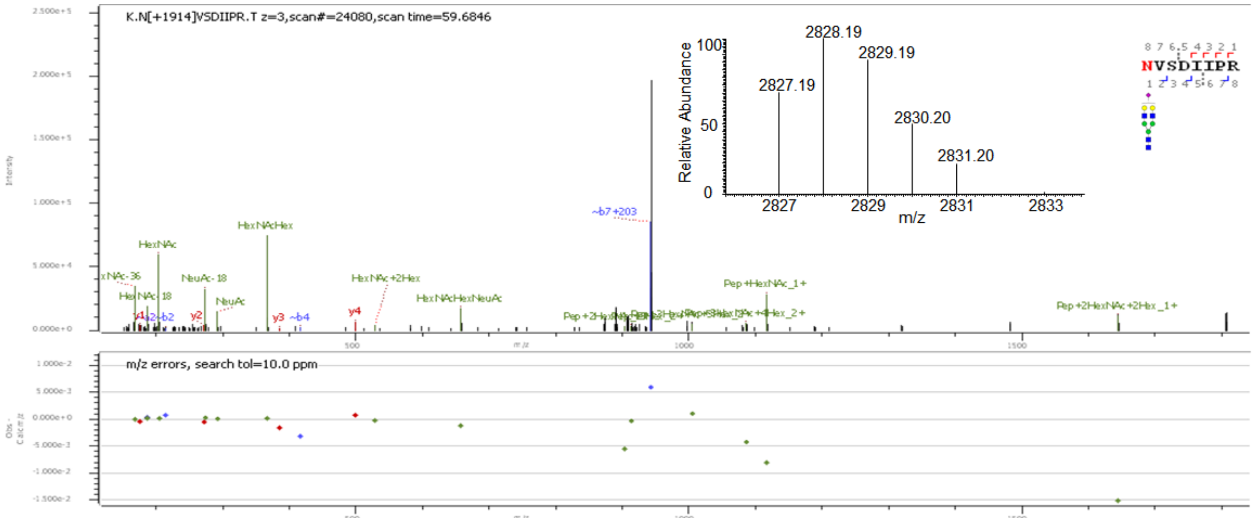
K.N[+1914]VSDIIPR.T z=3,scan#=24080,scan time=59.6846
NVSDIIPR
HexNAc
HexNAcHex
NeuAc-18
NeuAc
HexNAc+2Hex
HexNAcHexNeuAc
~b7+203
Pep+HexNAc_1+
Pep+2HexNAc+2Hex_1+
Relative Abundance
2828.19
2829.19
2827.19
2830.20
2831.20
m/z
m/z errors, search tol=10.0 ppm

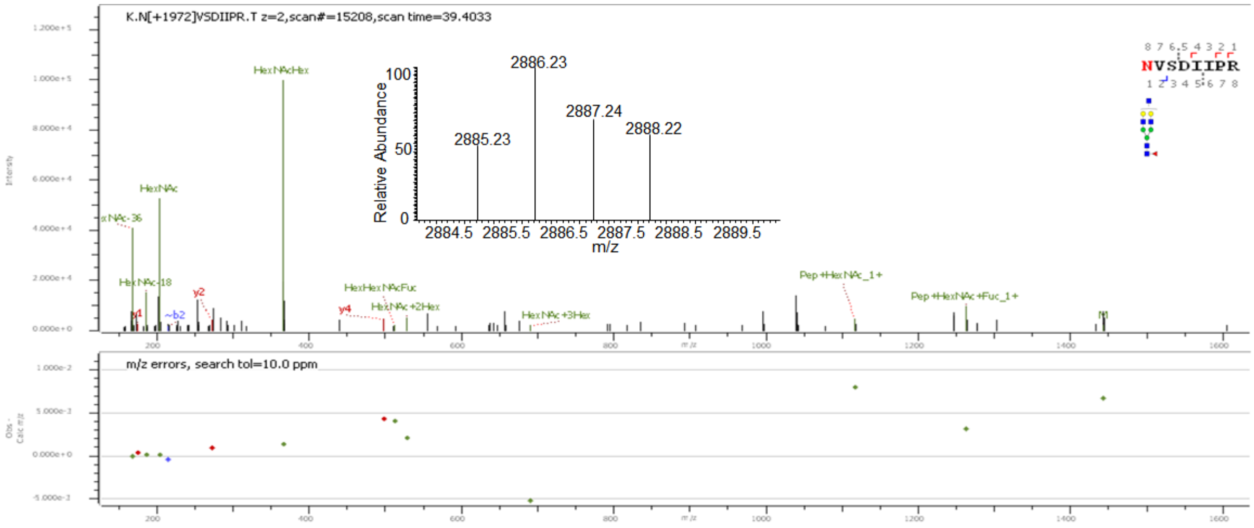
K.N[+1972]VSDIIPR.T z=2,scan#=15208,scan time=39.4033
NVSDIIPR
HexNAcHex
HexNAc
HexHexNAcFuc
HexNAc+2Hex
HexNAc+3Hex
Pep+HexNAc_1+
Pep+HexNAc+Fuc_1+
Relative Abundance
2886.23
2887.24
2888.22
2885.23
m/z
m/z errors, search tol=10.0 ppm

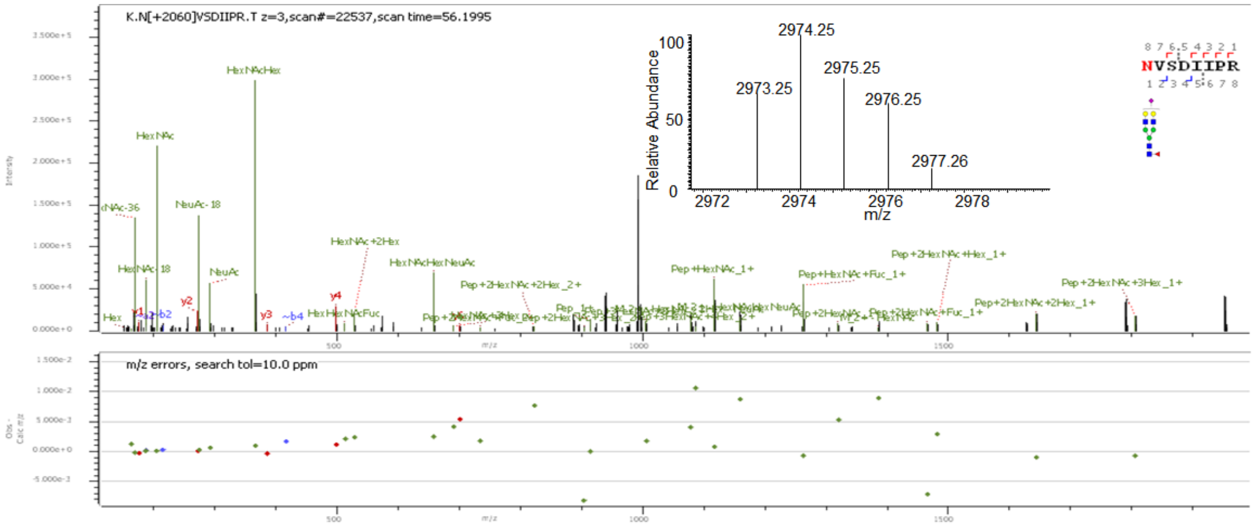
K.N[+2060]VSDIIPR.T z=3,scan#=22537,scan time=56.1995
NVSDIIPR
HexNAcHex
HexNAc
NeuAc-18
NeuAc
HexNAc+2Hex
HexNAcHexNeuAc
Pep+2HexNAc+2Hex_2+
Pep+HexNAc_1+
Pep+HexNAc+Fuc_1+
Pep+2HexNAc+Hex_1+
Pep+2HexNAc+3Hex_1+
Pep+2HexNAc+2Hex_1+
Relative Abundance
2974.25
2975.25
2973.25
2976.25
2977.26
m/z
m/z errors, search tol=10.0 ppm

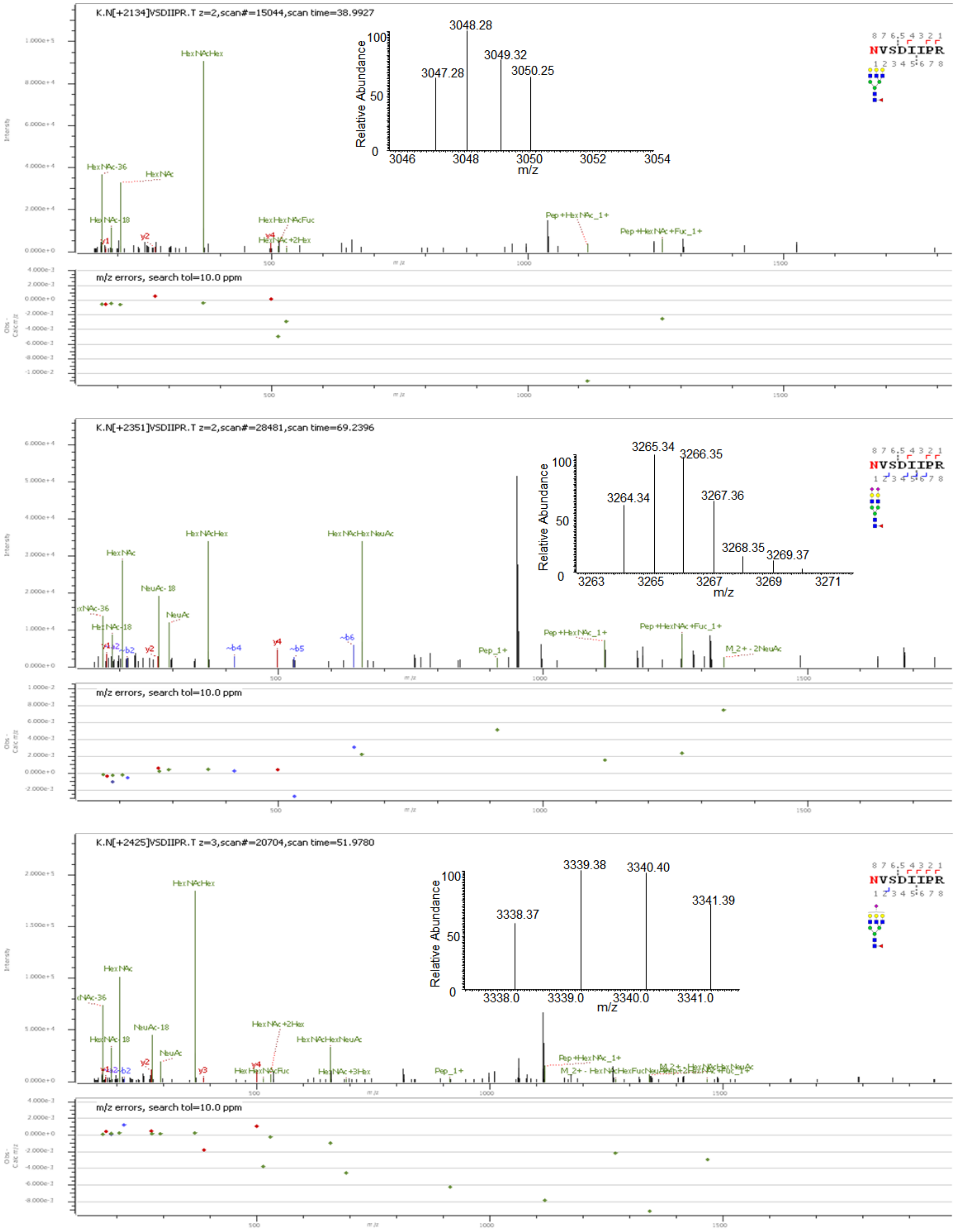


**Supp. Figure S7:** Deconvoluted precursor ion isotopic patterns (insets) and annotated HCD MS/MS spectra of intact N-glycopeptides with assigned N-glycans at N690.

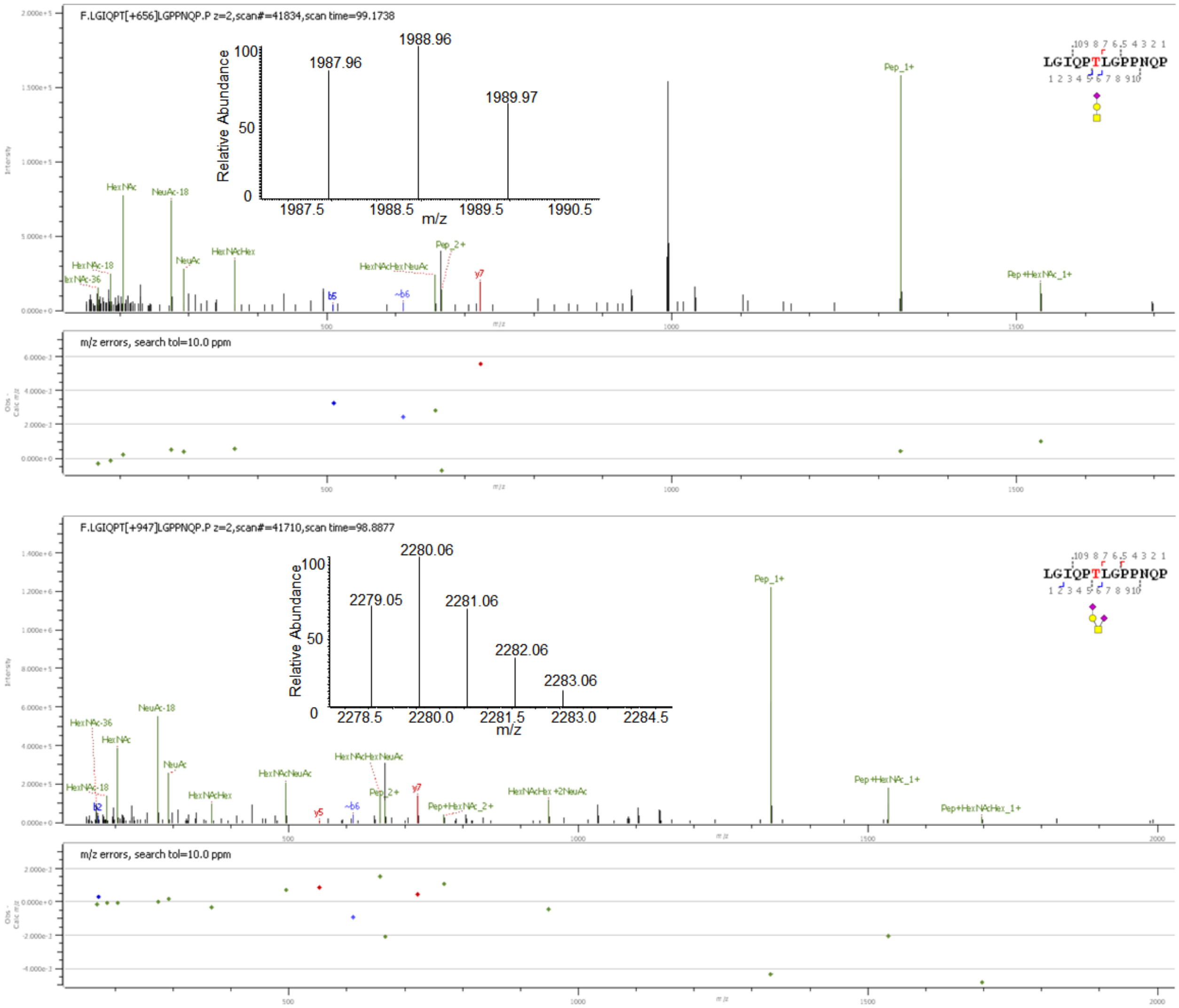


**Supp. Figure S8:** Deconvoluted precursor ion isotopic patterns (insets) and annotated HCD MS/MS spectra of intact O-glycopeptides with assigned O-glycans at T730.

Glycomics Analysis:

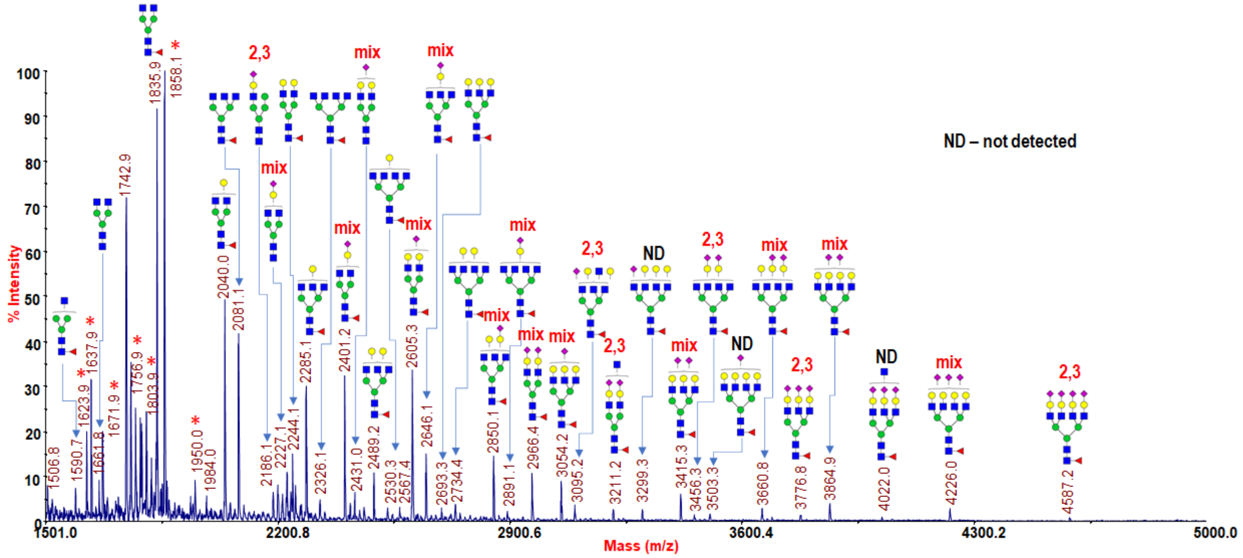


**Supp. Figure S9:** MALDI-MS spectrum showing the permethylated N-glycans released from hACE2 by PNGase F along with sialic acid linkages (determined by ESI-MS$^n$).

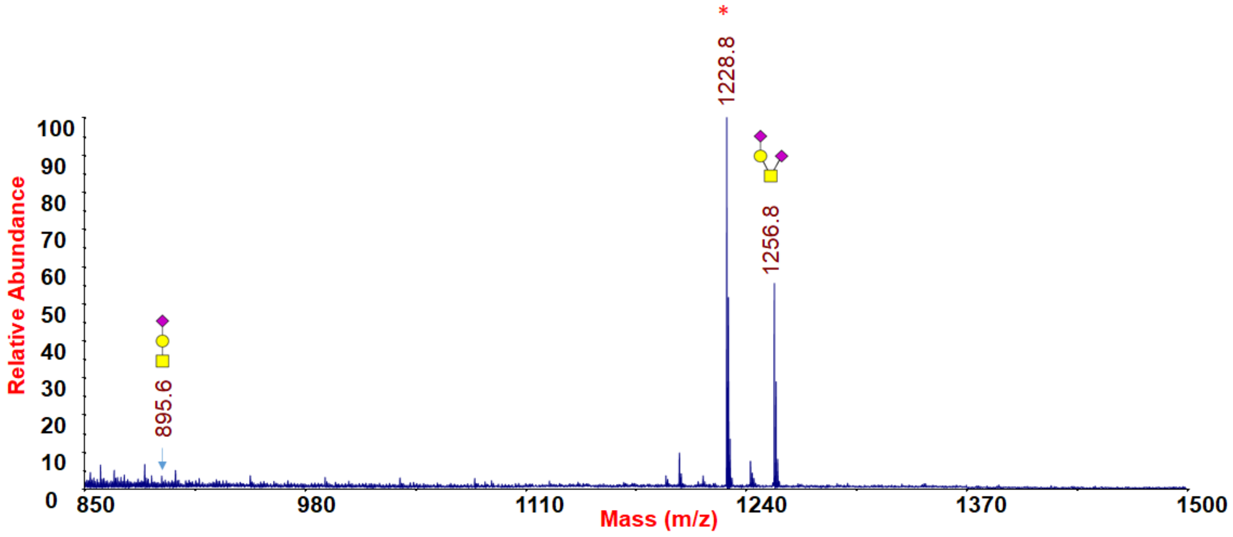


**Supp. Figure S10:** MALDI-MS spectrum showing the permethylated O-glycans released from hACE2 by β-elimination.

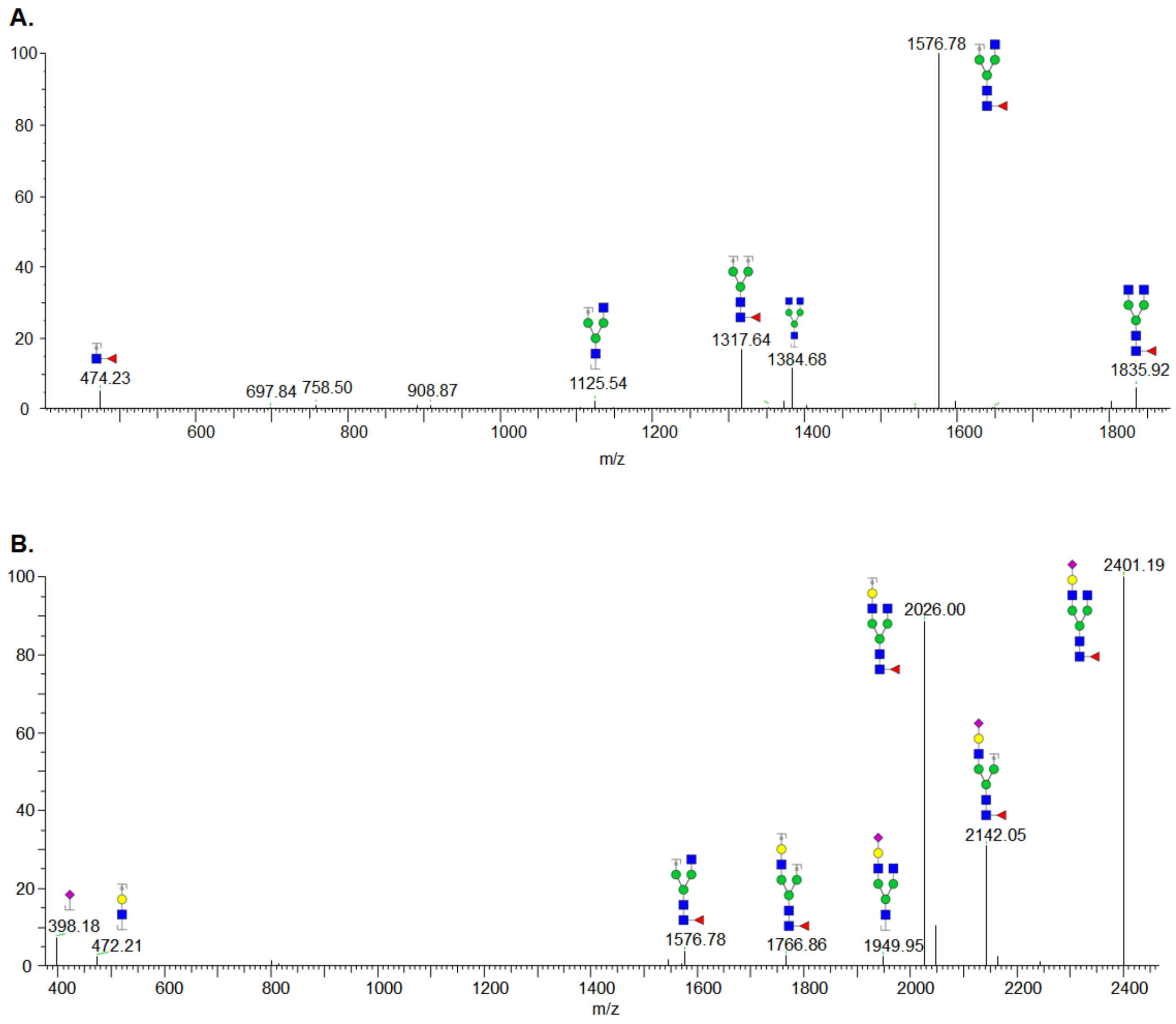


**Supp. Figure S11:** Representative CID ESI-MS/MS spectrum showing the fragments of permethylated N-glycans released from hACE2; **A.** N-glycan $GlcNAc_2Fuc_1Man_3GlcNAc_2$; **B.** N-glycan $GlcNAc_2Fuc_1Man_3GlcNAc_2Gal_1NeuAc_1$.

.
