## Supplementary material for "Comprehensive characterization of N- and O- glycosylation of SARS-CoV-2 human receptor angiotensin converting enzyme 2": Supp_Table 1

| **Position** | **Sequence** | **Modification** | **m/z** | **Charge** | **Ppm err.** | **RT** | **Delta Mod. Score** |
| --- | --- | --- | --- | --- | --- | --- | --- |
| N53 [N36] | NYNTNITEE | unmodified | ND | ND | ND | ND | ND |
|  | NYNTNITEE | HexNAc(4)(Hex(3)Fuc(1) | 1271.5092 | 2 | 1.06 | 26.3 | 198.5 |
|  | NYNTNITEE | HexNAc(4)Hex(4)Fuc(1) | 859.0126 | 3 | 1.78 | 12.3 | 46.4 |
|  | NYNTNITEE | HexNAc(5)Hex(3)Fuc(1) | 915.7014 | 3 | 0.72 | 25.8 | 111.8 |
|  | NYNTNITEE | HexNAc(5)Hex(4)Fuc(1) | 969.7180 | 3 | -0.36 | 25.7 | 59.8 |
|  | NYNTNITEE | HexNAc(6)Hex(3)Fuc(1) | 983.3944 | 3 | 0.48 | 25.6 | 111.6 |
|  | NYNTNITEE | HexNAc(6)Hex(4)Fuc(1) | 1037.4110 | 3 | -0.46 | 26.2 | 62.8 |
|  | NYNTNITEE | HexNAc(4)Hex(4)Fuc(1)NeuAc(1) | 999.0576 | 3 | 0.57 | 35.0 | 84.9 |
|  | NYNTNITEE | HexNAc(4)Hex(5)Fuc(1)NeuAc(1) | 1053.0746 | 3 | -0.09 | 35.1 | 31.6 |
|  | NYNTNITEE | HexNAc(5)Hex(4)Fuc(1)NeuAc(1) | 1066.7505 | 3 | 0.30 | 35.9 | 146.8 |
|  | NYNTNITEE | HexNAc(5)Hex(5)Fuc(1)NeuAc(1) | 1120.7659 | 3 | -1.70 | 33.8 | 118.1 |
|  | NYNTNITEE | HexNAc(6)Hex(4)Fuc(1)NeuAc(1) | 1134.4444 | 3 | 0.92 | 34.4 | 142.8 |
|  | NYNTNITEE | HexNAc(6)Hex(5)Fuc(1)NeuAc(1) | 1188.4605 | 3 | -0.36 | 35.8 | 5.3 |
|  | NYNTNITEE | HexNAc(6)Hex(6)Fuc(1)NeuAc(1) | 1242.4782 | 3 | -0.35 | 33.58 | manually |
| N90 [N73] | AQMYPLQEIQNL | unmodified | ND | ND | ND | ND | ND |
|  | AQMYPLQEIQNL | HexNAc(3)Hex(3) | 1272.0636 | 2 | -0.61 | 95.3 | 135.4 |
|  | AQMYPLQEIQNL | HexNAc(4)Hex(3) | 1373.6058 | 2 | 1.21 | 94.2 | 187.9 |
|  | AQMYPLQEIQNL | HexNAc(4)Hex(3)Fuc(1) | 964.7577 | 3 | -0.08 | 93.7 | 185.3 |
|  | AQMYPLQEIQNL | HexNAc(4)Hex(4) | 970.0891 | 3 | -0.30 | 94.0 | 173.6 |
|  | AQMYPLQEIQNL | HexNAc(4)Hex(4)Fuc(1) | 1018.7751 | 3 | -0.34 | 93.4 | 247.6 |
|  | AQMYPLQEIQNL | HexNAc(4)Hex(4)NeuAc(1) | 1067.1214 | 3 | 0.12 | 103.1 | 394.8 |
|  | AQMYPLQEIQNL | HexNAc(4)Hex(5)Fuc(1) | 1072.7937 | 3 | 0.64 | 92.8 | 316.8 |
|  | AQMYPLQEIQNL | HexNAc(4)Hex(4)Fuc(1)NeuAc(1) | 1115.8063 | 3 | -0.81 | 103.2 | 195.1 |
|  | AQMYPLQEIQNL | HexNAc(4)Hex(5)NeuAc(1) | 1121.1389 | 3 | 0.09 | 102.7 | 175.4 |
|  | AQMYPLQEIQNL | HexNAc(4)Hex(5)Fuc(1)NeuAc(1) | 1169.8256 | 3 | 0.69 | 102.4 | 132.2 |
| N103 [N86] | ALQQNGSSVLSEDK | unmodified | ND | ND | ND | ND | ND |
|  | ALQQNGSSVLSEDK | HexNAc(3)Hex(3)Fuc(1) | 1129.4997 | 2 | 0.22 | 38.2 | 338.3 |
|  | ALQQNGSSVLSEDK | HexNAc(4)Hex(3)Fuc(1) | 974.0940 | 3 | -0.05 | 38.1 | 398.1 |
|  | ALQQNGSSVLSEDK | HexNAc(4)Hex(4)Fuc(1) | 1028.1118 | 3 | 0.22 | 29.8 | 362.0 |
|  | ALQQNGSSVLSEDK | HexNAc(4)Hex(5)Fuc(1) | 1082.1287 | 3 | -0.50 | 29.6 | 323.6 |
|  | ALQQNGSSVLSEDK | HexNAc(4)Hex(5)Fuc(1)NeuAc(1) | 1179.1591 | 3 | -1.63 | 39.1 | 113.8 |
| N322 [N305] | VSVGLPNMTQGFW | unmodified | ND | ND | ND | ND | ND |
|  | VSVGLPNMTQGFW | HexNAc(4)Hex(3)Fuc(1) | 960.7509 | 3 | 0.14 | 101.8 | 257.5 |
|  | VSVGLPNMTQGFW | HexNAc(5)Hex(3)Fuc(1) | 1542.1617 | 2 | -0.33 | 101.6 | 124.0 |
|  | VSVGLPNMTQGFW | HexNAc(5)Hex(4)Fuc(1) | 1082.4632 | 3 | 1.56 | 101.2 | 253.4 |
|  | VSVGLPNMTQGFW | HexNAc(6)Hex(3)Fuc(1) | 1096.1358 | 3 | -1.12 | 101.5 | 129.9 |
|  | VSVGLPNMTQGFW | HexNAc(5)Hex(5)Fuc(1) | 1136.4789 | 3 | -0.16 | 100.9 | 83.5 |
|  | VSVGLPNMTQGFW | HexNAc(6)Hex(4)Fuc(1) | 1150.1534 | 3 | -1.07 | 101.2 | 196.9 |
|  | VSVGLPNMTQGFW | HexNAc(5)Hex(4)Fuc(1) NeuAc(1) | 1179.4931 | 3 | -0.16 | 107.3 | 253.6 |
|  | VSVGLPNMTQGFW | HexNAc(5)Hex(5)Fuc(1) NeuAc(1) | 1233.5112 | 3 | 0.22 | 106.8 | 120.9 |
|  | VSVGLPNMTQGFW | HexNAc(5)Hex(6)Fuc(1) NeuAc(1) | 1287.5273 | 3 | -0.95 | 106.7 | 211.9 |
|  | VSVGLPNMTQGFW | HexNAc(5)Hex(6)Fuc(1) NeuAc(2) | 1384.5605 | 3 | 0.13 | 114.2 | 88.0 |
|  | VSVGLPNMTQGFW | HexNAc(6)Hex(5)Fuc(1) NeuAc(2) | 1398.2347 | 3 | -0.82 | 114.0 | 76.6 |
|  | VSVGLPNMTQGFW | HexNAc(6)Hex(6)Fuc(1) NeuAc(2) | 1452.2337 | 3 | 0.20 | 113.7 | 72.6 |
|  | VSVGLPNMTQGFW | HexNAc(6)Hex(7)Fuc(1) NeuAc(2) | 1506.2710 | 3 | 0.74 | 113.1 | 60.9 |
| N432 [N415] | LSPDFQEDNETEINF | unmodified | 899.3950 | 2 | 0.67 | 95.62 | manually |
|  | LSPDFQEDNETEINF | HexNAc(4)Hex(3)Fuc(1) | 1081.4455 | 3 | 2.04 | 87.3 | 164.0 |
|  | LSPDFQEDNETEINF | HexNAc(4)Hex(4)Fuc(1) | 1135.4601 | 3 | -0.66 | 85.6 | 208.1 |
|  | LSPDFQEDNETEINF | HexNAc(5)Hex(3)Fuc(1) | 1149.1361 | 3 | -0.21 | 86.1 | 196.1 |
|  | LSPDFQEDNETEINF | HexNAc(4)Hex(5)Fuc(1) | 1189.4782 | 3 | -0.22 | 84.8 | 56.2 |
|  | LSPDFQEDNETEINF | HexNAc(5)Hex(4)Fuc(1) | 1203.1561 | 3 | 1.73 | 84.9 | 120.7 |
|  | LSPDFQEDNETEINF | HexNAc(4)Hex(4)Fuc(1)NeuAc(1) | 1232.4916 | 3 | -0.84 | 98.5 | 65.1 |
|  | LSPDFQEDNETEINF | HexNAc(5)Hex(5)Fuc(1) | 1257.1716 | 3 | -0.02 | 84.8 | 80.1 |
|  | LSPDFQEDNETEINF | HexNAc(4)Hex(5)Fuc(1)NeuAc(1) | 1286.5107 | 3 | 0.31 | 97.7 | 19.0 |
|  | SIGLLSPDFQEDNETEINF | HexNAc(5)Hex(4)Fuc(1)NeuAc(1) | 1423.5930 | 3 | 0.40 | 49.0 |  |
|  | LSPDFQEDNETEINF | HexNAc(5)Hex(6)Fuc(1) | 1408.2253 | 3 | 3.03 | 97.0 | 42.4 |
|  | SIGLLSPDFQEDNETEINF | HexNAc(5)Hex(5)Fuc(1)NeuAc(1) | 1477.6097 | 3 | -0.63 | 30.2 |  |
|  | LSPDFQEDNETEINF | HexNAc(4)Hex(5)Fuc(1)NeuAc(2) | 1383.8732 | 3 | 2.23 | 106.5 | 48.2 |
|  | LSPDFQEDNETEINF | HexNAc(5)Hex(6)Fuc(1)NeuAc(1) | 1408.2253 | 3 | 3.03 | 97.0 | 42.4 |
|  | LSPDFQEDNETEINF | HexNAc(5)Hex(6)Fuc(1)NeuAc(2) | 1628.6603 | 3 | 0.20 | 8.2 |  |
| N546 [529] | cDISNSTEAGQK | unmodified | ND | ND | ND | ND | ND |
|  | cDISNSTEAGQK | HexNAc(2)Hex(5) | 1263.5007 | 2 | 0.92 | 13.3 | 230.1 |
|  | cDISNSTEAGQK | HexNAc(3)Hex(3)Fuc(1) | 1276.0153 | 2 | 0.18 | 14.5 | 492.7 |
|  | cDISNSTEAGQK | HexNAc(3)Hex(4)Fuc(1) | 1357.0414 | 2 | -0.22 | 14.4 | 316.7 |
|  | cDISNSTEAGQK | HexNAc(4)Hex(3)Fuc(1) | 1377.5549 | 2 | -0.11 | 14.9 | 517.3 |
|  | cDISNSTEAGQK | HexNAc(4)Hex(4)Fuc(1) | 972.7243 | 3 | 0.99 | 14.7 | 437.8 |
|  | cDISNSTEAGQK | HexNAc(5)Hex(3)Fuc(1) | 9863997 | 3 | 0.88 | 14.3 | 391.1 |
|  | cDISNSTEAGQK | HexNAc(4)Hex(5)Fuc(1) | 1539.6082 | 2 | 0.24 | 14.7 | 196.5 |
|  | cDISNSTEAGQK | HexNAc(5)Hex(4)Fuc(1) | 1560.1197 | 2 | -0.92 | 14.0 | 167.6 |
|  | cDISNSTEAGQK | HexNAc(4)Hex(4)Fuc(1)NeuAc(1) | 1604.1302 | 2 | 0.65 | 21.2 | 219.9 |
| N690 [673] | NVSDIIPR | unmodified | 913.5099 | 1 | -0.33 | 46.63 |  |
|  | NVSDIIPR | HexNAc(3)Hex(3) | 1004.9563 | 2 | -0.70 | 43.9 | 186.4 |
|  | NVSDIIPR | HexNAc(2)Hex(5) | 1065.4709 | 2 | 0.68 | 41.4 | 254.1 |
|  | NVSDIIPR | HexNAc(3)Hex(3)Fuc(1) | 1077.9856 | 2 | -0.30 | 41.9 | 411.2 |
|  | NVSDIIPR | HexNAc(3)Hex(4) | 1085.9837 | 2 | 0.25 | 41.6 | 308.0 |
|  | NVSDIIPR | HexNAc(4)Hex(3) | 1106.4961 | 2 | -0.50 | 42.7 | 405.4 |
|  | NVSDIIPR | HexNAc(2)Hex(6) | 1146.4963 | 2 | -0.26 | 40.7 | 282.5 |
|  | NVSDIIPR | HexNAc(3)Hex(4)Fuc(1) | 1159.0138 | 2 | 1.27 | 41.5 | 301.9 |
|  | NVSDIIPR | HexNAc(4)Hex(3)Fuc(1) | 1179.5251 | 2 | -0.49 | 42.0 | 397.7 |
|  | NVSDIIPR | HexNAc(4)Hex(5) | 1268.5498 | 2 | 0.26 | 41.1 | 296.3 |
|  | NVSDIIPR | HexNAc(4)Hex(4)Fuc(1) | 1260.5519 | 2 | -0.11 | 41.0 | 336.3 |
|  | NVSDIIPR | HexNAc(4)Hex(5) | 1268.5510 | 2 | 1.19 | 43.7 | 176.3 |
|  | NVSDIIPR | HexNAc(5)Hex(3)Fuc(1) | 1281.0662 | 2 | 0.68 | 40.8 | 216.8 |
|  | NVSDIIPR | HexNAc(4)Hex(4)NeuAc(1) | 1333.0705 | 2 | -0.25 | 55.1 | 233.5 |
|  | NVSDIIPR | HexNAc(4)Hex(5)Fuc(1) | 1341.5794 | 2 | 0.69 | 41.3 | 343.6 |
|  | NVSDIIPR | HexNAc(5)Hex(4)Fuc(1) | 1362.0923 | 2 | 0.43 | 40.6 | 229.6 |
|  | NVSDIIPR | HexNAc(4)Hex(4)Fuc(1)NeuAc(1) | 937.7362 | 3 | 0.61 | 54.1 | 251.8 |
|  | NVSDIIPR | HexNAc(4)Hex(5) NeuAc(1) | 943.0675 | 3 | 0.25 | 54.2 | 304.9 |
|  | NVSDIIPR | HexNAc(5)Hex(5)Fuc(1) | 1443.1172 | 2 | -0.64 | 39.3 | 180.2 |
|  | NVSDIIPR | HexNAc(4)Hex(5)Fuc(1)NeuAc(1) | 1487.1266 | 2 | 0.31 | 53.3 | 283.7 |
|  | NVSDIIPR | HexNAc(5)Hex(6)Fuc(1) | 1524.1457 | 2 | 0.73 | 39.0 | 106.0 |
|  | NVSDIIPR | HexNAc(4)Hex(5)Fuc(1)NeuAc(2) | 1632.6736 | 2 | -0.16 | 69.2 | 260.0 |
|  | NVSDIIPR | HexNAc(5)Hex(6)Fuc(1)NeuAc(1) | 1113.4645 | 3 | 0.51 | 52.1 | 206.1 |
| T730 [713] | LGIQPTLGPPNQP | Unmodified | ND | ND | ND | ND | ND |
|  | LGIQPTLGPPNQP | HexNAc(1)Hex(1)NeuAc(1) | 994.4840 | 2 | 0.69 | 99.2 | 112.4 |
|  | LGIQPTLGPPNQP | HexNAc(1)Hex(1)NeuAc(2) | 1027.4757 | 2 | 0.34 | 88.2 | 224.6 |

ND = Not Detected

manually = indicates a glycopeptide that was manually observed.

Followed amino acid numbering from Uniprot.org for consistency. Actual amino acid locations are indicated in square brackets.

Site N53 data are from following LC-MS/MS .raw and Byonic search files

- “hACE2_GluC_ChymDig_InSoln_180min_HCDpdCID.raw”
- “hACE2_GluC_ChymDig_InSoln_180min_HCDpdCID_xtract.raw” [deconvoluted spectra]
- “hACE2_GluC_Chy_180min_Byonic”
- “hACE2_GluC_Chym_180min_FullACE2_TargetedN53glycans”

Sites N90, N103, N322, N432, N546, N690 and T730 data are from LC-MS/MS .raw files

- “hACE2_Trp_ChymDig_InSoln_180min_HCDpdCID.raw”
- “hACE2_Trp_ChymDig_InSoln_180min_HCDpdCID_xtract.raw” [deconvoluted spectra]
- “hACE2_Trp_Chy_180min_Byonic”
- “hACE2_DeGlycosylated_Trp_Chy_180min_Byonic”

Glycopost repository - https://glycopost.glycosmos.org/preview/15928494275f039184c7229; pin: 5771.
